# High-resolution structures of the actomyosin-V complex in three nucleotide states provide insights into the force generation mechanism

**DOI:** 10.1101/2021.09.07.459262

**Authors:** Sabrina Pospich, H. Lee Sweeney, Anne Houdusse, Stefan Raunser

**Affiliations:** Department of Structural Biochemistry, Max Planck Institute of Molecular Physiology, Dortmund, 44227, Germany; Department of Pharmacology and Therapeutics and the Myology Institute, University of Florida, College of Medicine, Gainesville, Florida 32610-0267, USA; Structural Motility, Institut Curie, Paris Université Sciences et Lettres, Sorbonne Université, CNRS UMR144, Paris, France

## Abstract

The molecular motor myosin undergoes a series of major structural transitions during its force-producing motor cycle. The underlying mechanism and its coupling to ATP hydrolysis and actin binding is only partially understood, mostly due to sparse structural data on actin-bound states of myosin. Here, we report 26 high-resolution cryo-EM structures of the actomyosin-V complex in the strong-ADP, rigor, and a previously unseen post-rigor transition state that binds the ATP analog AppNHp. The structures reveal a high flexibility of myosin in each state and provide valuable insights into the structural transitions of myosin-V upon ADP release and binding of AppNHp, as well as the actomyosin interface. In addition, they show how myosin is able to specifically alter the structure of F-actin. The unprecedented number of high-resolution structures of a single myosin finally enabled us to assemble a nearly complete structural model of the myosin-V motor cycle and describe the molecular principles of force production.

## Introduction

The molecular motor myosin is well known for its central role in muscle contraction (Hanson & Huxley, 1953; Szent-Györgyi, 2004). By using the actin cytoskeleton as tracks, myosin also powers cellular cargo transport processes and can serve as a molecular anchor and force sensor (Hartman, Finan, Sivaramakrishnan, & Spudich, 2011; Woolner & Bement, 2009). Due to its versatility, myosin is key to numerous essential cellular processes including cytokinesis, transcription, signal transduction, cell migration and adhesion, and endo- and exocytosis (Coluccio, 2020; Krendel & Mooseker, 2005). While this variety in functions is well reflected by the diversity of the myosin superfamily (Sellers, 2000), the ATP-dependent force generation mechanism as well as the architecture of the motor domain is shared by all myosins (Cope, Whisstock, Rayment, & Kendrick-Jones, 1996).

The myosin motor domain consists of four subdomains: the actin-binding upper and lower 50 kDa (U50 and L50) domains, which are separated by the central actin binding cleft, the N-terminal domain and the converter domain, containing the long α-helical extension known as the lever arm (Rayment, Rypniewski, et al., 1993b). The active site of myosin is located at the interface of the U50 domain and the N-terminal domain and is allosterically coupled to both the actin binding interface and the lever arm (Sweeney & Houdusse, 2010). This coupling ultimately enables the amplification of small rearrangements at the active site to large, force-producing conformational changes of the lever arm (Holmes, 1997; Rayment, Holden, et al., 1993a).

The ATP-driven mechanism of myosin force generation relies on several major structural transitions and is described in the myosin motor cycle (Huxley, 1958; Lymn & Taylor, 1971). Initially, hydrolysis of ATP places myosin in a conformation known as the pre-powerstroke (PPS) state. The mechano-chemical energy stored in this conformation is released by binding to filamentous actin (F-actin), which serves as an activator and initiates a cascade of allosteric structural changes (Reubold, Eschenburg, Becker, Kull, & Manstein, 2003). These changes eventually result in phosphate release —potentially via a phosphate release (P_i_R) state (Llinas et al., 2015)— and the major, force-producing lever arm swing known as the powerstroke. Subsequent release of ADP from myosin in the strong-ADP state gives rise to a second, smaller lever arm swing, leaving myosin strongly bound to F-actin in the rigor state (Mentes et al., 2018). Binding of ATP to the now unoccupied active site causes a transition to the post-rigor state and eventual detachment from F-actin (Kühner & Fischer, 2011). Finally, ATP hydrolysis triggers the repriming of the lever arm through the so-called recovery stroke, thus completing the myosin motor cycle.

Decades of biochemical studies have brought great insights into the diversity and kinetics of the myosin superfamily (Coluccio, 2020; Geeves, Fedorov, & Manstein, 2005). However, detailed structural information is ultimately required to understand the mechanism of force generation. Over the years, X-ray crystallography has revealed the structures of various myosins in the post-rigor state (Rayment, Rypniewski, et al., 1993b), the PPS state (C. A. Smith & Rayment, 1996), the rigor-like state (Coureux et al., 2003), a putative P_i_R state (Llinas et al., 2015), as well as the intermediate recovery stroke state (Blanc et al., 2018) (for a recent review of all available crystal structures see (Sweeney, Houdusse, & Robert-Paganin, 2020)). Due to the reluctance of F-actin to crystallize, actin-bound states of myosin are not accessible by X-ray crystallography. Instead, electron cryo microscopy (cryo-EM) has proven to be an optimal tool to study filamentous proteins (Pospich & Raunser, 2018) such as the actomyosin complex (Behrmann et al., 2012; von der Ecken, Heissler, Pathan-Chhatbar, Manstein, & Raunser, 2016). To date, the structure of the actomyosin rigor complex has been determined for a variety of myosins (Banerjee et al., 2017; Behrmann et al., 2012; Doran et al., 2020; Fujii & Namba, 2017; Gong et al., 2021; Gurel et al., 2017; Mentes et al., 2018; Risi et al., 2020; Robert-Paganin et al., 2021; Vahokoski et al., 2020; von der Ecken et al., 2016). States other than the nucleotide-free rigor state have proven more difficult to study, mainly due to lower binding affinities and short lifetimes. In fact, the only other state solved to date is the strong-ADP state; and only two (myosin-IB, myosin-XV) (Gong et al., 2021; Mentes et al., 2018) of four independent studies (myosin-Va, myosin-VI) (Gurel et al., 2017; Wulf et al., 2016) have achieved high resolution (< 4 Å). However, the actin-bound states of myosin, in particular weakly bound transition states for which no structure is yet available, are precisely those that are urgently needed to understand important properties of the myosin motor cycle, such as binding to and detachment from F-actin (recently reviewed in (Schröder, 2020)). In addition, high-resolution structures of other myosins in the rigor and especially strong-ADP state are required to identify conserved and specific features within the myosin superfamily. Finally, structures of all key states of the motor cycle need to be determined for a single myosin to allow the assembly of a reliable structural model, since the structures of different myosins vary considerably within the same state (Merino, Pospich, & Raunser, 2019).

High-duty ratio myosins, such as myosin-I, V and VI, are characterized by comparatively high binding affinities for F-actin and long lifetimes of actin-bound states (De La Cruz & Ostap, 2004; De La Cruz, Ostap, & Sweeney, 2001; De La Cruz, Wells, Rosenfeld, Ostap, & Sweeney, 1999; Laakso, Lewis, Shuman, & Ostap, 2008). Therefore, they are best suited to structurally study actin-bound states other than the rigor. Today, class V and VI myosins are probably the best-characterized unconventional myosins, both structurally and biochemically (Coluccio, 2020). Cryo-EM studies of actomyosin-V have further reported structures of the strong-ADP and rigor state (Wulf et al., 2016), as well as a potential pre-powerstroke transition state (Volkmann et al., 2005). However, due to the limited resolution of these structures, atomic details could not be modeled and the structural transition of actin-bound myosin-V during its motor cycle has consequently remained elusive. Interestingly, myosin-V was also shown to be sensitive to the nucleotide state of phalloidin-stabilized F-actin, preferring young ATP/ADP-P_i_-bound F-actin over aged ADP-bound F-actin (Zimmermann, Santos, Kovar, & Rock, 2015). The structural basis and implications of this preference have not yet been uncovered.

Here, we present high-resolution cryo-EM structures of the actomyosin-V complex in three nucleotide states. Specifically, we have solved the structure of myosin-V in the strong-ADP state (ADP), the rigor state (nucleotide free), and a previously unseen post-rigor transition state, which has the non-hydrolysable ATP analogue AppNHp bound to its active site. To investigate the structural effect the nucleotide state of F-actin has on myosin-V, we have also determined the structure of the rigor complex starting from young ADP-P_i_-bound F-actin, rather than from aged ADP-bound F-actin. In addition to these structures and their implications, we report a pronounced conformational heterogeneity of myosin-V in all our data sets and characterize it in detail based on 18 high-resolution subset structures. Finally, we present what is, to our knowledge, currently the most complete structural model of the myosin motor cycle and identify key elements of force production.

## Results and Discussion

### High-resolution cryo-EM structures of the actomyosin-V complex

With the aim to shed light on the structural transition of myosin along its motor cycle, we determined the structure of the actomyosin-V complex in three different nucleotide states using single particle cryo-EM. Specifically, we have decorated aged ADP-bound F-actin (rabbit skeletal α-actin) stabilized by phalloidin (PHD) (Lynen & Wieland, 1938) with myosin-Va —S1 fragment bound to one essential light chain, hereafter referred to as myosin-V. The complex was either prepared in absence of a nucleotide, or after brief incubation of myosin with Mg^2+^-ADP or Mg^2+^-AppNHp (see Methods for details). AppNHp, also known as AMPPNP, is an ATP analog that has been shown to be non-hydrolysable by myosin-V (Yengo, De La Cruz, Safer, Ostap, & Sweeney, 2002). It is coordinated similarly to ATP in crystal structures of myosin-II (Bauer, Holden, Thoden, Smith, & Rayment, 2000; Gulick, Bauer, Thoden, & Rayment, 1997) and has also been reported to lead to a mixture of a pre- and post-powerstroke conformations in myosin-V (Volkmann et al., 2005). These results suggest that AppNHp can potentially mimic both ATP and ADP-P_i_ and is thus well suited to capture short-lived actin-bound transition states, such as the weakly bound PPS and post-rigor states (Sweeney & Houdusse, 2010).

We collected cryo-EM data sets of the different samples (Table 1) and processed them using the helical processing pipeline implemented in the SPHIRE package (Moriya et al., 2017; Pospich et al., 2021; Stabrin, Schoenfeld, Wagner, Pospich, Gatsogiannis, & Raunser, 2020a), which applies helical restraints but no symmetry. For each data set, two all-particle density maps were reconstructed (Table 1—Supplementary Figure 1, see Methods for details). In this way, we achieved nominal resolutions of 3.0 Å/ 3.1 Å (ADP), 3.2 Å/ 3.3 Å (Rigor) and 2.9 Å/ 2.9Å (AppNHp), respectively (Table 1—Supplementary Figures 1-2, Table 1 —Supplementary Tables 1-3), allowing us to reliably model each state and analyze its molecular interactions.

**Table 1.** Data collection statistics of F-actin and actomyosin data sets. Aged PHD-stabilized F-actin (F-actin-PHD) was decorated with myosin-V in the rigor (no nucleotide), strong-ADP (bound to Mg^2+^-ADP) and post-rigor transition (PRT) state (bound to Mg^2+^-AppNHp). Young JASP-stabilized F-actin (F-actin-JASP) was imaged in absence and presence of myosin-V in the rigor state. Refinement and model building statistics can be found in Table 1— Supplementary Tables 1-3 and Figure 7—Table 1. See Table 1— Supplementary Figures 1 and 2 for an overview of the processing pipeline and the cryo-EM data, respectively.

| Microscopy | Aged F-actin-PHD |  |  |  |  | Young F-actin-JASP |  |
| --- | --- | --- | --- | --- | --- | --- | --- |
|  | ADP | Rigor | AppNHp 4 °C | AppNHp 25 °C | AppNHp<sup>a</sup> | Actin only | Rigor |
| Microscope | Titan Krios - Cs 2.7 mm |  | Titan Krios - Cs-corrected |  |  |  |  |
| Voltage [kV] | 300 |  |  |  |  |  |  |
| Camera | K2 - super resolution |  |  |  |  |  |  |
| Energy filter slit width [eV] | 20 |  |  |  |  |  |  |
| Pixel size [Å] | 1.06 |  | 1.10 |  |  |  |  |
| Frames per movie | 40 |  |  |  |  |  |  |
| Exposure time [s] | 15 |  |  |  |  |  |  |
| Total electron dose [e/Å<sup>2</sup>] | 82 | 79 | 81 | 81 | 81 | 80 | 80 |
| Final electron dose [e/Å<sup>2</sup>] | Dose weighted |  |  |  |  | Polished particles |  |
| Defocus range [μm] | 0.3 – 3.2 | 0.5 – 3.0 | 0.3 – 3.0 | 0.3 – 3.0 | 0.3 – 3.0 | 0.3 – 2.9 | 0.3-3.0 |
| Number of images<sup>b</sup> | 4,571 (5,908) | 2,304 (3,623) | 5,858 (7,121) | 6,617 (7,023) | 12,475 | 936 (1,064) | 2,970 (3,336) |

<sup>a</sup>Combined from two data sets (4 °C and 25 °C)
<sup>b</sup>In parenthesis is the initial number of images

### Different conformations in the strong-ADP state of myosin-V and myosin-IB

The structure of F-actin decorated with myosin-V in complex with Mg^2+^-ADP represents the strong-ADP state, which directly precedes the nucleotide-free rigor state within the myosin motor cycle. The overall structure encompasses all hallmarks of the strong-ADP state including a closed actin binding cleft, that allows strong binding to F-actin, and a post-powerstroke lever arm orientation (Figure 1, Figure 1—Video 1), in line with an earlier medium-resolution structure of the same complex (Wulf et al., 2016).

**Figure 1.**
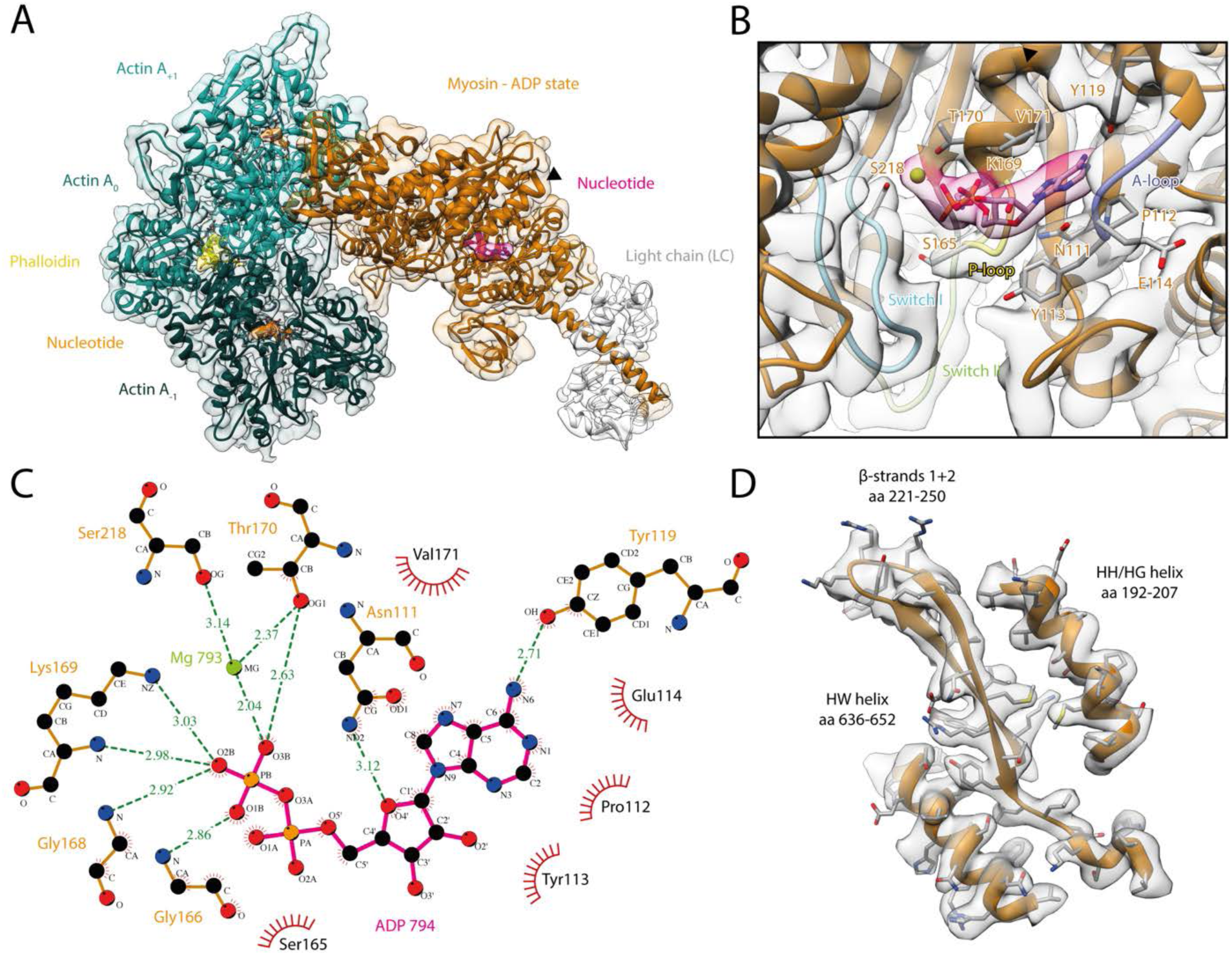
Structure and active site of the aged actomyosin-V complex bound to ADP. **(A)** Atomic model and LAFTER density map of the central myosin-V-LC subunit (orange, LC: white) bound to aged F-actin-PHD (shades of sea green, three subunits shown, A_-1_ to A_+1_). Nucleotides and PHD are highlighted in orange, pink and yellow, respectively. The HF helix is marked by a black arrowhead. **(B)** Close-up view of the myosin active site consisting of the P-loop (yellow, 164-168), switch I (blue, aa 208-220), switch II (green, aa 439-448) and the A-loop (purple, aa 111-116). Only side chains involved in the binding of ADP are displayed, also see Figure 1—figure supplement 1. **(C)** 2D protein-ligand interaction diagram illustrating the coordination of Mg^2+^-ADP by hydrogen bonds (dashed green lines) and hydrophobic interactions (red rays). **(D)** Illustration of the model-map agreement within a central section of myosin. Most side chains are resolved by the post-refined density map (transparent gray). See Figure 1—Video 1 for a three-dimensional visualization and Figure 1—figure supplement 2 for a comparison of the strong-ADP state of different myosins. **Figure 1—Video 1. Structure of the aged actomyosin-V complex bound to ADP.** Three-dimensional visualization of the aged actomyosin-V complex bound to ADP. Overview of the **(A)** overall structure, **(B)** active site and **(C)** a central section of myosin, as shown in Figure 1 A, B and D.

The density corresponding to Mg^2+^-ADP is pronounced, indicating high to complete saturation of the active site (Figure 1, Figure 1—figure supplement 1). The β-phosphate of ADP is tightly coordinated by the P-loop (aa 164-168) via a conserved Walker-A nucleotide binding motif (Walker, Saraste, Runswick, & Gay, 1982), which is also found in other ATPases as well as G-proteins (Kull & Endow, 2013; Vale, 1996). The HF helix (aa 169-183) and switch I (aa 208-220) mediate additional contacts by either directly binding to the β-phosphate or coordinating the Mg^2+^ ion (Figure 1B,C). The third key loop of the active site, switch II (aa 439-448), does not directly contribute to the binding of Mg^2+^-ADP, which is in agreement with its proposed role in ATP hydrolysis and the subsequent release of the inorganic phosphate (Sweeney et al., 2020). Yet, switch II contributes to the stability of the active site by forming a hydrogen bond with the HF helix (D437-T170, predicted by PDBsum (Laskowski, Jabłońska, Pravda, Vařeková, & Thornton, 2018)). In addition to the coordination of the β-phosphate, ADP binding is mediated by primarily hydrophobic interactions of the adenosine moiety with the purine binding loop (Bloemink, Hsu, Geeves, & Bernstein, 2020) (aa 111-116) – for brevity, hereafter referred to as A-loop (adenosine-binding loop) (Figure 1B,C, Figure 1—Video 1, Figure 1—figure supplement 1). A tyrosine (Y119) trailing the A-loop forms another putative hydrogen bond with the adenosine, completing the coordination of ADP.

The coordination of Mg^2+^-ADP in the strong-ADP state of myosin-V differs noteworthy from the one found in the so-called weak-ADP rigor-like crystal structure of the same myosin obtained in the absence of actin (Coureux et al., 2003). At the same time, it closely resembles the one reported for the strong-ADP state of myosin-IB (Mentes et al., 2018). Solely the position of switch I differs appreciably as it localizes closer to the nucleotide and converter domain in myosin-IB, corresponding to a shift of 2-3 Å relative to the structure of myosin-V. This difference highlights, that while the general architecture of the active site is common to all myosins, small local reorganizations occur and possibly account for the different kinetics within the myosin superfamily. In contrast to the resemblance of the active site, the overall structures of the strong-ADP state of myosin-V and myosin-IB differ considerably, resulting in lever arm orientations deviating by 71° (Figure 1—figure supplement 2).

### Structural transition of myosin-V upon ADP release

The structure of the actomyosin-V complex in absence of any nucleotide in myosin represents the rigor state (Figure 2, Figure 2—Video 1). In addition to an unoccupied and open active site (Figure 2—Video 1, Figure 1—figure supplement 1), the actin binding cleft is closed, facilitating strong binding to F-actin, and the lever arm adopts a post-powerstroke orientation (Figure 2, Figure 2—Video 1). These features are common to all rigor structures solved to date (Banerjee et al., 2017; Behrmann et al., 2012; Doran et al., 2020; Fujii & Namba, 2017; Gong et al., 2021; Gurel et al., 2017; Mentes et al., 2018; Risi et al., 2020; Robert-Paganin et al., 2021; Vahokoski et al., 2020; von der Ecken et al., 2016). Yet, the structures of different myosins vary, in particular in the orientation of the lever arm (Figure 1—figure supplement 2A).

**Figure 2.**
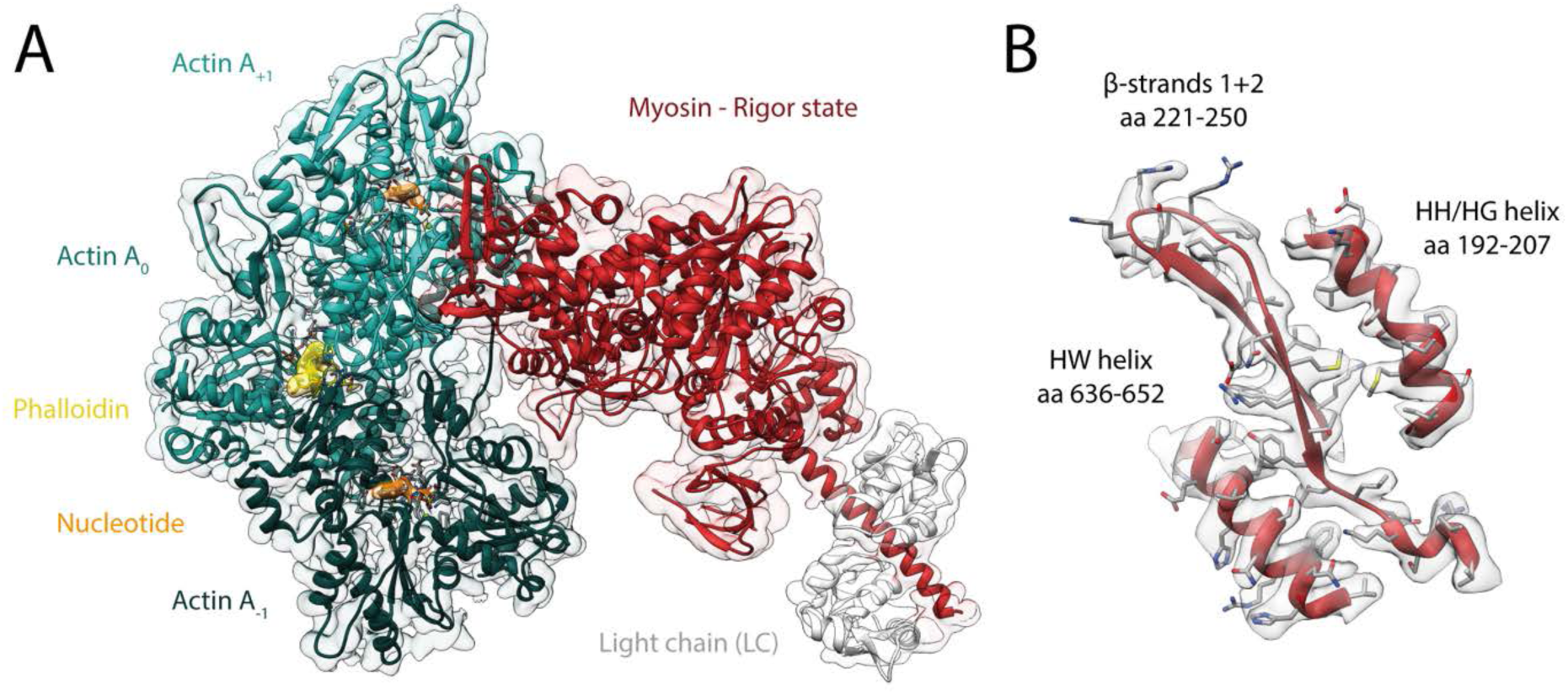
Structure of the aged actomyosin-V complex in the rigor state. **(A**) Atomic model and LAFTER density map of the central myosin-V-LC subunit (red, LC: white) bound to aged F-actin-PHD (shades of sea green, three subunits shown, A_-1_ to A_+1_). Nucleotides and PHD are highlighted in orange and yellow, respectively. **(B)** Illustration of the model-map agreement within a central section of myosin. Most side chains are resolved by the post-refined density map (transparent gray). See Figure 2—Video 1 for a three-dimensional visualization and Figure 2—figure supplement 1 for a comparison of the cryo-EM rigor structure with crystal structures of the rigor-like state. **Figure 2—Video 1. Structure of the aged actomyosin-V complex in the rigor state.** Three-dimensional visualization of the aged actomyosin-V complex in the rigor state. Overview of the **(A)** overall structure, **(B)** active site and **(C)** a central section of myosin, as shown in Figure 2.

While the actomyosin interface of the rigor state of myosin-V is basically indistinguishable from the one of the strong-ADP state, the lever arm orientations of the two states differ by ∼ 9° (Figure 3A, Figure 3—Video 1A), in agreement with a previously reported rotation of 9.5° (Wulf et al., 2016). The overall architecture of our rigor state structure not only is in good agreement with the medium-resolution cryo-EM structure published earlier (Wulf et al., 2016), but also strongly resembles the rigor-like crystal structures solved for this myosin isoform (Figure 2—figure supplement 1) (Coureux et al., 2003; Coureux, Sweeney, & Houdusse, 2004). Nevertheless, the structures are not identical and show remarkable differences in both the actin interface and lever arm orientation, which can be readily attributed to the absence of F-actin and crystal packing, respectively.

**Figure 3.**
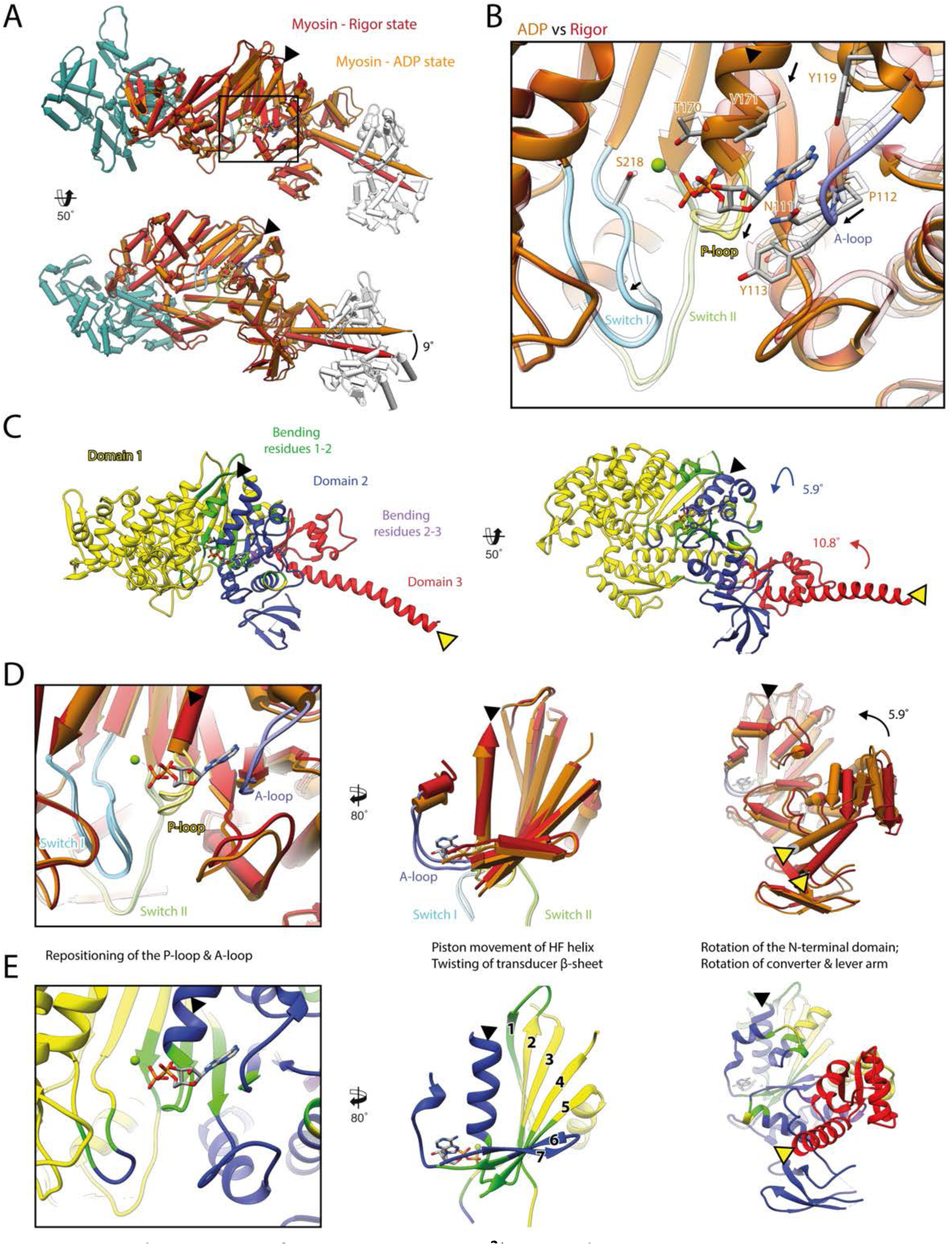
Structural transition of myosin-V upon Mg^2+^-ADP release. **(A)** Superposition of the strong-ADP (orange) and rigor (red) atomic models. Changes at the active site (black box) are not transmitted to the actomyosin interface, but to the N-terminal and converter domain resulting in a lever swing of 9°. **(B)** Close-up view of the active site showing the structural rearrangements upon Mg^2+^-ADP release (indicated by black arrows). The rigor structure is shown as transparent, see Figure 1 for color code. **(C)** Illustration of domain movements associated with Mg^2+^-ADP release predicted by DynDom (Hayward & Lee, 2002). **(D)** Scheme illustrating the structural changes associated with Mg^2+^-ADP release. **(E)** Same views as in D), but colored by DynDom domains, also see C). The HF helix and the lever arm are highlighted by a black and a yellow arrowhead, respectively. Models were aligned on F-actin. See Figure 3—Video 1 for a three-dimensional visualization. **Figure 3—Video 1. Structural transition of myosin-V upon ADP release.** Three-dimensional visualization of the structural transition of myosin-V upon Mg^2+^-ADP release. **(A-B)** Morph from the strong-ADP (orange) to the rigor state (red) of myosin-V colored by state **(A)** and DynDom domains **(B)**, for color code see Figure 3. **(C-E)** and **(F-H)** Animated scheme illustrating the structural changes associated with Mg^2+^-ADP release as shown in Figure 3 C, D.

As the strong-ADP and rigor state represent sequential states within the myosin motor cycle, a comparison of the respective high-resolution structures allows the detailed description of the structural transition of myosin-V upon Mg^2+^-ADP release (Figure 3, Figure 3—Video 1). In addition to the ∼ 9° lever arm rotation described above (Figure 3A), the two sequential states differ primarily in their conformation of the central transducer β-sheet and the N-terminal domain, which twist and rotate, respectively (Figure 3C-E, Figure 3—Video 1). Notably, the structural changes are not transmitted to the U50 and L50 domains and thus do not alter the actin binding interface (Figure 3A, C, Figure 3—Video 1A-B).

The transducer rearrangements are directly linked to a reorganization of the active site that accounts for the reduced Mg^2+^-ADP affinity of the rigor state. By promoting a piston movement of the HF helix, twisting of the transducer increases the distance between the P-loop and switch I. In the rigor state, the P-loop and the A-loop shift considerably outward (away from switch I) thereby opening the active site (Figure 3B, D, Figure 3—Video 1A-C). The resulting conformation is incompatible with the Mg^2+^-coordinating hydrogen bond between the HF helix and switch II (T170-D437). Loss of Mg^2+^ is thought to lead to the weak-ADP state of myosin (Coureux et al., 2004), which is named after its poor nucleotide affinity and promotes the release of ADP. The subsequent rigor state is stabilized by a new network of hydrogen bonds formed between lysine K169 (HF helix, previously coordinated to the β-phosphate of ADP), and aspartate D437 and isoleucine I438 (switch II). The accompanying conformation of the P-loop further reduces the ADP affinity of the rigor state (Coureux et al., 2003).

Upon Mg^2+^-ADP release, the A-loop also undergoes a small lateral shift (Figure 3B, D-E, Figure 3—Video 1C-D). In this way, it likely stabilizes the twisting of the transducer and the N-terminal domain rotation. Surprisingly, the role of the A-loop in both the coordination of ADP and the coupling of the active site to the periphery has been hardly recognized before, although it is also involved in nucleotide binding in other myosins (Bloemink et al., 2020). Given their central importance for the coordination of Mg^2+^-ADP (Figure 1), we propose that the P-loop, the A-loop and switch I contribute to the sensing of the nucleotide state and its transmission from the nucleotide binding pocket to the periphery. Their mutual interplay defines the orientation of the N-terminal domain relative to the U50 and L50 subdomains. In this way, small changes in the active site (∼1-2 Å) are amplified into significant rotations of the N-terminal and converter domain, eventually leading to a lever arm swing of ∼ 9° upon Mg^2+^-ADP release (Figure 3, Figure 3—Video 1).

Our high-resolution structures of the strong-ADP and rigor state are consistent with the sequential release of Mg^2+^ and ADP due to the isomerization of myosin to a conformation with reduced nucleotide affinity. In line with this, ADP binding to the rigor state can favor the reversal of this isomerization in presence of Mg^2+^.

An independent analysis of domain motions using DynDom (Hayward & Lee, 2002) endorses the described structural changes of myosin-V upon Mg^2+^-ADP release (Figure 3C,E, Figure 3—Video 1,F-H). Specifically, it reported a rotation of the N-terminal domain by ∼ 6° alongside to a ∼ 11° rotation of the converter domain, which ultimately causes a small lever arm swing. The semi-independent rotation of the converter domain is likely due to its coupling to both the N-terminal domain and the relay helix (aa 449-479), which is part of the U50 domain and does not change upon Mg^2+^-ADP release (Figure 3—Video 1B,H).

A similar structural transition upon Mg^2+^-ADP release has been reported for myosin-IB, V and VI based on medium- and high-resolution cryo-EM structures (Gurel et al., 2017; Mentes et al., 2018; Wulf et al., 2016), suggesting a common coupling mechanism. Although most of the details are intriguingly similar, for example the remodeling of hydrogen bonds due to the piston movement of the HF helix (Mentes et al., 2018), we still find two notable differences.

First, the extent of the lever arm swing associated with Mg^2+^-ADP release differs strongly between myosin isoforms. Whereas we find a small rotation of ∼ 9° for myosin-V (Figure 3), much larger rotations have been reported for myosin-IB (25°) (Mentes et al., 2018) and myosin-VI (30°) (Gurel et al., 2017). The extent of the swing could be directly related to the force-sensitivity of myosin, which is almost 40-fold stronger for myosin-IB compared to myosin-V (Laakso et al., 2008; Veigel, Schmitz, Wang, & Sellers, 2005). It is well known that the lifetime of the strong-ADP state is prolonged under load, preventing early detachment and increasing processivity (Kovács, Thirumurugan, Knight, & Sellers, 2007; Laakso et al., 2008; Takagi, Homsher, Goldman, & Shuman, 2006; Veigel et al., 2005). Prolongation of the strong-ADP state has been suggested to be due to the inhibition of the lever arm swing associated with ADP release (Mentes et al., 2018). Since load will more easily prevent the isomerization if Mg^2+^-ADP release requires a large converter swing, we propose that the force sensitivity increases with the extent of the lever arm swing upon Mg^2+^-ADP release.

Second, partial unwinding of the relay helix has been reported for myosin-IB and myosin-VI (Gurel et al., 2017; Mentes et al., 2018). Considering the marked difference in the lever arm swings of the different myosins, unwinding of the relay helix is probably a prerequisite for large swings, as found in myosin-IB and VI. A comparison of the structures of different myosins illustrates, that differences in the overall conformation lead to different lever arm orientations in both the rigor and strong-ADP state (Figure 1—figure supplement 2). These variations highlight the need to solve all key states of the motor cycle for a single myosin to reliably describe its structural transitions and ultimately the force generation mechanism.

### AppNHp gives rise to a strongly bound post-rigor transition state

We determined the structure of F-actin-myosin-V in complex with the non-hydrolyzable ATP analog AppNHp with the aim to characterize a potentially short-lived, weakly-bound state of myosin. The resulting cryo-EM density map shows strong density for AppNHp, indicating high to complete saturation (Figure 4, Figure 4—Video 1, Figure 1—figure supplement 1). Interestingly, the density also suggests the presence of two ions, both likely corresponding to Mg^2+^, given the size of the density and the buffer composition. While one ion occupies approximately the position that Mg^2+^ takes in the active site of the strong-ADP state, namely close to the γ-phosphate of AppNHp, the other one resides in between the α- and β-phosphates of AppNHp (Figure 4, Figure 1).

**Figure 4.**
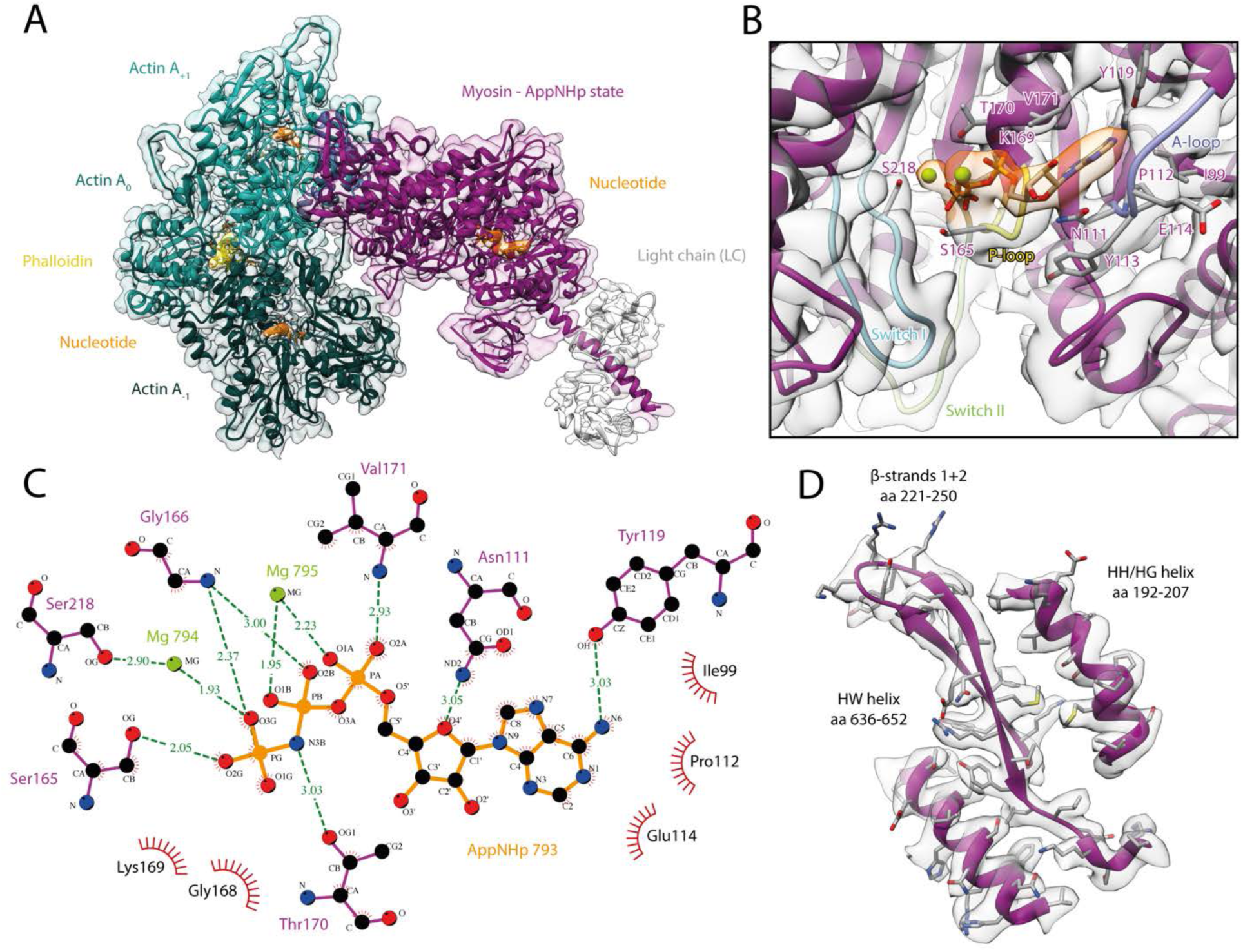
Structure and active site of the aged actomyosin-V complex bound to AppNHp. **(A)** Atomic model and LAFTER density map of the central myosin-V-LC subunit (purple, LC: white) bound to aged F-actin-PHD (shades of sea green, three subunits shown, A_-1_ to A_+1_). Nucleotides and PHD are highlighted in orange and yellow, respectively. **(B)** Close-up view of the myosin active site, see Figure 1 for color code. Only side chains involved in the binding of AppNHp are displayed. The density suggests the presence of two Mg^2+^ ions coordinating the γ, and α- and β-phosphate, respectively; also see Figure 1—figure supplement 1 and Figure 4—Video 1. **(C)** 2D protein-ligand interaction diagram illustrating the coordination of Mg^2+^-AppNHp by hydrogen bonds (dashed green lines) and hydrophobic interactions (red rays). **(D)** Illustration of the model-map agreement within a central section of myosin. Most side chains are resolved by the post-refined density map (transparent gray). See Figure 4—figure supplements 1-3 for comparisons of the AppNHp-myosin-V structure with other structures as well as an analysis of unbound myosin in the AppNHp data set. **Figure 4—Video 1. Structure of the aged actomyosin-V complex bound to AppNHp.** Three-dimensional visualization of the aged actomyosin-V complex bound to AppNHp. Overview of the **(A)** overall structure, **(B)** active site and **(C)** a central section of myosin, as shown in Figure 4 A,B and D.

Similar to ADP, AppNHp is coordinated by a network of hydrogen bonds and additional hydrophobic interactions with the P-loop, switch I and the A-loop (Figure 4C, Figure 1C). The details of the interactions, however, differ due to the different sizes of the two nucleotides and their relative positions in the active site, i.e. the γ-phosphate of AppNHp almost takes the position of the β-phosphate of ADP relative to the HF helix (Figure 4, Figure 1—figure supplement 1, Figure 1).

Surprisingly, and in contrast to previous low-resolution cryo-EM results (Volkmann et al., 2005), the overall structure of AppNHp-bound myosin-V is reminiscent of the rigor state (Figure 4—figure supplement 1). In particular, myosin is strongly bound to F-actin and adopts a post-powerstroke lever arm orientation (Figure 4, Figure 4—Video 1, Figure 4—figure supplement 1A-B). The active site of AppNHp-bound myosin also closely resembles that of the rigor state, and thereby significantly deviates from the conformation found in the strong-ADP state (Figure 3—figure supplement 1C-D).

The compatibility of an ATP analog, specifically the presence of a γ-phosphate at the active site, with strong F-actin binding is initially puzzling and seemingly at odds with the reported reciprocal nature of these two processes (Coureux et al., 2004; Kühner & Fischer, 2011). A comparison of our AppNHp-bound structure with a rigor-like crystal structure of myosin-V with ADP weakly-bound to its active site (Coureux et al., 2004) resolves this conflict (Figure 4—figure supplement 2A). The position and coordination of AppNHp and ADP in these two structures is almost identical, suggesting that AppNHp is only weakly bound in our structure and therefore compatible with strong F-actin binding. Interestingly, a similar coordination was observed for Mg^2+^-ADP in a putative strong-ADP to rigor transition state cryo-EM structure of myosin-IB (Mentes et al., 2018) (Figure 4—figure supplement 2B). These comparisons indicate that AppNHp and ADP can both weakly bind to myosin in a conformation reminiscent of the rigor.

Based on these results, we propose that our AppNHp-bound myosin-V structure represents a post-rigor transition (PRT) state that allows to visualize how ATP binds in the rigor state, prior to the transition that promotes its detachment from F-actin. The characteristic weak coordination of AppNHp in the PRT state allows myosin to remain strongly bound to F-actin until a strong coordination of the nucleotide is established. The report of a transition state with weakly-bound ADP (Mentes et al., 2018) (Figure 4—figure supplement 2B) suggests that weak nucleotide binding is a common scheme and that the PRT state is therefore not limited to AppNHp. The visualization of an ATP analog bound to a state reminiscent of the rigor shows that ATP mainly binds via its adenine ring, as does ADP (Figure 1). It also explains how the γ phosphate can fit into the relatively small pocket created by the rigor conformation of the P-loop (Figure 4), and how its presence leads to local changes of the active site facilitating a tight coordination (Figure 4—figure supplement 1). In this way, the PRT state provides new insights on how myosin detaches from F-actin and indicates that the theoretical weakly-bound post-rigor state (Sweeney & Houdusse, 2010; Walklate, Ujfalusi, & Geeves, 2016) is unlikely to be populated within the motor cycle.

Although we find myosin-V-AppNHp strongly bound to F-actin in the PRT state (Figure 4—figure supplement 1), we had to significantly increase the myosin concentration to achieve full decoration of actin filaments (see Methods for details), in agreement with a weaker binding affinity (Konrad & Goody, 1982; Yengo et al., 2002). We therefore conclude that AppNHp can potentially lead to different structural states, similar to ADP in myosin-IB (Mentes et al., 2018). Likely due to large differences in the binding affinity of these states or rapid detachment of myosin from F-actin , we only find myosin bound to F-actin in the PRT state. In line with this assumption, we find a significant amount of unbound myosin in the background of our AppNHp data sets (Figure 4—figure supplement 3). The 3D reconstruction and thus identification of the structural state of the background myosin was unfortunately impeded by a strong orientational preference of the myosin particles (Figure 4—figure supplement 3B). Further studies are therefore required to test the conformation of AppNHp-bound myosin-V in absence of F-actin.

### Conservation and specificity of the actomyosin-V interface

A comparison of the three states of the actomyosin-V complex (strong-ADP, rigor and PRT state) reveals a striking similarity of the actomyosin interface (Figure 5, Figure 5— Video 1). The atomic models superimpose almost perfectly with only little variations in the orientation of some incompletely resolved side chains. The remarkable similarity suggests that the same set of interactions is maintained during all strongly-bound states of the myosin motor cycle, despite their varying F-actin binding affinities. Differences in the affinity might therefore not be linked to altered contacts, but rather to the degree of structural flexibility inherent to each state (see below).

**Figure 5.**
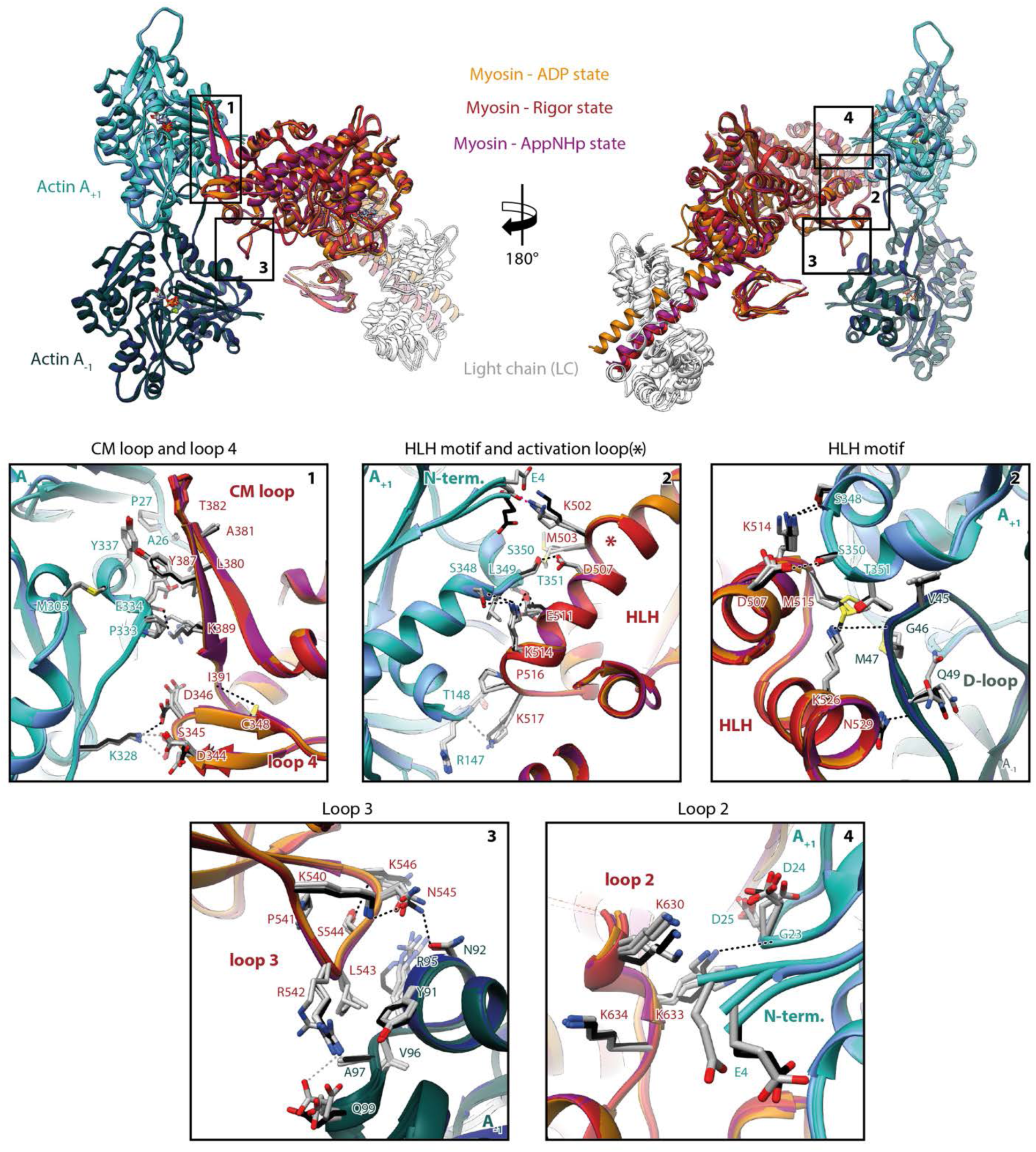
Indistinguishable actomyosin interfaces in the strong-ADP, rigor and PRT state. Comparison of the actomyosin-V interface within all three states (rigor: red, strong-ADP: orange and AppNHp-bound PRT: purple) illustrating the remarkable similarity of interactions with F-actin. **(Top)** Front and back view of the central myosin molecule and the two actin subunits it is bound to (shades of green and blue, A_+1_ and A_-1_, see Figure 1—figure supplement 1 for color code). Black boxes indicate the location of close-up views shown below. **(Bottom)** Close-up views of all actin-myosin interfaces including the cardiomyopathy (CM) loop, the helix-loop-helix (HLH) motif, loops 2-4 and the activation loop (highlighted by an asterisk). Side chains of key residues are displayed and labeled for all states (rigor: black, ADP and AppNHp: gray). Dashed lines indicate hydrogen bonds predicted for the rigor (black) and ADP/AppNHp state (gray), respectively. See Figure 5—Video 1 for a three-dimensional visualization. **Figure 5—Video 1. Conservation of the actomyosin-V interface.** Comparison of the actomyosin-V interface within all three states (rigor: red, strong-ADP: orange and AppNHp-bound PRT: purple). **(A, J)** Overview of the central myosin molecule and the two actin subunits it is bound to (shades of green and blue, A_+1_ and A_-1_, for color code see Figure 1—figure supplement 1). **(B-I)** Close-up views highlighting the localization and molecular details of all actin-myosin interfaces including the CM loop and loop 4 **(B-D)**, the HLH motif and activation loop (highlighted by an asterisk) **(D-G)**, loop 3 **(B,D,G-H)** and loop 2 **(D,G,I)**. In close-up views of specific interfaces, the structure of the aged actomyosin-V complex in the rigor state is first shown on its own, followed by a superposition of all models illustrating the remarkable similarity of the actomyosin interface. See Figure 5 for residue labels and predicted hydrogen bonds.

The actomyosin-V interface is comprised of six structural elements, namely the cardiomyopathy (CM) loop (aa 376-392), loop 4 (aa 338-354), the helix-loop-helix (HLH) motif (505-531), the activation loop (aa 501-504), loop 3 (aa 532-546) and loop 2 (aa 594-635) (Figure 5, Figure 5—Video 1 ). While these elements represent a common set of actin-binding elements, most of which have conserved hydrophobic and electrostatic properties, not all myosins utilize all of them. Moreover, the precise nature of individual interactions and the residues involved vary considerably among myosins, largely due to sequence variations known to tune the kinetic properties of myosin (Mentes et al., 2018; Robert-Paganin et al., 2021). Comparisons of the actomyosin interface of different myosins are therefore essential for identifying common and specific features of the myosin superfamily.

A detailed comparison of the actomyosin interface of myosin-V with previously published actomyosin structures (Banerjee et al., 2017; Behrmann et al., 2012; Doran et al., 2020; Gong et al., 2021; Gurel et al., 2017; Mentes et al., 2018; Risi et al., 2020; Robert-Paganin et al., 2021; Vahokoski et al., 2020; von der Ecken et al., 2016) shows many common features, but also some myosin-V specific ones. The tightest and most conserved contact is formed by the HLH motif (Robert-Paganin et al., 2021). In analogy to other myosins, it relies primarily on extensive hydrophobic contacts with F-actin, complemented by a series of hydrogen bonds (predicted by PDBsum (Laskowski et al., 2018), Figure 5, Figure 5—Video 1E-F). The comparably short CM loop of myosin-V is also highly conserved, with respect to its hydrophobic nature. However, unlike the CM loop of other myosins (Fujii & Namba, 2017; Gurel et al., 2017; Mentes et al., 2018; Risi et al., 2020; von der Ecken et al., 2016), its tip does not engage in complementary electrostatic interactions (Figure 5, Figure 5—Video 1C). The conformation we found for loop 4 differs from all others reported so far. Not only is it more compact, folding in a β-hairpin, but it also localizes closer to the base of the CM-loop, where it is stabilized by a non-conserved hydrogen bond between C348 and I391 (Figure 5, Figure 5—Video 1C). However, its electrostatic interactions with F-actin are reminiscent of those reported for other myosins (Fujii & Namba, 2017; Gurel et al., 2017; Risi et al., 2020; von der Ecken et al., 2016). Loop 2 is exceptionally long in myosin-V and only partially resolved in our structures (Figure 5—Video 1). While this is also the case for most actomyosin structures resolved so far (Banerjee et al., 2017; Doran et al., 2020; Gong et al., 2021; Risi et al., 2020; Robert-Paganin et al., 2021; von der Ecken et al., 2016), loop 2 of myosin-V stands out by the unique α-helical fold of its C-terminal part (Figure 5—Video 1I). This fold facilitates a compact packing of basic residues and thereby promotes the electrostatic interactions commonly found at the loop 2 interface. The activation loop is a structural element that does not contribute to F-actin binding in all myosins (Gurel et al., 2017; Robert-Paganin et al., 2021). In myosin-V, it forms primarily electrostatic interactions with the N-terminus of F-actin, but does not lead to its ordering, as has been reported for other myosins (Figure 5, Figure 5—Video 1E) (Banerjee et al., 2017; Behrmann et al., 2012; Fujii & Namba, 2017; Mentes et al., 2018; Vahokoski et al., 2020). The last structural element involved in actin binding is loop 3. It forms the so-called Milligan contact (Milligan, Whittaker, & Safer, 1990), which is strong in myosin-V and includes electrostatic and hydrophobic interactions as well as several hydrogen bonds (Figure 5, Figure 5—Video 1H). The contact is furthermore strengthened by hydrogen bonds between K540-N545 and S544-K546 that stabilize the conformation of loop 3. Interestingly, a strong Milligan contact has also been reported for the high-duty ratio myosins IB and VI (Gurel et al., 2017; Mentes et al., 2018), whereas no or only weak interactions were found in class II myosins (Doran et al., 2020; Fujii & Namba, 2017; Risi et al., 2020; von der Ecken et al., 2016). We therefore speculate that an intimate Milligan contact might be a general feature of high-duty ratio myosins, which need to bind particularly tightly to F-actin to fulfill their function as cargo-transporters or molecular anchors.

In summary, we demonstrated that myosin-V establishes a maximum of contacts with F-actin, utilizing all six potential binding elements (Figure 5, Figure 5—Video 1E-F,). In addition, we have identified a previously unseen α-helical fold of the C-terminus of loop 2 (Figure 5, Figure 5—Video 1I), which possibly strengthens the interactions at this interface.

### Myosin-V specifically selects the closed D-loop conformation of F-actin

To assess the structural effect of myosin binding on F-actin, we compared the structure of aged F-actin-PHD in the presence (rigor state, representative for all states) and absence of myosin-V (PDB: 6T20, (Pospich, Merino, & Raunser, 2020)) (Figure 6). The observed differences are subtle and primarily involve the DNase-binding loop (D-loop, aa 39-55) of F-actin and loops known for their flexibility (Pospich et al., 2020). The most prominent alteration involves glutamine Q49 within the D-loop, which moves away from the actomyosin interface by ∼2 Å to enable the formation of a hydrogen bond with N529 in the HLH motif of myosin (Figure 6B, Figure 5). Similar, but not identical, subtle changes have been reported for other actomyosins (Behrmann et al., 2012; Gong et al., 2021; Gurel et al., 2017; Robert-Paganin et al., 2021; von der Ecken et al., 2016), in addition to an ordering of the N-terminus of actin (Banerjee et al., 2017; Behrmann et al., 2012; Fujii & Namba, 2017; Mentes et al., 2018; Vahokoski et al., 2020; von der Ecken et al., 2016), which we do not observe for myosin-V.

**Figure 6.**
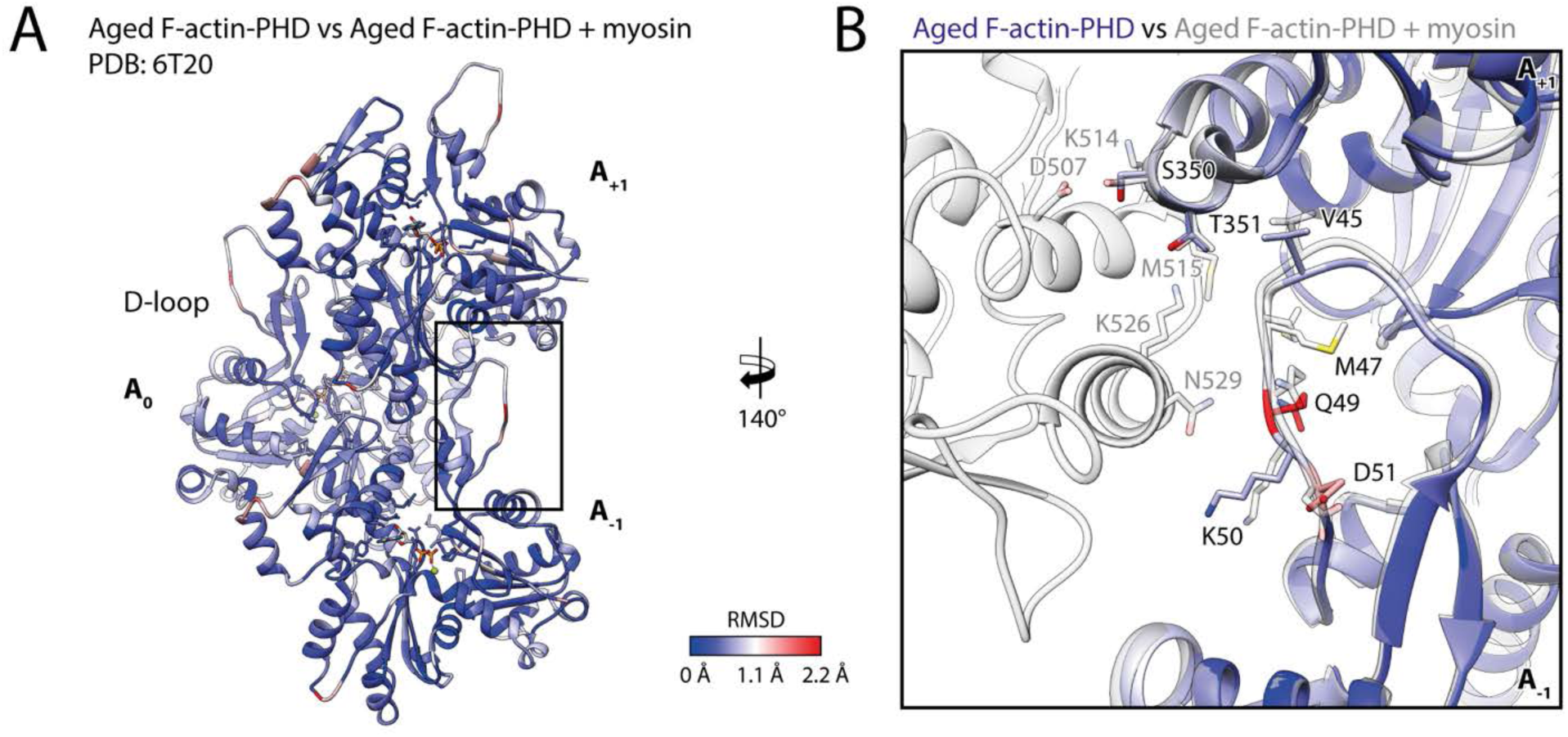
Myosin binding gives rise to subtle structural changes of aged PHD-stabilized F-actin. Illustration of the structural similarity of aged F-actin-PHD in absence and presence of myosin. **(A)** Atomic model of aged F-actin-PHD (PDB: 6T20, (Pospich et al., 2020), three subunits shown, A_-1_ to A_+1_) color-coded by the backbone root mean square deviation (RMSD) of this structure with the one of aged F-actin-PHD decorated with myosin-V in the rigor state. **(B)** Close-up view of the D-loop interface illustrating that the structural changes associated with myosin binding are small. For a direct comparison, the atomic model of the rigor actomyosin-V complex is superimposed (transparent gray). F-actin subunits were aligned individually to account for errors in the calibration of the pixel size. See Figure 6—Table 1 for a comparison of helical symmetry parameters.

Notably, our data show no significant change of the helical symmetry parameters upon myosin binding, neither in rigor nor in any other state of myosin (Figure 6—Table 1). This is in stark contrast to an earlier medium-resolution study of myosin-V, which reported additional twisting of PHD-stabilized F-actin in dependence of the nucleotide state of myosin (Wulf et al., 2016).

It was reported that myosin-V is sensitive to the nucleotide state of F-actin and prefers young PHD-stabilized F-actin over aged F-actin-PHD (Zimmermann et al., 2015). We have recently shown that young ATP/ADP-P_i_-bound and aged ADP-bound F-actin primarily differs in its conformation of the D-loop-C-terminus interface and that actin-binding proteins like coronin-IB (Cai, Makhov, & Bear, 2007) probably read the nucleotide state of F-actin from this interface (Merino et al., 2018). We have furthermore shown that the short-lived ATP/ADP-P_i_-bound state of F-actin can be specifically stabilized using either PHD (Lynen & Wieland, 1938) or jasplakinolide (JASP) (Crews, Manes, & Boehler, 1986; Pospich et al., 2020). To reveal the structural mechanism by which myosin-V senses the nucleotide state of F-actin, we have solved the structure of myosin-V in the rigor state in complex with young JASP-stabilized F-actin (F-actin-JASP) to 3.2 Å (Figure 7, Figure 7— Table 1, Figure 7—figure supplement 1, Table 1—Supplementary Figure 1, Table 1). The atomic model of myosin in this structure superimposes perfectly with the one bound to aged F-actin-PHD (Figure 7C-D), indicating that the nucleotide state of F-actin has no structural effect on myosin-V in the rigor state. Surprisingly, and despite having ADP-P_i_ bound to its active site (Figure 1—figure supplement 1), F-actin adopts the closed D-loop state, which is characteristic for aged ADP-bound F-actin (Figure 7) (Merino et al., 2018). However, a control structure of F-actin-JASP alone (3.1 Å, Figure 7—figure supplement 1, Figure 7— Table 1, Table 1, Table 1—Supplementary Figure 1) confirms that actin was successfully stabilized in the desired young state, having a characteristic open D-loop conformation (Figure 7—figure supplement 2) and ADP-P_i_ bound to its active site (Figure 1—figure supplement 1). Thus, we conclude that binding of myosin-V to young F-actin-JASP induces structural changes that ultimately result in the closed D-loop conformation (Figure 8, Figure 8—Video 1, Figure 8—figure supplement 1), thereby abolishing the effect of JASP (Pospich et al., 2020). Interestingly, our data show that the open D-loop state would not clash with bound myosin (Figure 8C-D). The closed conformation may therefore be selected for its superior shape complementarity to myosin, which possibly establishes a strong-binding interface between the D-loop and HLH motif and by doing so contributes to the high-binding affinity of the rigor state (Figure 8).

**Figure 7.**
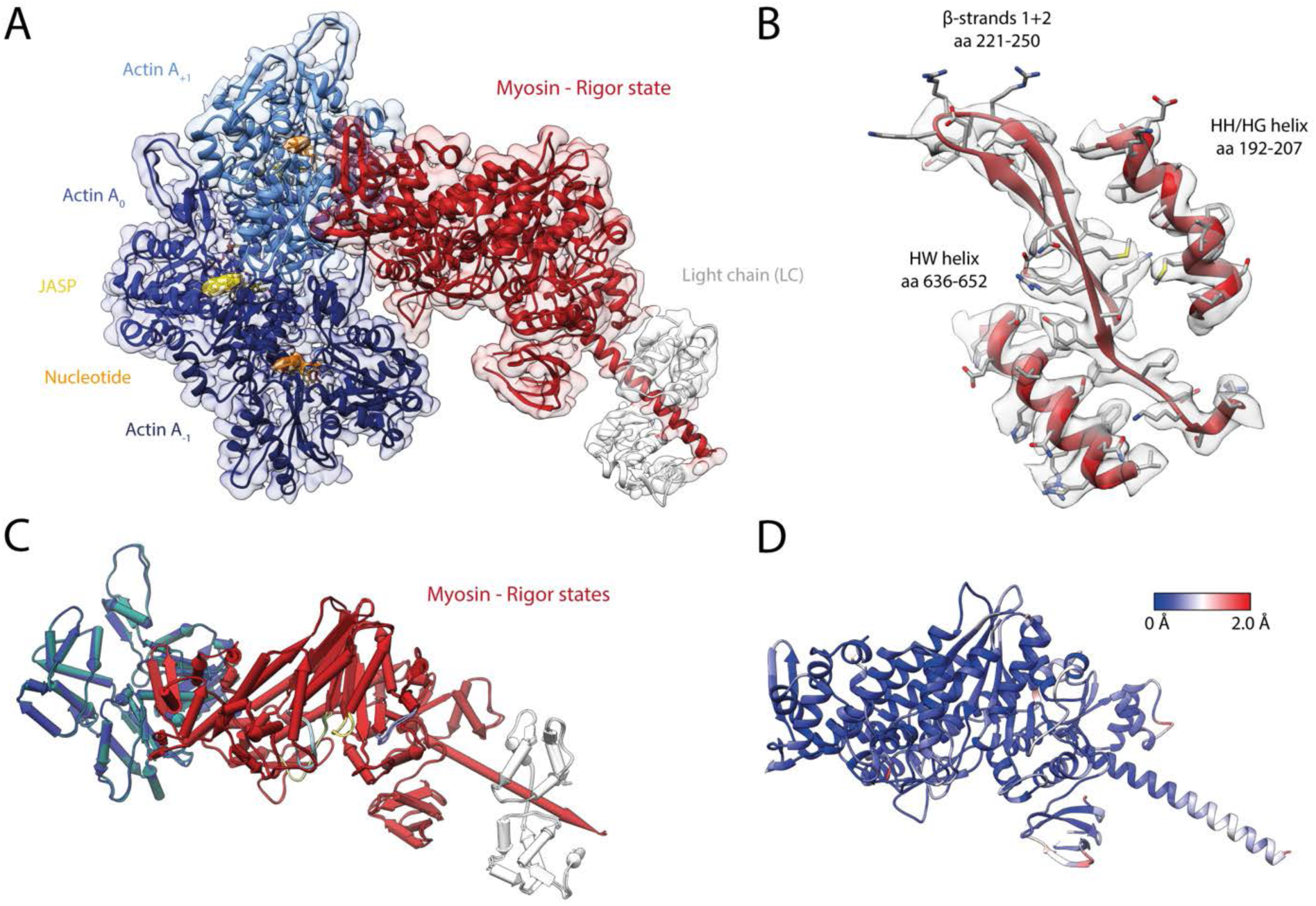
Structure of the young actomyosin-V complex in the rigor state. **(A)** Atomic model and LAFTER density map of the central myosin-V-LC subunit (red, LC: white) bound to young F-actin-JASP (shades of blue, three subunits shown, A_-1_ to A_+1_). Nucleotides and JASP are highlighted in orange and yellow, respectively; also see Figure 8—Video 1F-H. **(B)** Illustration of the model-map agreement within a central section of myosin. Most side chains are resolved by the post-refined density map (transparent gray). **(C)** Superposition and **(D)** color-coded root mean square deviation (RMSD) of the young and aged actomyosin-V complex in the rigor state illustrating their structural identity. Residues with increased RMSD solely localize to regions of lower local resolution and can therefore be explained by modeling inaccuracies. See Figure 7—figure supplement 1 and Figure 7—Table 1 for an overview of the cryo-EM data and refinement and model building statistics, respectively. The structure of young F-actin-JASP in absence of myosin is shown in Figure 7—figure supplement 2.

**Figure 8.**
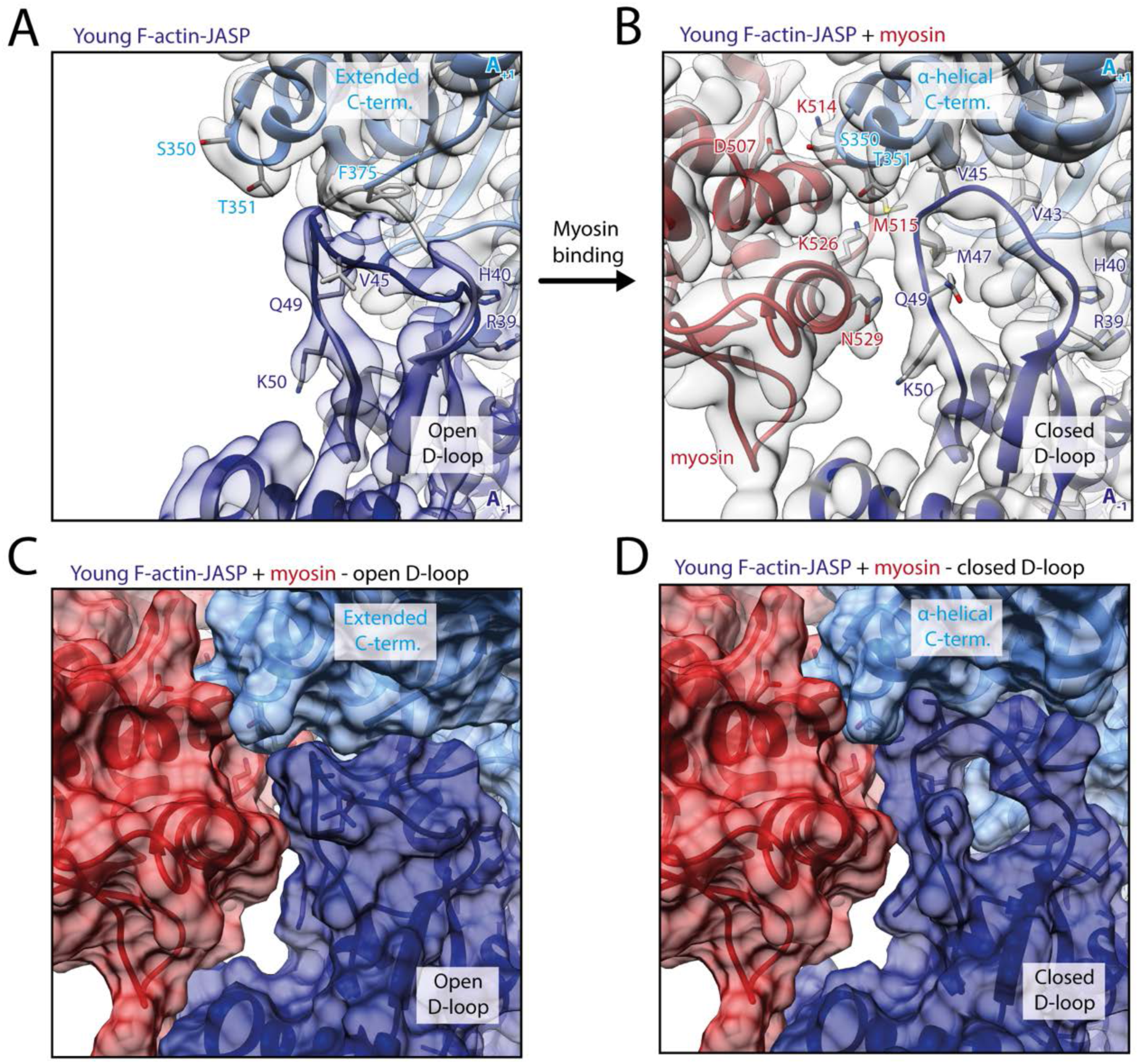
Myosin-V binding causes closure of the D-loop in young JASP-stabilized F-actin. **(A)** Atomic model and LAFTER density map of young F-actin-JASP (shades of blue, subunits A_-1_, and A_+1_). Before myosin binding, the D-loop primarily adopts the open conformation and the C-terminus is extended. A superimposed atomic model (gray) highlights a minor density potentially corresponding to the closed D-loop conformation. **(B)** Binding of myosin-V in the rigor state (red) causes a structural transition to the closed D-loop conformation, which comes with an α-helical C-terminus, also see Figure 8—Video 1 and Figure 8—figure supplement 1. **(C)** Surface representation of young F-actin-JASP (open D-loop, as shown in A) illustrating that the open D-loop conformation would not clash with myosin (computationally docked). **(D)** Surface representation of the young JASP-stabilized actomyosin complex (closed D-loop, as shown in B). See Figure 8—figure supplement 2 for an illustration how pyrene-labeling might interfere with myosin binding. **Figure 8—Video 1. Structural changes of JASP-stabilized F-actin upon binding of myosin-V**. **(A-C)** Overview of the structure of young F-actin-JASP. **(A)** Atomic model and LAFTER density map of the central three actin subunits (shades of blue, A_-1_ to A_+1_). The D-loop primarily adopts the open conformation typical for JASP-stabilized F-actin. Nucleotides and JASP are highlighted in orange and yellow, respectively. **(B)** Close-up view of the F-actin active site. There is clear density for an inorganic phosphate P_i_, which is characteristic for young F-actin, also see Figure 1—figure supplement 1. **(C)** Model-map agreement within a central section of F-actin. Most side chains are resolved by the post-refined density map (transparent gray, also shown in B). **(D)** Animation schematically illustrating the closure of the D-loop upon binding of myosin-V in the rigor state (red, LC: white). **(E)** Close-up view showing the structural rearrangement of the D-loop C-terminus interface of young F-actin-JASP upon myosin binding. For guidance, myosin is shown as transparent in the unbound state. **(F-H)** Overview of the structure of the young actomyosin-V complex in the rigor state. **(F)** Atomic model and LAFTER density map of the central myosin-V-LC subunit bound to young F-actin-JASP. The D-loop solely adopts a closed conformation, despite the presence of JASP (yellow). **(G)** Close-up view of the active site illustrating that myosin binding does not affect the nucleotide state of F-actin. **(H)** Model-map agreement within a central section of myosin. Most side chains are resolved within the post-refined density map (transparent gray, also shown in G). Note that the myosin-binding process is intentionally shown over-simplistically.

Our finding that myosin-V specifically selects the closed D-loop state of F-actin has interesting potential implications for the remodeling of the actin cytoskeleton (Merino et al., 2019). By inducing the aged conformation of F-actin, myosin-V potentially fosters phosphate release and thereby aging of actin filaments. This, in turn, could promote the directed depolymerization of F-actin sections after myosin-V has already passed on its way to the barbed end. In this way, myosin-V might increase the pool of monomeric actin available for filament elongation and thus ultimately facilitate faster end-to-end cargo transport.

Our structure does not provide a structural explanation for the reported nucleotide-sensitivity of myosin-V (Zimmermann et al., 2015). This could be due to three, possibly complementary, reasons. First, myosin-V might be sensitive to the nucleotide state of F-actin only in certain structural states, such as the initially binding PPS (Wulf et al., 2016) and P_i_R state (Llinas et al., 2015). Second, the structural plasticity of young ATP/ADP-P_i_-bound F-actin (Kueh & Mitchison, 2009), rather than the open D-loop conformation, might be beneficial for myosin binding. Third, the open D-loop conformation might promote the formation of initial contacts with myosin-V. Once these are established, the subsequent transition from a weak-to a strong-binding state potentially causes a structural transition of F-actin, eventually locking it in the closed D-loop conformation. In line with these theories a number of biochemical and biophysical studies suggested that a structural rearrangement of F-actin and its structural plasticity are critical for proper myosin activity (Anson et al., 1995; Drummond, Peckham, Sparrow, & White, 1990; Kim, Bobkova, Hegyi, Muhlrad, & Reisler, 2002; Nishikawa et al., 2002; Noguchi et al., 2012; Oztug Durer, Kamal, Benchaar, Chance, & Reisler, 2011; Prochniewicz & Thomas, 2001; Prochniewicz et al., 2010). Moreover, the D-loop C-terminus interface was predicted to contribute to the initial binding interface of myosin (Gurel et al., 2017; Lehman, Orzechowski, Li, Fischer, & Raunser, 2013; Risi et al., 2017; Robert-Paganin, Pylypenko, Kikuti, Sweeney, & Houdusse, 2020).

Finally, the conformational selection mechanism of myosin-V offers a structural explanation for the quenching of pyrene fluorescence upon myosin binding. Pyrene conjugated to cysteine 374 in the C-terminus of F-actin has been often used to report not only actin kinetics, but also myosin binding (Kouyama & Mihashi, 1981). Closure of the actin binding cleft of myosin is thought to expose pyrene to the solvent and thus cause fluorescence quenching (Chou & Pollard, 2020), but the exact timing and the structural basis are not yet known (Llinas et al., 2015; Robert-Paganin et al., 2020). A recent cryo-EM structure of pyrene-labeled F-actin has revealed that pyrene wedges itself between the tip of the D-loop and the hydrophobic groove surrounding it, partially pushing the D-loop out of its binding pocket (Chou & Pollard, 2020). This likely interferes with myosin selecting the closed D-loop state (Figure 8—figure supplement 2), which explains the significantly lower affinity for pyrene-labeled F-actin (Taylor, 1991). We furthermore suggest that myosin quenches the fluorescence of pyrene by pushing it out of its binding pocket when selecting the closed D-loop state during its transition to a strong binding state.

### Pronounced structural heterogeneity of myosin-V

To identify a potential mixture of structural states, we performed 3D classifications of signal-subtracted particles for all our data sets (Table 1—Supplementary Figure 1). Interestingly, the results indicate a continuous conformational heterogeneity of myosin-V, as opposed to a mixture of several discrete structural states (see Methods for details). Based on the identified 3D classes, we solved and modeled a total of 18 high-resolution (< 3.7 Å) structures of actomyosin-V (Table 1—Supplementary Figure 1, Table 1 —Supplementary Tables 1-3, Figure 7—Table 1). A superposition of all structures from one data set illustrates the nature of the underlying conformational heterogeneity (Figure 9). While the L50 domain, F-actin and the actomyosin interface vary little in all states (see also Figure 5), we observe a pronounced structural flexibility for all other domains (Figure 9A). Primarily, the U50 domain pivots and moves toward or away from the actin interface. In doing so, it causes twisting and shifting of the central transducer β-sheet, which is in turn coupled to a rotation of the N-terminal domain and eventually the converter domain (Figure 9A). In this way, pivoting of the U50 domain leads to different lever arm positions within the 3D classes of a single data set. The associated lever swing is not limited to a single axis and thus results in a two-dimensional distribution of lever arm conformations around the position within the average structure (Figure 9A, Figure 9— Videos 1-3). The extent (∼ 9° - 12°) of relative lever arm swings within this distribution is intriguing (Figure 9A, Figure 9—figure supplement 1), considering that the swing associated with Mg^2+-^ADP-release is only ∼ 9° for myosin-V (Figure 3).

**Figure 9.**
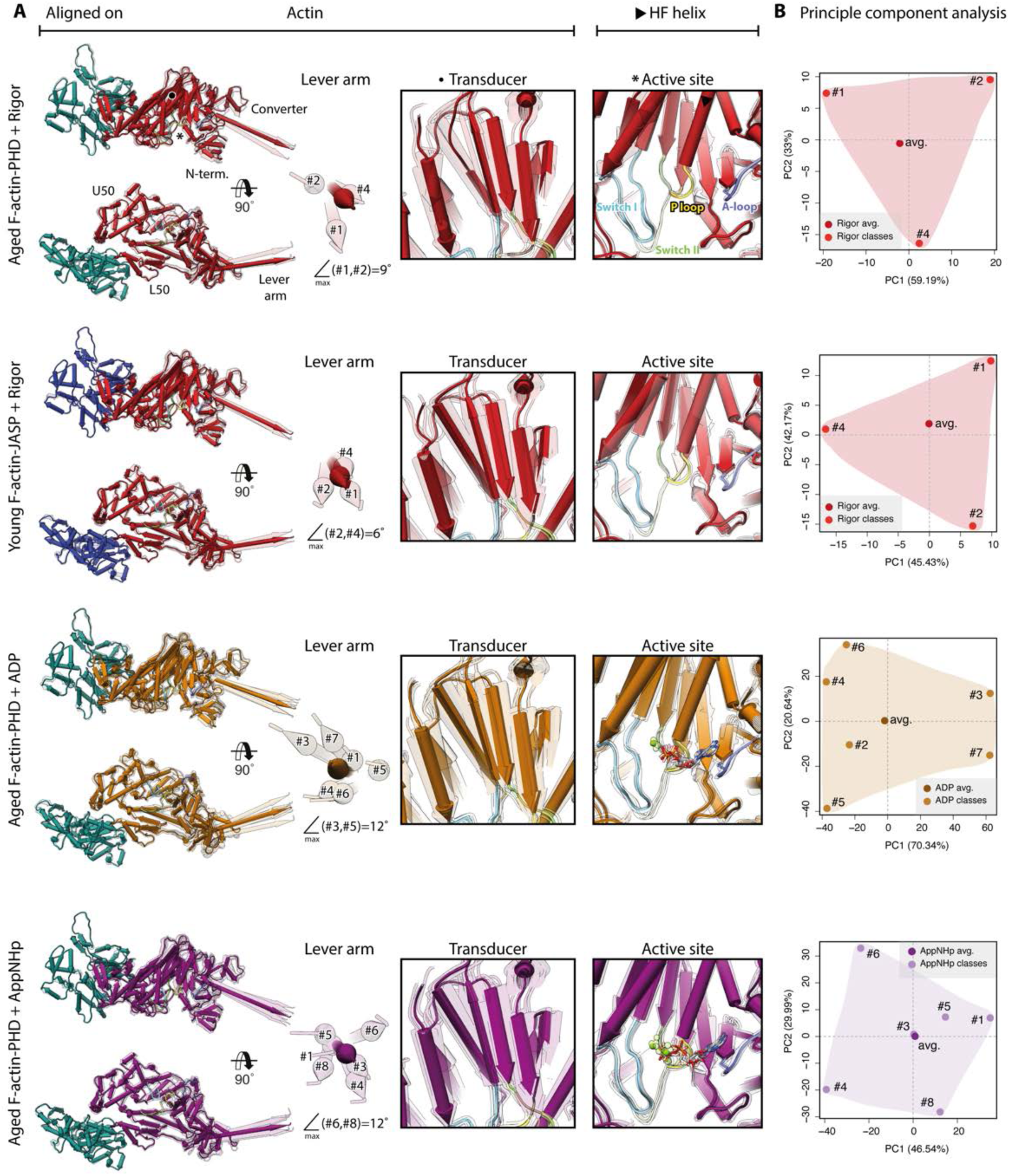
Conformational heterogeneity of myosin-V. Illustration of the conformational heterogeneity of myosin-V in the rigor (red), strong-ADP (orange) and AppNHp-bound PRT state (purple) when bound to F-actin (aged F-actin-PHD: sea green, young F-actin-JASP: blue). **(A)** Superposition of all atomic models (central 1er, average: opaque, 3D classes: transparent) built for each state. Models were either aligned on the F-actin subunit or the HF helix (indicated by black arrow head). Pivoting of the U50 domain in combination with shifting and twisting of the central transducer β-sheet results in a rotation of the N-terminal and converter domain, giving rise to an altered lever arm orientation. The extent of these changes depends on the nucleotide state and is largest in the strong-ADP and PRT state. Inserts show either the transducer β-sheet (black dot) or the active site (asterisk), which basically remains unchanged within all models of one state. **B)** Mapping of atomic models (average and 3D classes) into the first two principal components of a principal component analysis (PCA) illustrating the overall conformational space covered. Classes are labeled by their number (#1-#8, also see Table 1—Supplementary Figure 1). For a comparison of conformational extremes see Figure 9—figure supplement 1. Morphs of extremes and trajectories along the principal components are visualized in Figure 9—Videos 1-3.

Our data show that the conformational heterogeneity of myosin-V is not caused by variations of the active site or mixed nucleotide states (Figure 9). Nevertheless, the presence of a nucleotide does affect the extent of flexibility, as ADP and AppNHp lead to a greater change in lever arm position (Figure 9A). This tendency is also reflected in a principal component (PC) analysis of all models belonging to one data set (Figure 9B). The PC plots not only demonstrate that individual 3D classes represent sampling points within a continuous conformational space, but also show larger overall spaces for the ADP and AppNHp data sets.

To impartially compare the conformations of the different nucleotide states of myosin-V, we performed a PC analysis of all models (Figure 10). Within the PC plot, the average structures of the rigor and PRT state as well as their conformational spaces colocalize (Figure 10A), in agreement with their structural similarity. However, the significantly larger conformational space of the PRT data indicates a considerable difference to the rigor state. In accordance to its distinct conformation, the strong-ADP average structure does not map close to any of the other averages. Its conformational space also overlaps only partially with the PRT space and to a lesser extent with that of the young rigor state (Figure 10A). The relative mapping of individual conformational spaces is also well reflected by the lever arm positions of corresponding atomic models (Figure 10B).

**Figure 10.**
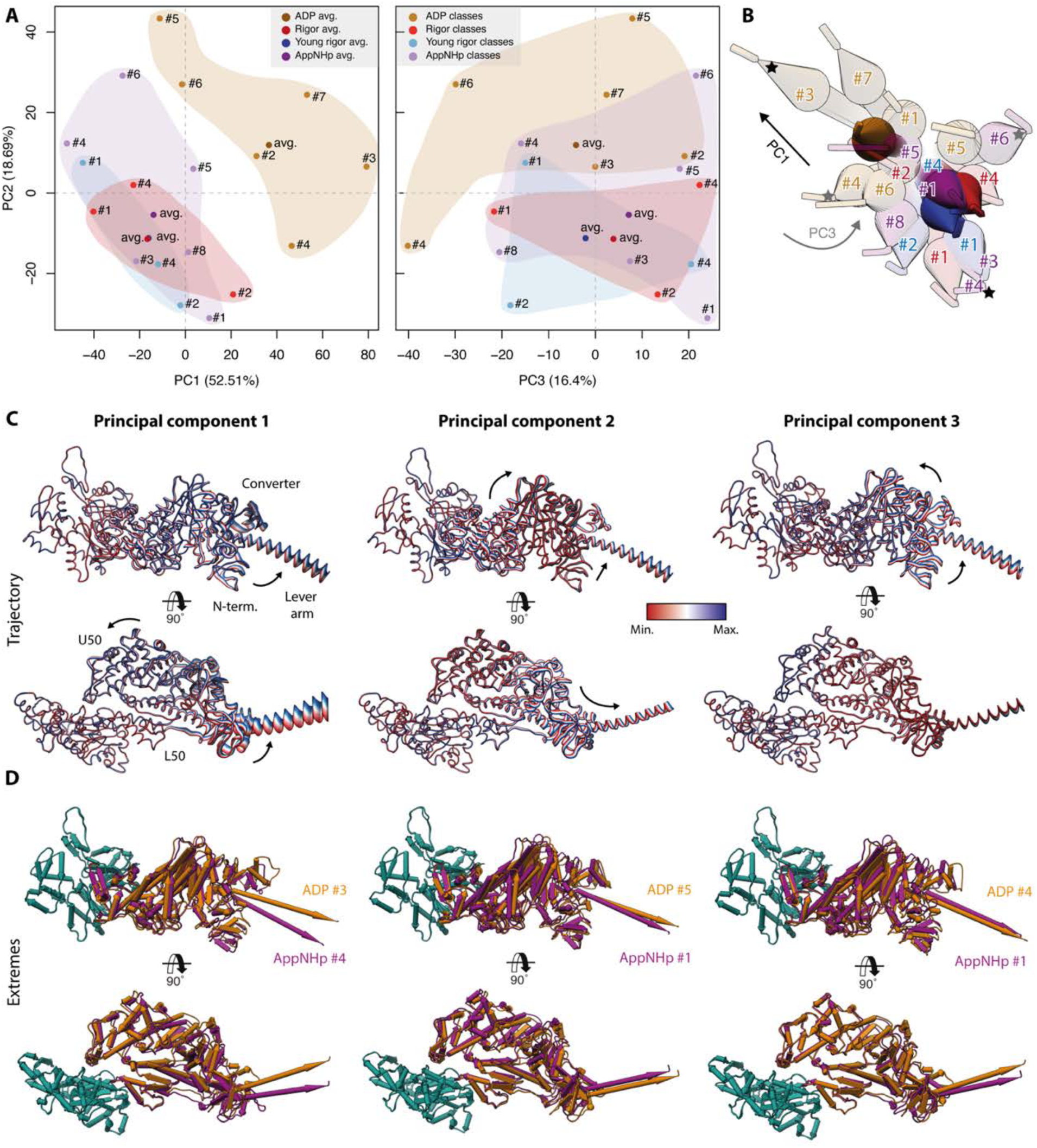
Principal component analysis of all myosin-V models. Principal component analysis of all atomic models of the actomyosin-V complex, including average and 3D class average models of the strong-ADP, rigor and PRT state (central actomyosin subunit only). **(A)** Mapping of atomic models into the first and second as well as the second and third principal component. Data points are colored by the state of the actomyosin-V complex (aged rigor: red, aged strong-ADP: orange, aged AppNHp-bound PRT: purple and young rigor: blue). Atomic models of average structures are shown as opaque and models of 3D classes as transparent. The conformational space covered within each state is indicated by a correspondingly colored 2D polygon. **(B)** Superposition of all lever arm positions. Changes along the first and third principal component are highlighted by a black and gray arrow, respectively (extremes marked with asterisks). **(C)** Color-coded trajectories along the first, second and third principal component (red minimum, blue maximum). Arrows indicate the mapped conformational changes. **(D)** Same views as in C) but showing the extreme structures along each principal component, see Figure 9 for color code. For an animation of trajectories and morphs of the extreme structures see Figure 10—Video 1. **Figure 10—Video 1. Structural heterogeneity of myosin-V — PCA of all atomic models.** Three-dimensional visualization of the trajectories and morphs of extreme structures along the first **(A-B)**, second **(C-D)** and third **(E-F)** principal component as identified in a PCA of all atomic models of the actomyosin-V complex. The localization of each model within the PC space is indicated for guidance. See Figure 10 for color code and an overview of PCA results.

The conformational changes mapped on each PC are readily illustrated by their corresponding trajectories as well as the extreme structures along each PC (Figure 10C, Figure 10—Video 1). The motions along the first and second PC correspond to an almost perpendicular pivoting of the U50 domain, causing a twist and shift of the central transducer β-sheet and ultimately rotations of the N-terminal and converter domain. While both PCs describe lever arm swings, the overall extent is mostly due to rearrangements along the first PC (Figure 10B-C, Figure 10—Video 1A-B). The third PC maps a rotation of the N-terminal and converter domain around the transducer, which acts as a hinge region. Since all average structures localize close to the origin of PC 3 (Figure 10A, Figure 10—Video 1E-F), we suggest that this PC accounts for an inherent flexibility of the transducer β-sheet.

The rearrangements, especially along the first PC, are reminiscent of the structural transition of myosin-V upon Mg^2+^-ADP release (Figure 3, Figure 10C, Figure 10—Video 1). In line with this, we find the strong-ADP and rigor average structures to be arranged diagonally within the PC 1–PC 2 space (Figure 10A). This indicates that the conformational heterogeneity of myosin-V as well as the isomerization associated with Mg^2+^-ADP release rely on the same principal coupling mechanism. Furthermore, this suggests that the structural transition of myosin-V along its motor cycle is driven, at least in part, by its conformational flexibility. Based on this, we therefore propose that the active site of myosin-V is not mechanically and thus rigidly coupled to the surrounding domains, in particular, the lever arm, as previously proposed (Fischer, Windshügel, Horak, Holmes, & Smith, 2005). Rather, its coupling seems to be statistical in nature, ultimately leading to a thermodynamic ensemble of conformations within each state. The associated structural flexibility of myosin-V possibly initiates transitions between structural states by giving rise to short-lived in-between conformations with favorable nucleotide binding affinities. Interactions with a nucleotide would consequently not trigger the transition, but merely stabilize myosin in its transient conformation, thereby promoting the transition to a new structural ensemble state.

A non-rigid, stochastic coupling of the active site of myosin-V is in good agreement with the release of Mg^2+^-ADP due to an isomerization and the existence of the PRT state. It also provides a good explanation for the different binding affinities of the rigor and strong-ADP state. Specifically, we propose that the extent of conformational heterogeneity tunes the binding affinity, rather than changes in the actomyosin interface since these are almost the same in all three nucleotide states studied (Figure 5). Restrictions of the conformational space by external forces, i.e. the lever arm mobility of the strong-ADP state, could furthermore account for the load-dependent delay of ADP release (Mentes et al., 2018).

While we cannot deduce from our data whether the statistical coupling and inherent conformational flexibility (Figure 9, Figure 10) is a general feature of the myosin superfamily or a hallmark of myosin-V, there is a number of independent indications for the former. First, statistical coupling of the active site has also been proposed for myosin-VI based on a recovery stroke intermediate crystal structure, showing that the lever arm can partially re-prime while the active site remains unchanged (Blanc et al., 2018). Second, conformational heterogeneity has also been reported for other myosins based on either structural (Behrmann et al., 2012; Kollmar, Dürrwang, Kliche, Manstein, & Kull, 2002; Mentes et al., 2018) or biochemical data (Klein et al., 2008). Noteworthy, a flexibility reminiscent of the one observed for myosin-V (Figure 9, Figure 10) was reported for myosin-IE in the rigor state (Behrmann et al., 2012). Conversely, no flexibility was described for myosin-IB, which adopts a single state in the absence of a nucleotide and two discrete states when bound to Mg^2+^-ADP (Mentes et al., 2018). Whether these results reflect properties of specific myosins or rather current limitations of data analysis methods, e.g., number of particles, low signal-to-noise ratio, robustness of 3D classifications (Pospich & Raunser, 2018), remains to be investigated. In general, there is little data on the conformational dynamics of myosin as most structures originate either from small cryo-EM data sets, which have an insufficient number of particles for extensive 3D classifications, or from X-ray crystallography. We therefore believe that the characterization of myosin’s energetic and dynamic landscape will provide novel insights into the details of force generation.

### Structural model of the myosin-V motor cycle

To understand the mechanism of force generation of myosin, detailed knowledge about its structural transition along the motor cycle is required. By combining our cryo-EM structures of the actomyosin-V complex with previously published crystal structures of myosin-V in the post-rigor (Coureux et al., 2004) and PPS state (Wulf et al., 2016), we were able to assemble a close-to-complete structural model of the motor cycle (Figure 11). It includes all structural states of myosin solved today, except the putative phosphate release (P_i_R) state (Llinas et al., 2015) and the intermediate recovery stroke state (Blanc et al., 2018) – hereafter referred to as intermediate state –, which have only been solved for myosin-VI. While homology models of the two states can theoretically fill the gaps in the motor cycle (Figure 11), their validity is limited to highly conserved structural elements, such as the active site, as myosin-V and myosin-VI only share ∼38 % of their sequence and furthermore walk into opposite directions along the actin filament (Wells et al., 1999).

**Figure 11.**
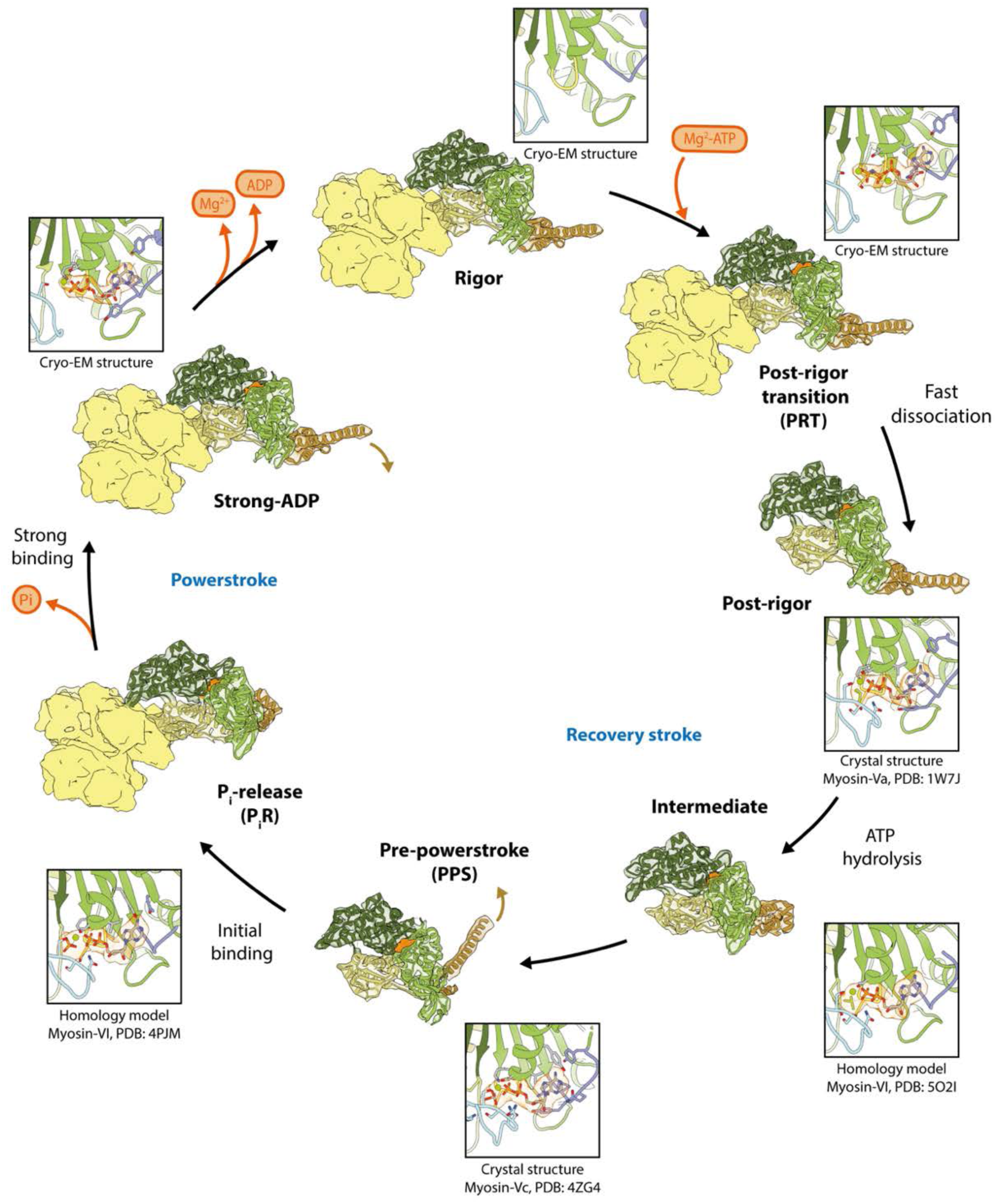
Structural schematic of the motor cycle of myosin-V. Compilation of all available structural states that have been resolved to high-resolution up to date. The strong-ADP, rigor and PRT state of myosin-Va were solved by cryo-EM within this work. Crystal structures of myosin-Va and Vc have revealed the post-rigor and PPS state, respectively (Coureux et al., 2004; Wulf et al., 2016). The putative intermediate and phosphate release (P_i_R) states are homology models based on myosin-VI crystal structures (Blanc et al., 2018; Llinas et al., 2015). Inserts show the active site of each state highlighting the nucleotide (orange) and its coordination by the P-loop, switch I, switch II and the A-loop, for color code see Figure 1. PDB accession codes are stated below each insert. Densities were simulated from atomic models. F-actin is colored in yellow and myosin is colored by domains; U50 domain (dark green), L50 domain (tan), N-terminal domain (light green) and converter domain including the lever arm (brown). See Figure 11—figure supplements 1-4 for illustrations of the structural transition along the motor cycle and Video 1 for an animation of the motor cycle.

Based on the composite structural model of the motor cycle (Figure 11), we were able to analyze the force-generating structural transition of myosin-V in detail (Video 1 and Figure 11—figure supplements 1-4). Starting from the strongly-bound rigor state, ATP would bind weakly to myosin-V, giving rise to minor local rearrangements of the active site and thereby the PRT state (Video 1, Figure 11—figure supplements 1-2). The transition to the subsequent post-rigor state is likely initiated by closure of the active site, resulting in strong nucleotide binding. The accompanying major rotation of the U50 domain leads to the opening of the actin binding cleft and thus detachment from F-actin. These changes also cause a rotation of the N-terminal domain, leading to a rearrangement of the SH1-SH2-relay helix interface and thereby to a small lever arm swing of ∼ 13° (Video 1, Figure 11—figure supplements 1-2).

Once detached, myosin rapidly transitions to the PPS state (Robert-Paganin et al., 2020). This transition involves a major rearrangement of the SH1-SH2-relay helix interface due to rotations of the U50 and N-terminal domain. Here, kinking of the relay helix allows the lever arm to perform the ∼ 81° rotation associated with the recovery stroke (Video 1, Figure 11— figure supplements 3). While it has long been assumed that ATP hydrolysis triggers these conformational changes, recent data on myosin-VI indicate that the repriming of the lever arm is driven, at least in part, by thermodynamic fluctuations (Blanc et al., 2018). ATP hydrolysis thus only eventually stabilizes the primed lever arm conformation via switch II interactions.

Although structures of initial, weakly-bound states of myosin are missing (Schröder, 2020), the partially closed actin binding cleft of the PPS state probably allows weak binding of myosin-V to F-actin (Video 1, Figure 11—figure supplements 3). Binding to F-actin in turn activates the release of inorganic phosphate from the active site of myosin, which was suggested to proceed via a second binding site (Llinas et al., 2015). In particular, it has been proposed that switch II opens a back door for P_i_ to exit and that P_i_ remains bound to a second binding site consisting of S153, T197, S203 and E461 (equivalent to S165, T211, S217 and E442 in myosin-V) until the powerstroke has completed (Llinas et al., 2015; Robert-Paganin et al., 2020) (Video 1, Figure 11—figure supplements 4). Loss of P_i_ from the active site of myosin-V, on the one hand, causes closing of the actin binding cleft via a rotation of the U50 domain (Video 1, Figure 11—figure supplements 4). The resulting conformation facilitates myosin to strongly bind to F-actin and is characteristic for the strong-ADP state (Video 1, Figure 11— figure supplements 4). On the other hand, P_i_ release is linked to rearrangements of the SH1-SH2-relay helix interface, in particular, straightening of the relay helix, which gives rise to the force-generating powerstroke (Video 1, Figure 11—figure supplements 4). Our data indicate that the powerstroke of myosin-V amounts to ∼ 77°, in contrast to a previous estimate of only 58° based on medium resolution cryo-EM data (Wulf et al., 2016).

The sequential release of Mg^2+^ and ADP completes the motor cycle. It is accompanied by a rotation of the N-terminal domain and a small lever arm swing of ∼ 9° (Video 1, Figure 11—figure supplements 4, also see Figure 3). Conversely, it does not affect the actin-binding interface (also see Figure 5), so that myosin-V remains strongly bound to F-actin in the rigor state.

Despite the complexity of the structural transitions of myosin along its motor cycle (Video 1 and Figure 11—figure supplements 1-4), a few structural elements stand out by acting as key role players. One of these is switch II, which undergoes major rearrangements during both the recovery stroke and the powerstroke and furthermore opens an exit tunnel for the P_i_. While it was previously thought that the conformation of switch II determines the position of the lever arm (Reubold et al., 2003), this view proved too simplistic in later studies (Coureux et al., 2004; Llinas et al., 2015). Instead, the lever arm orientation is likely defined by the interplay of switch II with switch I and the central transducer β-sheet. In agreement with this, we observe a small lever arm swing upon ADP release which is not associated to a movement of switch II (Figure 3, Video 1). The interplay of switch I and II probably also tunes the opening and closing of the actin binding cleft, explaining why strong binding of a triphosphate nucleotide and to F-actin are reciprocal (Coureux et al., 2004).

Another key structural element is the central transducer β-sheet. It connects all domains to the active site and undergoes various deformations, mostly twisting and shifting, when the active site rearranges during the myosin motor cycle (Figure 3, Figures 9-10, Video 1). While coupling to the actin interface has already been suggested previously (Coureux et al., 2003), our data indicate that the transducer also transmits changes to the N-terminal domain and thus to the converter domain (Figure 3, Figures 9-10). The centrality of the transducer is further supported by the suggestion that neighboring regulatory elements, such as loop 1 (aa 184-191), alter the kinetic constants of myosin by tuning the plasticity of the transducer β-sheet (Coureux et al., 2004; Sweeney et al., 1998).

The last key structural element is the relay helix. It is part of the SH1-SH2-relay helix interface that ultimately determines the orientation of the converter domain and thus the lever arm (Video 1 and Figure 11—figure supplements 1-4). Interestingly, we find two alternative mechanisms by which this interface is remodeled during the motor cycle. On the one hand, the relay helix kinks or straightens due to changes of switch I and switch II, resulting in the recovery stroke and powerstroke, respectively (Video 1, Figure 11—figure supplements 3-4). Rotations of the N-terminal domain, on the other hand, remodel the interface when myosin transitions to the rigor and post-rigor state, respectively (Video 1, Figure 11—figure supplements 2,4).

Our results finally not only indicate two alternative mechanisms that control the lever arm position, but also two complementary types of allosteric coupling for the active site and the actin-binding cleft of myosin-V, respectively. The pronounced structural heterogeneity of myosin-V, in particular the flexibility of the lever arm, suggests a non-rigid, statistical coupling of the active site. Conversely, our data show no variation of the actin binding cleft in all three nucleotide states (Figure 9). Based on this observation and the marked effects that binding to and detachment from F-actin has on the overall structure of myosin-V (Video 1), we propose a rigid mechanical coupling of the actin binding site to the periphery. In this way, F-actin binding acts as an “on/off“ switch for strong ATP/ADP-P_i_ binding, ultimately facilitating the rectification of force production.

### Summary

The presented high-resolution cryo-EM structures of the actomyosin-V complex in three nucleotide states – nucleotide-free, Mg^2+^-ADP and Mg^2+^-AppNHp – (Table 1) provide unprecedented insights into the structural basis of force generation. First, a comparison of the strong-ADP (Figure 1) and rigor state (Figure 2) has revealed the structural transition of myosin-V upon Mg^2+^-ADP-release (Figure 3), which is reminiscent of the one of myosin-IB (Mentes et al., 2018) and yet differs in its details. Second, the structure of Mg^2+^-AppNHp-bound myosin-V has uncovered a previously unseen post-rigor transition (PRT) state (Figure 4). Myosin in the PRT state is strongly bound to F-actin and adopts a conformation resembling the rigor state. Because of the weak binding to the active site, AppNHp, and probably ATP, does not directly trigger the detachment from F-actin and thus the transition to the post-rigor state. Instead, strong nucleotide binding likely needs to be established to eventually initiate detachement. Since a coordination similar to Mg^2+^-AppNHp has been reported for Mg^2+^-ADP in myosin-IB (Mentes et al., 2018), weak nucleotide binding might not be restricted to AppNHp, but represents a commonality of all transition states that are strongly-bound to F-actin.

Interestingly, and despite the differences in the F-actin binding affinity, we find the actin binding interface basically indistinguishable in all three nucleotide states (Figure 5), suggesting that strongly-bound states utilize a common binding scheme. Furthermore, a comparison of the interface with the one of other myosins has revealed specific features of the myosin-V interface and indicates that a strong Milligan contact (Milligan et al., 1990) is characteristic of myosins with high duty-ratios.

In contrast to previous reports (Wulf et al., 2016), our results elucidate that myosin-V hardly alters the structure of aged F-actin-PHD (Figure 6). Conversely, it has a remarkable effect on the structure of young F-actin-JASP, specifically selecting the closed D-loop state (Figure 7 and Figure 8 ) and thereby overriding the “rejuvenating effect” of JASP (Merino et al., 2018; Pospich et al., 2020). Whilst this result does not reveal the structural basis of myosin-V’s nucleotide sensitivity (Zimmermann et al., 2015), it has possible implications for the directed remodeling of the actin cytoskeleton (Merino et al., 2019) and offers an explanation for pyrene fluorescence quenching upon myosin binding (Kouyama & Mihashi, 1981).

Additional heterogeneity analysis of our data revealed a pronounced structural flexibility of myosin-V (Figures 9 and 10), indicating a non-rigid, stochastic coupling of the active site. Structural transitions of myosin-V are hence likely not initiated by binding of a specific nucleotide, but rather by thermodynamic fluctuations, as previously suggested for myosin-VI (Blanc et al., 2018). Changes of the actin binding cleft associated with binding to and detachment from F-actin, conversely, are coupled rigidly to the overall conformation of myosin-V. In this way, F-actin can effectively promote and rectify the force generation mechanism.

Finally, our high-resolution structures have enabled the assembly of what is, to our knowledge, the currently most complete structural model of the myosin motor cycle (Figure 11) as well as the detailed description of the associated structural transition of myosin-V (Video 1). Based on this, we not only have identified structural key role players, but also two complementary mechanisms by which the lever arm orientation is tuned.

Taken together, we have elucidated many, previously unknown details of the force generation mechanism. The general validity of these results, i.e., if they are limited to myosin-V or hold for the complete myosin superfamily, as well as the possible implications of our findings have to be thoroughly tested in future studies. Structural data on how actin activates myosin and how myosin eventually detaches will surely be of special interest (Robert-Paganin et al., 2020; Schröder, 2020; Sweeney et al., 2020). Yet, great insights could also come from a characterization of the energetic and dynamic landscape of myosin. Finally, unraveling of the structural basis of nucleotide sensitivity (Zimmermann et al., 2015) will further promote our understanding of the regulation of both myosin and the actin cytoskeleton (Merino et al., 2019).

## Material and methods

### Key Resources Table

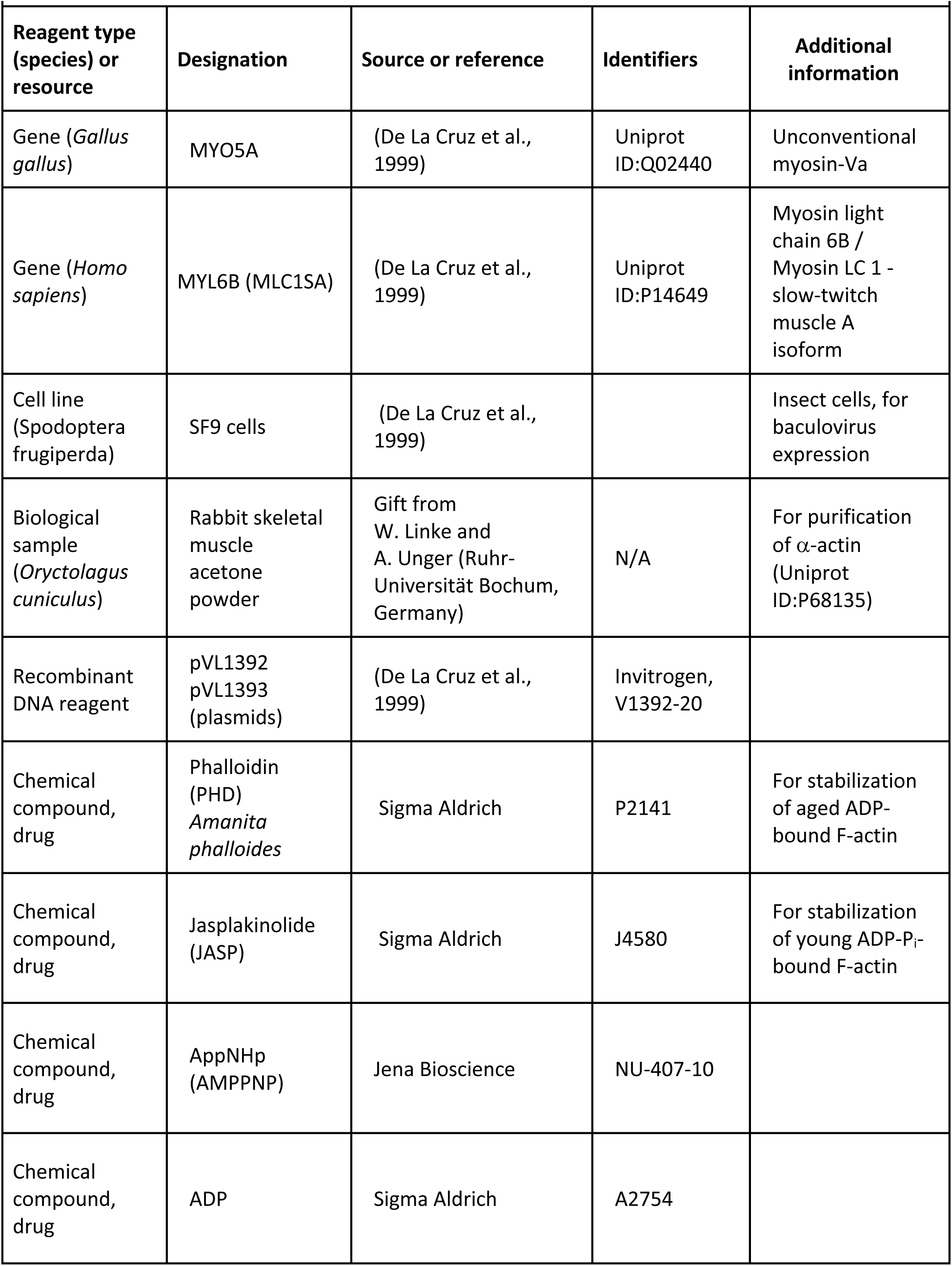
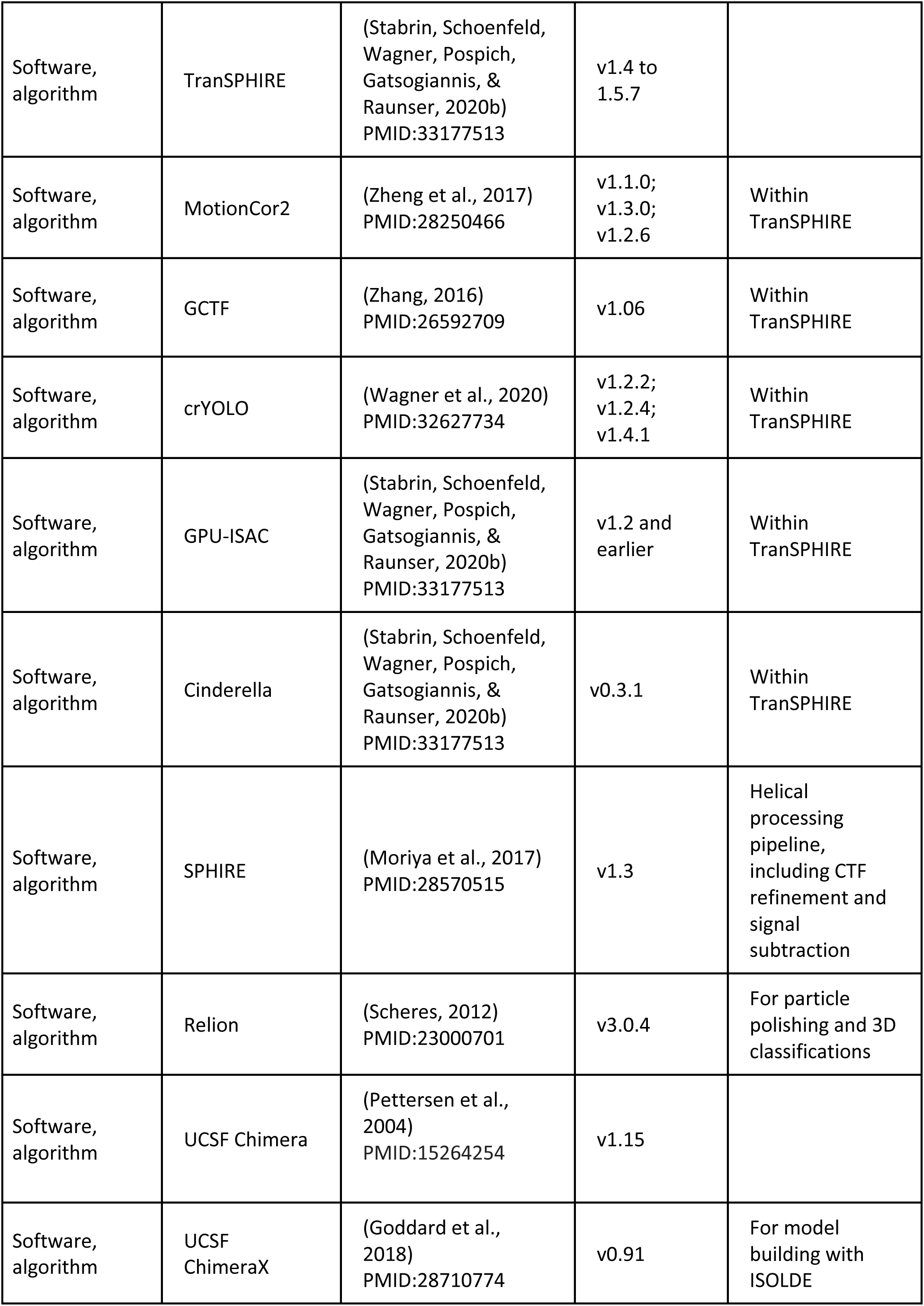
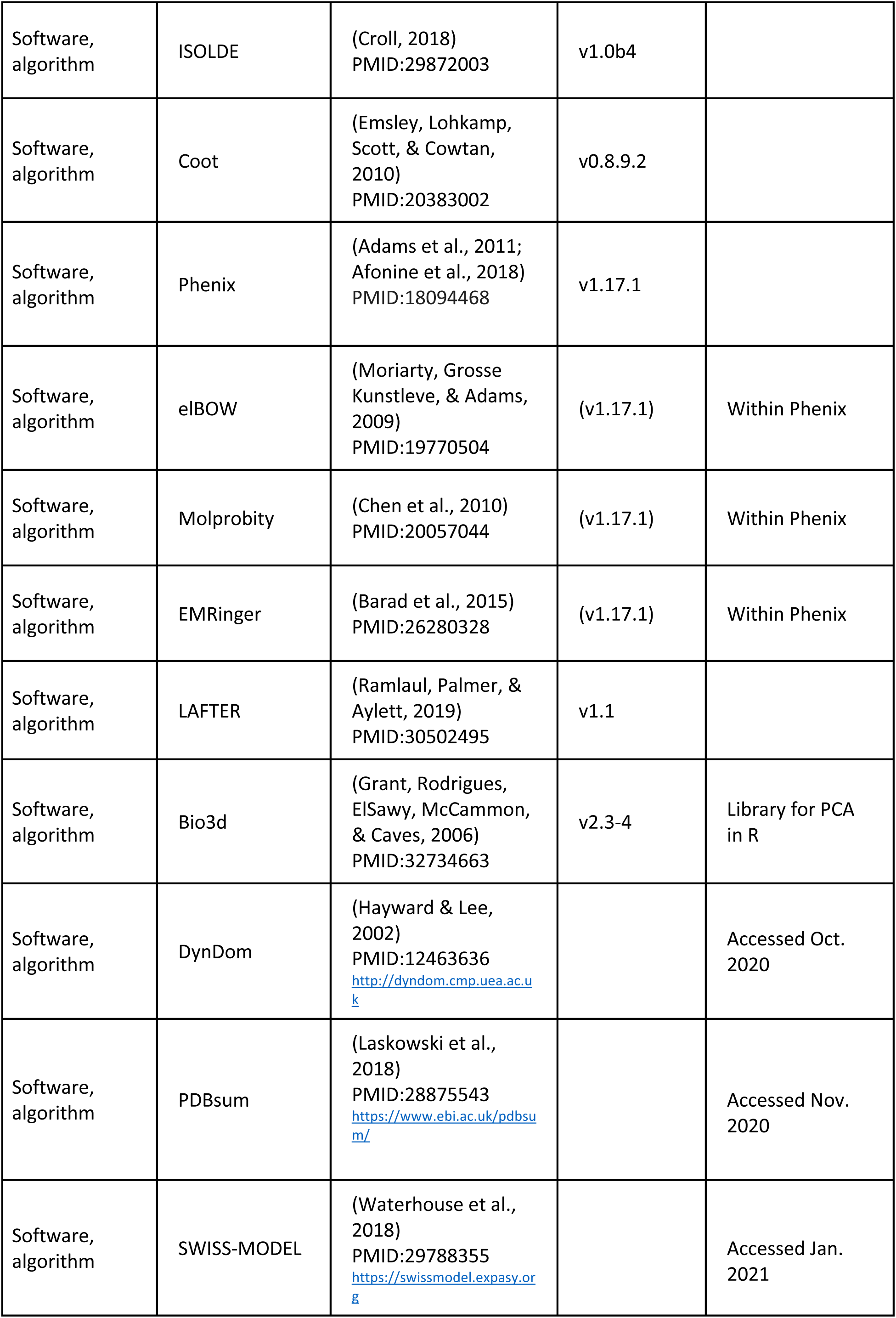
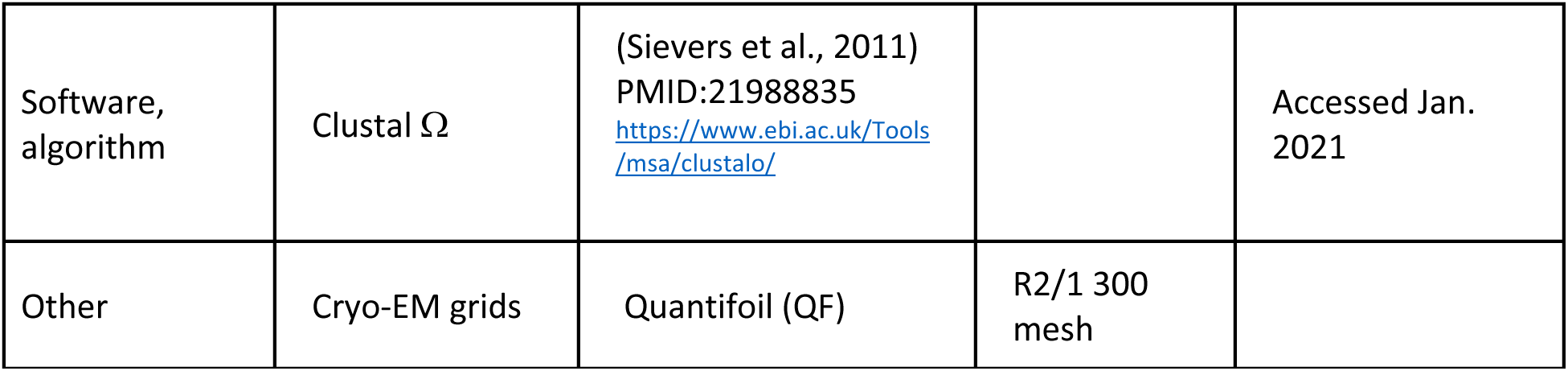

### Protein expression and purification

Actin was purified from rabbit skeletal muscle acetone powder by cycles of polymerization and depolymerization as described previously (Merino et al., 2018; Pardee & Spudich, 1982; Pospich et al., 2020). Purified G-actin was flash-frozen and stored in G-actin buffer (5 mM Tris pH 7.5, 1 mM DTT, 0.2 mM CaCl_2_, 2 mM NaN_3_ and 0.5 mM ATP) at -80 °C.

Myosin V was expressed using the baculovirus/SF9 cell expression system. To create the recombinant virus used for expression, the cDNA coding for chicken myosin Va was truncated after the codon corresponding to Arg792. This construct encompassed the motor domain and the first light chain/calmodulin-binding site of myosin Va. A “Flag” tag DNA sequence (encoding GDYKDDDDK) (Hopp et al., 1988) was appended to the truncated myosin V coding sequence to facilitate purification. A truncated cDNA for the LC1-sa light chain (De La Cruz, Wells, Sweeney, & Ostap, 2000) was co-expressed with the truncated myosin V heavy chain in SF9 cells as described in (De La Cruz et al., 1999). The cells were grown for 72 hr in medium containing 0.2 mg/ml biotin, harvested and lysed by sonication in 10 mM imidazole, pH 7.4, 0.2 M NaCl, 1 mM EGTA, 5 mM MgCl_2_, 7% (w/v) sucrose, 2 mM DTT, 0.5 mM 4-(2-aminoethyl)benzenesuflonyl fluoride, 5 μg/ml leupeptin and 2 mM MgATP. An additional 2 mM MgATP was added prior to a clarifying spin at 200,000 × g for 40 min. The supernatant was purified using FLAG-affinity chromatography (Sigma). The column was washed with 10 mM imidazole pH 7.4, 0.2 M NaCl and 1 mM EGTA and the myosin eluted from the column using the same buffer plus 0.1 mg/ml FLAG peptide. The fractions containing myosin were pooled and concentrated using an Amicon centrifugal filter device (Millipore) and dialyzed overnight against F-actin buffer (10 mM HEPES pH 7,5, 100 mM KCl, 2mM MgCl_2_, 1mM DTT and 1mM NaN_3_). Purified myosin-V-LC was flash-frozen and stored at -80 °C

### Sample preparation for cryo-EM

Aliquots of G-actin were freshly thawed and cleared by ultracentrifugation (Beckmann Rotors, TLA 120.1, 100.000 g, 1 h, 4 °C). The concentration of G-actin was measured by absorption spectroscopy (Spectrophotometer DS-11, DeNovix, E_290 nm_ ≈ 22,000 M^-1^ cm^-1^ at 290 nm (Hertzog & Carlier, 2005)). Polymerization was induced by adding 100 mM KCl, 2 mM MgCl_2_ and 0.5 mM ATP. In case of young JASP-stabilized F-actin, actin was polymerized in presence of a 2x molar excess of JASP (Jasplakinolide, Sigma Aldrich, freshly solved in DMSO, 1 mM stock). After 2 h of incubation at room temperature, the sample was transferred to 4 °C for further polymerization overnight. Filaments were collected by ultracentrifugation (Beckmann Rotors, TLA 120.1, 100.000 g, 2 h, 4 °C) and pellets rinsed and resuspended in F-actin buffer (10 mM HEPES pH 7.5, 100 mM KCl, 2 mM MgCl_2_, 1 mM DTT, 1 mM NaN_3_) supplemented with 0.02 w/v % tween 20 (to improve spreading of the sample droplet on the cryo-EM grid). No additional ADP or JASP was added. In case of aged PHD-stabilized F-actin, a 2x molar excess of PHD (Phalloidin, Sigma Aldrich, freshly solved in methanol, 1.25 mM stock) was added to resuspended filaments, which have aged, i.e. hydrolyzed ATP and released the inorganic phosphate, during the overnight polymerization step. Filaments were stored at 4 °C for a few hours before preparation of cryo-EM grids.

Aliquots of myosin-V-LC were freshly thawed, diluted 1:1 with F-actin buffer and cleared by centrifugation (Eppendorf centrifuge 5424R, 21,000 g, 5 min, 4 °C). The concentration was determined by absorption spectroscopy (Spectrophotometer DS-11, DeNovix, E_280 nm_ ≈ 106,580 M^-1^ cm^-1^ at 280 nm).

### Cryo-EM grid preparation and screening

To avoid bundling of actomyosin filaments, F-actin was decorated with myosin-V-LC on the grid, as described previously (von der Ecken et al., 2016). A freshly glow-discharged holey-carbon grid (QF R2/1 300 mesh, Quantifoil) was mounted to a Vitrobot cryoplunger (Thermo Fisher). 3 µl of F-actin (3-4 µM) were applied onto the front of the grid and incubated for 60 s. Excess solution was manually blotted from the side using blotting paper (Whatman No. 4). Immediately, 3 µl of myosin-V-LC (3-13 µM) were applied onto the grid and incubated for 30 s. The grid was automatically blotted for 9 s (blot force -15 or -25, drain time 0-1 s) and plunged into liquid ethane. The temperature was set to 13 °C for all samples but the AppNHp sample, where either 4 °C or 25 °C were used (two settings and data sets, see Table 1).

Myosin was kept in F-actin buffer and was only diluted and supplemented with a nucleotide and tween 20 immediately before application to the grid to avoid any adverse effects. When preparing the strong-ADP state, myosin was diluted 1:1 in a 2x ADP buffer (F-actin buffer with 40 mM MgCl_2_, 4 mM ADP and 0.04 w/v % tween 20). For the rigor samples, myosin was diluted in F-actin buffer and supplemented with 0.02 w/v % tween 20. AppNHp-bound samples were prepared in analogy to rigor samples, but additional 5 mM AppNHp and 4 mM MgCl_2_ were added. As AppNHp hydrolyzes spontaneously, only freshly solved (10 mM HEPES pH 8.0, 1 mM DTT, 1 mM NaN_3_ and 2 mM MgCl_2_) or recently frozen AppNHp was used. Ion-pair reversed-phase chromatography experiments using freshly solved AppNHp indicated a purity of ≥ 98 %, with 1.5 % AppNH_2_ (hydrolysis product) and no preferential binding of AppNH_2_ to myosin. Thus, AppNH_2_ does not get enriched in the active site of myosin-V, as it is the case for F-actin (Cooke & Murdoch, 1973). To increase the binding affinity of AppNHp-bound myosin to F-actin (Konrad & Goody, 1982), the concentration of potassium chloride in the myosin sample buffer was reduced to 10-13 mM KCl by dilution with F-actin buffer without KCl. F-actin samples were diluted using F-actin buffer supplemented with 0.02 w/v % tween 20. After dilution to the final concentration, the PHD-stabilized F-actin samples contained 0.4-0.9 % methanol.

Protein concentrations were adjusted empirically based on the overall concentration on the grid and decoration of actin filaments. The concentration of myosin required to saturate F-actin (3-4 µM) strongly depended on the nucleotide state; while 3-4 µM myosin were sufficient in case of the rigor and strong-ADP state, 10-13 µM myosin were required for the AppNHp sample, even though the salt concentration of the buffer was lowered to increase the binding affinity.

Grids were screened on a Talos Arctica microscope (Thermo Fisher) operated at 200 kV and equipped with a Falcon III direct detector (Thermo Fisher).

In total, six different samples were plunged, screened and imaged; also see Table 1. On the one hand, aged PHD-stabilized F-actin was decorated with myosin-V-LC in three different nucleotide states, i.e. in absence of a nucleotide (rigor state) and bound to either Mg^2+^-ADP or Mg^2+^-AppNHp (aged rigor, ADP and AppNHp). For the AppNHp-bound sample, two data sets were collected from grids that were plunged using different incubation temperatures, i.e. 4 °C or 25 °C. On the other hand, young JASP-stabilized F-actin was imaged on its own and in complex with myosin-V-LC in the rigor state (young F-actin and rigor). The corresponding grids were prepared in one plunging session, i.e. within a short time frame of 1-2 h, using the same JASP-stabilized F-actin sample.

### Cryo-EM data acquisition

Data sets were acquired on Titan Krios microscopes (FEI Thermo Fisher) operated at 300 kV and equipped with a X-FEG using EPU. Specifically, data sets of the rigor and strong-ADP state were acquired on a standard Krios (Cs 2.7 mm, pixel size 1.06 Å), while a Cs-corrected Krios (pixel size 1.10 Å) was used for the remaining data sets. Equally dosed frames were collected using a K2 Summit (super-resolution mode, Gatan) direct electron detector in combination with a GIF quantum-energy filter (Bioquantum, Gatan) set to a slit width of 20 eV. For every hole four micrographs consisting of 40 frames were collected close to the carbon edge, resulting in a total electron dose of ∼79-82 eÅ^-2^ within an exposure time of 15 s. The defocus was varied within a range of ∼0.4-3.2 µm. Acquisition details of all six data sets (aged rigor, ADP and AppNHp 4 °C + 25 °C as well as young F-actin and rigor) including pixel size, electron dose, defocus range and the total number of images collected are summarized in Table 1. Data acquisitions were monitored and evaluated live using TranSPHIRE (Stabrin, Schoenfeld, Wagner, Pospich, Gatsogiannis, & Raunser, 2020a).

### Cryo-EM data processing

Data sets were automatically pre-processed on-the-fly during the data acquisition using TranSPHIRE (Stabrin, Schoenfeld, Wagner, Pospich, Gatsogiannis, & Raunser, 2020a). Pre-processing included drift correction and dose weighting by MotionCor2 (Zheng et al., 2017), CTF estimation using GCTF (Zhang, 2016) and particle picking with crYOLO (Wagner et al., 2020; 2019) (filament mode, box distance 26-27 px equivalent to one rise of `∼`27.5 Å, minimum number of boxes 6) for all data sets. The latest version of TranSPHIRE (Stabrin, Schoenfeld, Wagner, Pospich, Gatsogiannis, & Raunser, 2020b), which was used for the processing of the AppNHp data sets, also supported automatic, on-the-fly particle extraction (box size 320 px, filament width 200 px) as well as batch-wise 2D classification (batch size 13k, filament width 200 px, radius 150 px, 60-100 particles per class), 2D class selection and 3D refinement using software of the SPHIRE package (Moriya et al., 2017). In particular, a GPU-accelerated version of ISAC (Stabrin, Schoenfeld, Wagner, Pospich, Gatsogiannis, & Raunser, 2020a; Yang, Fang, Chittuluru, Asturias, & Penczek, 2012) and the deep-learning 2D class selection tool Cinderella (Wagner, 2019) were used. For all other data sets, particles were extracted and 2D classified after data collection using analogous settings and helical SPHIRE 1.3 (Moriya et al., 2017; Stabrin, Schoenfeld, Wagner, Pospich, Gatsogiannis, & Raunser, 2020a). Particles that were not accounted during the initial, batch-wise 2D classification, e.g. because they represent rare views, were merged and inputted to another round of 2D classification until no more stable classes were found. All micrographs were assessed manually and images sorted based on ice and protein quality, resulting in a removal of 6-36% of the data sets, see Table 1 for details. Particles contributing to classes found ‘good’ by either Cinderella or manual inspection and belonging to micrographs of good quality were written to virtual particle stacks for further processing in 3D.

As an initial 3D refinement and 3D classification revealed no differences in the overall structure of myosin in the two AppNHp data sets, i.e. plunged at 4 °C and 25 °C, corresponding particles were merged for further processing. The final number of particles ranged from 212,660 (young JASP-stabilized F-actin) to 2,446,218 (combined AppNHp data sets), see Table 1 —Supplementary Tables 1-3 and Figure 7—Table 1 for details. A concise overview of all key processing steps including the number of particles and nominal resolutions can be found in Table 1—Supplementary Figure 1.

All data sets were processed using the helical refinement program sp_meridien_alpha.py implemented in SPHIRE 1.3 (Moriya et al., 2017; Stabrin, Schoenfeld, Wagner, Pospich, Gatsogiannis, & Raunser, 2020a). In contrast to other helical refinement routines, SPHIRE does not refine or apply any helical symmetry, and thereby avoids possible symmetrization pitfalls. Instead, the software offers the usage of constraints tailored to helical specimen, e.g. on the tilt angle and shift along the filament, to guide the refinement (also see Methods of (Pospich et al., 2021)). For all 3D refinements the tilt angle was softly restrained to the equator during exhaustive searches (--theta_min 90 --theta_max 90 --howmany 10). The shift along the filament axis was furthermore limited to ± half of the rise (--helical_rise 27.5) to avoid shifts larger than one subunit. Finally, the smear (number of views considered for the reprojection of each particle) was reduced to a combined weight of 90 % (--ccfpercentage 90). An initial 3D reference was created from the atomic model of a previously published actomyosin complex in the rigor state (PDB:5JLH, without tropomyosin, (von der Ecken et al., 2016)) and filtered to 25 Å using EMAN2 (Tang et al., 2007) and SPHIRE (Moriya et al., 2017). For the initial 3D refinement a sampling angle of 3.7°, filament width of 120 px and a radius of 144 px (45 % of the box size), but no 3D mask were used. Based on the resulting 3D density map, a wide mask covering the central 85 % of the filament was created. This map and mask where then used to run a fresh, global 3D refinement using the same settings as before. Based on the results of this refinement, particles were CTF refined within SPHIRE (Moriya et al., 2017) providing the nominal resolution according to the FSC_0.143_-criterion. CTF refined particles were locally 3D refined using the final map of the previous 3D refinement filtered to 4 Å as reference. The fine angular sampling typically used in local refinements makes helical restraints superfluous, as projections parameters can only locally relax anyways. For this reason, particles were locally refined using the non-helical 3D refinement program sp_meridien.py in combination with a sampling angle of 0.9°, a shift range of 2 px and a shift step size of 0.5 px. In case of the young F-actin and young rigor data sets, the resolution could be further improved by particle polishing in Relion 3.0.4 (Scheres, 2012; Zivanov et al., 2018). For this purpose, refinement results were converted to Relion star format using sp_sphire2relion.py. Meta data of the initial motion correction step required for polishing were automatically created by TranSPHIRE and were directly provided. Polished particles were transferred back to SPHIRE and passed through another round of local 3D refinement using the same settings as before.

To focus the refinement on the central part of the filament, a wide mask containing the central three actin and central two myosin-V-LC subunits including all ligands (sub volume referred to as central 3er/2er map) was created and applied in a subsequent local 3D refinement. Post refinement of the resulting half maps using a central 3er/2er mask yielded maps with average resolutions ranging from 2.9 – 3.2 Å according to the FSC_0.143_-criterion, see Table 1—Supplementary Figure 1, Table 1 —Supplementary Tables 1-3 and Figure 7—Table 1 for details.

With the aim to further improve the density of myosin, the signal of all subunits but the central actomyosin subunit (sub volume referred to as central 1er map) was subtracted from the 2D particle images within SPHIRE 1.3 (Moriya et al., 2017). Particles were additionally recentered to bring the center of mass close to the center of the box. Signal-subtracted particles were subjected to another round of local 3D refinement applying a central 1er mask and filtering the centered reference map to 3.5 Å. Although post refinement of the resulting half maps using a central 1er mask did not yield density maps of higher nominal resolution, the map quality of especially myosin could be significantly improved, see Table 1—Supplementary Figures 1-2, Table 1 —Supplementary Tables 1-3, Figure 7—Table 1 and Figure 7—Supplementary Figure 2 for details.

The anisotropic quality of the final central 1er maps suggested structural heterogeneity within myosin. For this reason, signal-subtracted particles and corresponding projection parameters were transferred to and 3D classified in Relion 3.0.4 (Scheres, 2012). As domain movement were assumed to be small and to reduce the risk of overrefinement, 3D alignment was deactivated (--skip_align) and the resolution strictly limited to 8 Å (--strict_highres_exp 8). The final central 1er map filtered to 15 Å was inputted as a reference, while a corresponding wide mask was applied and solvent flattening and CTF correction activated. The regularization parameter T and number of classes K was empirically adjusted. While a parameter of T=40 (--tau2_fudge 40) proved well suited for all data sets, finding a suitable number of classes posed a challenge. Running multiple 3D classifications with different numbers of classes resulted in classes of various, related structural states with little overlap, i.e. classes of different runs could not be matched as they did generally not superimpose. The same was true, when rerunning a 3D classification job using the same settings but a different seed. These results suggest a continuous structural heterogeneity of myosin, in contrast to several discrete states. While software tailored to the characterization of cryo-EM data exhibiting continuous structural states has recently been published (Zhong, Bepler, Berger, & Davis, 2021), it proved unsuitable for the processing of signal-subtracted actomyosin filaments due to the need of 3D masking. To characterize the structural heterogeneity of myosin-V-LC by standard 3D classification in Relion 3.0.4 (Scheres, 2012) as good as possible, the number of 3D classes was optimized experimentally to yield the highest number of classes with a resolution and map quality sufficient for atomic modelling (≤ 3.7 Å). To do so, multiple 3D classifications with varying number of classes, e.g. from 2 to 12, were performed and particles split into subsets according to the classification results. Subsets were then transferred to SPHIRE and individually subjected to a local 3D refinement from stack (no reference required, same settings as before). Eventually, each subset was post-refined and the resulting map manually assessed. In the end, the 3D classification which yielded the most maps of high quality was chosen. In this way, a total of 18 high-resolution maps (referred to as 3D class averages or 3D classes) were achieved for the four actomyosin data sets. Corresponding subsets contained 81,757 to 365,722 particles, see Table 1 —Supplementary Table 1-3 and Figure 7— Table 1 for details. An overview of all refined maps, associated resolutions and the underlying number of particles is given in Table 1—Supplementary Figure 1.

To ease the interpretation of maps as well as model building, all final maps, i.e. central 3er/2er, central 1er and 3D class averages, were additionally filtered to local resolution using SPHIRE 1.3 (Moriya et al., 2017) and denoised using LAFTER (Ramlaul et al., 2019).

### Model building, refinement and validation

Previous cryo-EM structures of PHD-stabilized aged F-actin (PDB: 6T20, (Pospich et al., 2020)) and JASP-stabilized young F-actin (PDB: 5OOD, (Merino et al., 2018)) were used as starting models for F-actin in the rigor actomyosin complexes (aged and young rigor). The models of PHD and JASP were replaced by single-residue initial models generated from SMILES strings by elBow (Moriarty et al., 2009) within Phenix (Adams et al., 2011) using the --amber option. The corresponding cif constraints libraries were used for all further refinements. A rigor-like crystal structure of the myosin-V-LC complex (PDB: 1OE9, (Coureux et al., 2003)) was used as an initial model for myosin and the bound light chain within the aged rigor structure. Stubs were replaced by full residues and residues which are missing in the crystal structure, but are resolved in the cryo-EM density map, were added manually in Coot (Debreczeni & Emsley, 2012; Emsley et al., 2010). For all other models, i.e. of the ADP, AppNHp and young rigor state, the final refined model of the PHD-stabilized rigor actomyosin complex was used as starting model. Initial models of ADP and AppNHp are based on previous cryo-EM and crystal structures of myosin (PDB:6C1D, (Mentes et al., 2018) and PDB: 1MMN, (Gulick et al., 1997)). Starting models were rigid-body fitted into the density map using UCSF Chimera (Pettersen et al., 2004) and ligands were coarsely refined in Coot (Debreczeni & Emsley, 2012; Emsley et al., 2010) prior to model building.

Atomic models of the central actomyosin subunit, consisting of one F-actin, myosin, LC and PHD/JASP molecule (central 1er), were refined using ISOLDE (Croll, 2018) within UCSF ChimeraX (Goddard et al., 2018). For this purpose, hydrogens were added to the starting model using the addh command in UCSF Chimera (Pettersen et al., 2004) and manually adjusted when necessary. Custom residue definitions for PHD and JASP were created based on the elBOW output within the ISOLDE shell. To reliably model both high- and medium-resolution features, several maps, e.g. filtered to nominal or local resolution and sharpened by different B-factors, were loaded to ISOLDE. Maps filtered by LAFTER (Ramlaul et al., 2019) were also loaded for visual guidance, but excluded from the refinement (weight set to 0, MDFF deactivated). All density maps were segmented based on the starting model using the color zone tool within UCSF Chimera (Pettersen et al., 2004) to exclude density not corresponding to the central actomyosin subunit.

Each refinement in ISOLDE was started with a 2-3 min all atom simulation to reduce the overall energy of the system. Afterwards, overlapping stretches of the protein and atoms within close vicinity were successively adjusted and refined. When necessary rotamer and secondary structure restraints were introduced. After passing through the complete protein complex once, the quality of the model was assessed using the metrics provided by ISOLDE, i.e. Ramachandran plot, rotamer outlier and clash score, and outliers were locally addressed. Residues not resolved by the electron density map, e.g., due to flexibility, were not included in the respective atomic model, while incompletely resolved side chains were set to most likely rotamers.

The density corresponding to the light chain was of insufficient quality for reliable model building. Hence, the model of the light chain was kept fixed during refinements in ISOLDE. Afterwards, the reference crystal structure (PDB 1OE9, (Coureux et al., 2003)) was rotamer-optimized in Coot and rigid body fitted into the density using UCSF Chimera.

Finally, atomic models were real-space refined in Phenix (Adams et al., 2011; Afonine et al., 2018) against a sharpened density map filtered to nominal resolution (FSC_0.143_). To only relax and validate the model but prohibit large changes, local grid search, rotamer and Ramachandran restraints were deactivated and the starting model was used as a reference. Furthermore, NCS and secondary structure restraints were applied and cif libraries provided for PHD and JASP.

Only models of the central actomyosin subunit (central 1er) were built in ISOLDE. Atomic models of subsets, i.e. 3D class averages, were built starting from the average, all-particle model and the corresponding ISOLDE/UCSF ChimeraX session including restraints. Whereas average models of different states, i.e. rigor, ADP, AppNHp, were built within new sessions to avoid any bias. Atomic models consisting of three actin and two myosin-LC subunits (central 3er/2er) were assembled from the models of the monomeric complex (central 1er) by rigid-body fitting in UCSF Chimera. The filament interface was manually inspected in Coot and side chain orientations adjusted when necessary. Finally, the multimeric model was real-space refinement in Phenix.

After real-space refinement, the residue assignment of PHD was changed from a single non-standard residue to a hepta-peptide consisting of TRP-EEP-ALA-DTH-CYS-HYP-ALA. All atomic models were assessed and validated using model-map agreement (FSC, CC), MolProbity (Chen et al., 2010) and EMRinger (Barad et al., 2015) statistics.

In total, 27 atomic models were build based on density maps with a resolution ranging from 2.9 Å to 3.7 Å; models include five central 1er and four central 3er/2er all-particle models as well as 18 models representing subsets identified by 3D classification (see Table 1 — Supplementary Table 1-3 and Figure 7—Table 1 for details).

### Homology modeling

For the structural analysis of the complete myosin motor cycle, structure-based homology models of the PiR (phosphate release state, based on a crystal structure of myosin-VI, PDB: 4PJM, (Llinas et al., 2015)) and intermediate recovery stroke state (based on a crystal structure of myosin-VI, PDB: 5O2L, (Blanc et al., 2018)) were created using SWISS-MODEL (Waterhouse et al., 2018) and a sequence alignment generated by Clustal Ω (Sievers et al., 2011). While the overall conformation of the homology models is likely not meaningful, not least due to the reverse directionality of myosin-VI compared to myosin-V, the organization of the active site is likely representative.

### Structural analysis and visualization

Figures and movies were created with UCSF Chimera (Pettersen et al., 2004) and modified using image or movie processing software when required.

For the visualization of myosin and the actomyosin interface central 1er (central actyomyosin subunit) and central 3er/2er (central three F-actin and two myosin molecules) models and maps are shown, respectively, as they include all important contact sites and are best resolved. Models protonated by H++ (Anandakrishnan, Aguilar, & Onufriev, 2012) at pH 7.5 with HIC replaced by HIS were used for all surface representations. To optimally visualize features of different local resolution, a variety of maps is displayed within figures and movies (also see legends). Specifically, LAFTER maps are used to visualize the complete actomyosin structure and features of lower resolution, while post-refined maps are shown in close-up views, e.g of the active site.

For the structural comparison of different structural states or isoforms of myosin, the respective atomic models were superimposed on the HR (aa 493-514), HS (aa 520-531) and HW helices (aa 635-653) within the HLH motif (L50 domain), which is structurally highly conserved in all states strongly bound to F-actin. While overall structural changes are visualized well using this reference, the partially large domain movements make a comparison of the active sites challenging. For this reason, models were imposed on the HF helix (aa 169-190, trailing the P-loop) when comparing the active sites instead.

Relative rotation angles of the lever arm were computed as angles between axes created for the corresponding helix in Chimera (Pettersen et al., 2004) using default settings.

Protein-protein and protein-ligand interactions were analyzed with PDBsum (Laskowski, 2009). Conformational changes and structural heterogeneity of the central 1er models were characterized by principal component analysis (PCA) using the Bio3d library (Grant et al., 2006) in R (R Core Team, 2017). Models were superimposed on an automatically determined structural stable core, which encompasses almost the complete F-actin subunit and parts of the HLH-motif in the L50 domain. Gaps within the sequence and ligands were excluded. Data points were manually grouped and colored based on the underlying data set and type of model, i.e. average model vs. 3D class average. For the direct visualization of PCA results, trajectories along each principle component were exported and morphed in UCSF Chimera (Pettersen et al., 2004). Mobile domains within myosin (central 1er, chain A) and their motion were identified and analyzed using DynDom (Hayward & Lee, 2002)

## Data availability

The atomic models and cryo-EM maps are available in the PDB (Burley et al., 2018) and EMDB databases (Lawson et al., 2011), under following accession numbers: aged PHD-stabilized actomyosin-V in the strong-ADP state: 7PM5, EMD-13521 (central 1er), 7PM6, EMD-13522 (central 3er/2er), 7PM7, EMD-13523 (class 2), 7PM8, EMD-13524 (class 3), 7PM9, EMD-13525 (class 4), 7PMA, EMD-13526 (class 5), 7PMB, EMD-13527 (class 6), 7PMC, EMD-13528 (class 7) ; aged PHD-stabilized actomyosin-V in the rigor state: 7PLT, EMD-13501 (central 1er), 7PLU, EMD-13502 (central 3er/2er), 7PLV, EMD-13503 (class 1), 7PLW, EMD-13504 (class 3) and 7PLX, EMD-13505 (class 4); aged PHD-stabilized actomyosin-V in the PRT state: 7PMD, EMD-13529 (central 1er), 7PME, EMD-13530 (central 3er/2er), 7PMF, EMD-13531 (class 1), 7PMG, EMD-13532 (class 3), 7PMH, EMD-13533 (class 4), 7PMI, EMD-13535 (class 5), 7PMJ, EMD-13536 (class 6), 7PML, EMD-13538 (class 8); young JASP-stabilized actomyosin-V in the rigor state: 7PLY, EMD-13506 (central 1er), 7PLZ, EMD-13507 (central 3er/2er), 7PM0, EMD-13508 (class 1), 7PM1, EMD-13509 (class 2), 7PM2, EMD-13510 (class 4); and young JASP-stabilized F-actin: 7PM3, EMD-13511. The datasets generated during the current study are available from the corresponding author upon reasonable request.

## Author contributions

Conceptualization: S.R., A.H., and H.L.S. Investigation and Formal analysis: S.P. Resources: H.L.S and A.H. Visualization and writing – Original Draft: S.P. Writing – Review & Editing: S.P., S.R, A.H. and H.L.S. Supervision and funding acquisition: S.R.

## Acknowledgements

We thank E. Sirikia for the purification of myosin-V from SF9 cells. We thank O. Hofnagel and D. Prumbaum for assistance with data collection. We thank W. Linke and A. Unger for providing us with muscle acetone powder. We thank R.S. Goody for his friendly advice. This work was supported by the Max Planck Society (to S.R.), the European Research Council under the European Union’s Horizon 2020 Programme (ERC-2019-SyG, grant 856118) (to S.R.), the Centre national de la recherche scientifique (to A.H.), the Agence nationale de la recherche (grant ANR-17-CE11-0029-01) (to A.H.) and the National Institutes of Health (grant R01-DC009100) (to H.L.S.). S.P. was supported as a fellow of Studienstiftung des deutschen Volkes.

---

**Table 1—Supplementary Figure 1.**
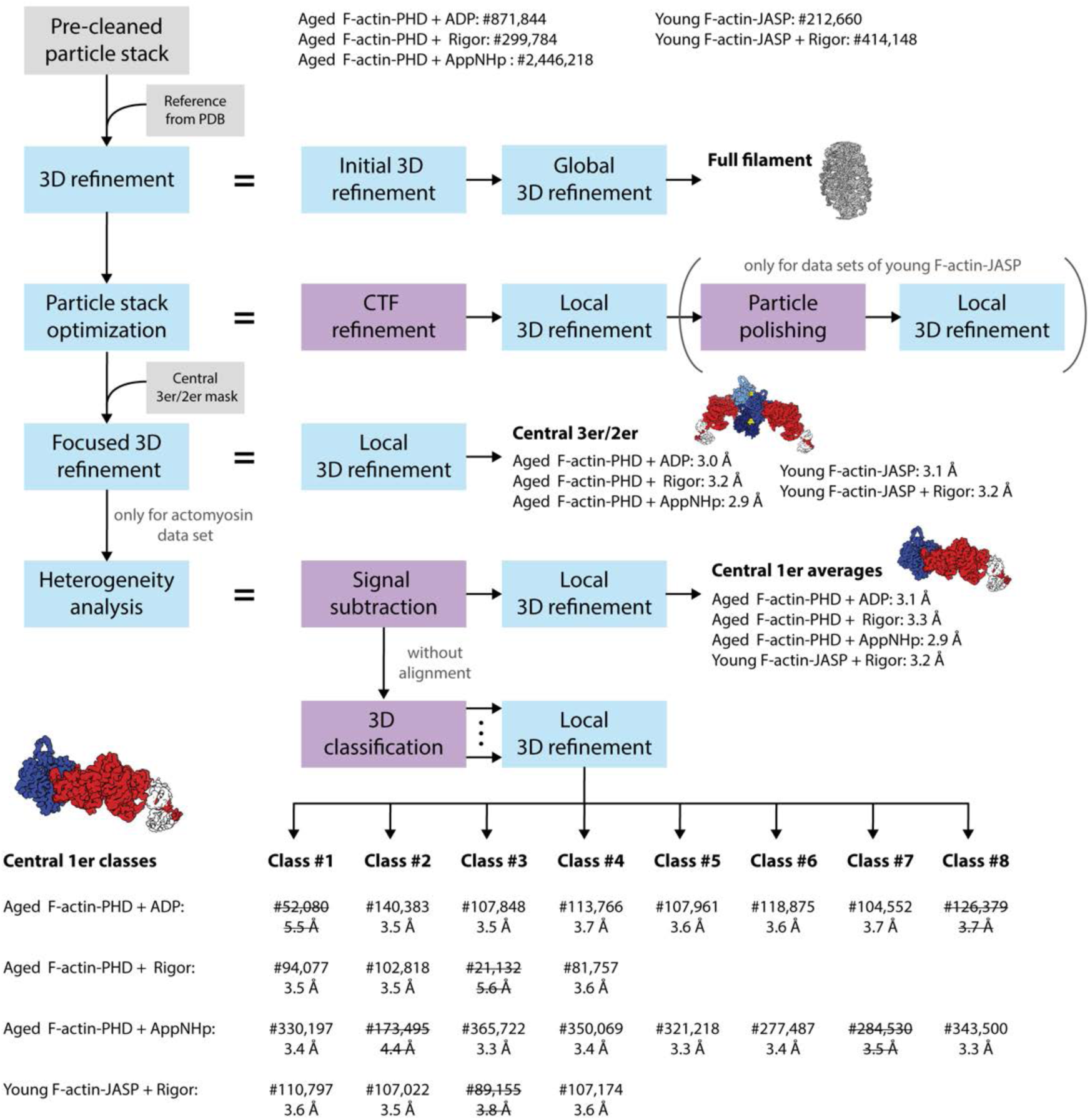
Schematic of the cryo-EM processing pipeline. Auto-picked particle stacks were initially pre-cleaned by 2D classification (final number of particles stated). Cleaned stacks were 3D refined against an initial reference generated from an atomic model of actomyosin (PDB: 5JLH, (von der Ecken et al., 2016)) without applying a 3D mask. The resulting 3D density map was used as reference volume in a subsequent masked 3D refinement yielding a first high-resolution structure of the full actomyosin filament. Based on this, particle stacks were optimized by CTF refinement and in case of the young F-actin-JASP data sets by additional particle polishing, followed by a local 3D refinement. By applying a mask including only the central three actin and two myosin molecules (central 3er/2er) the refinement was subsequently focused on the central section of the filament (central 3er/2er maps, resolutions stated). To account for the structural heterogeneity observed in the actomyosin data sets, a heterogeneity analysis was performed. Here, particles were initially signal subtracted to remove everything but the central actomyosin subunit (central 1er). These particles were then locally 3D refined to produce average structures (central 1er maps, resolutions stated). In addition, signal subtracted particles were 3D classified without alignment to separate distinct conformations. The number of classes was optimized experimentally to yield a maximum number of high-resolution 3D classes. Finally, subsets were locally 3D refined resulting in 18 high-resolution structures (central 1er classes, final number of particles and resolutions stated). Classes of insufficient quality (struck through) were not modelled and omitted in all subsequent analysis steps.

**Table 1—Supplementary Figure 2.**
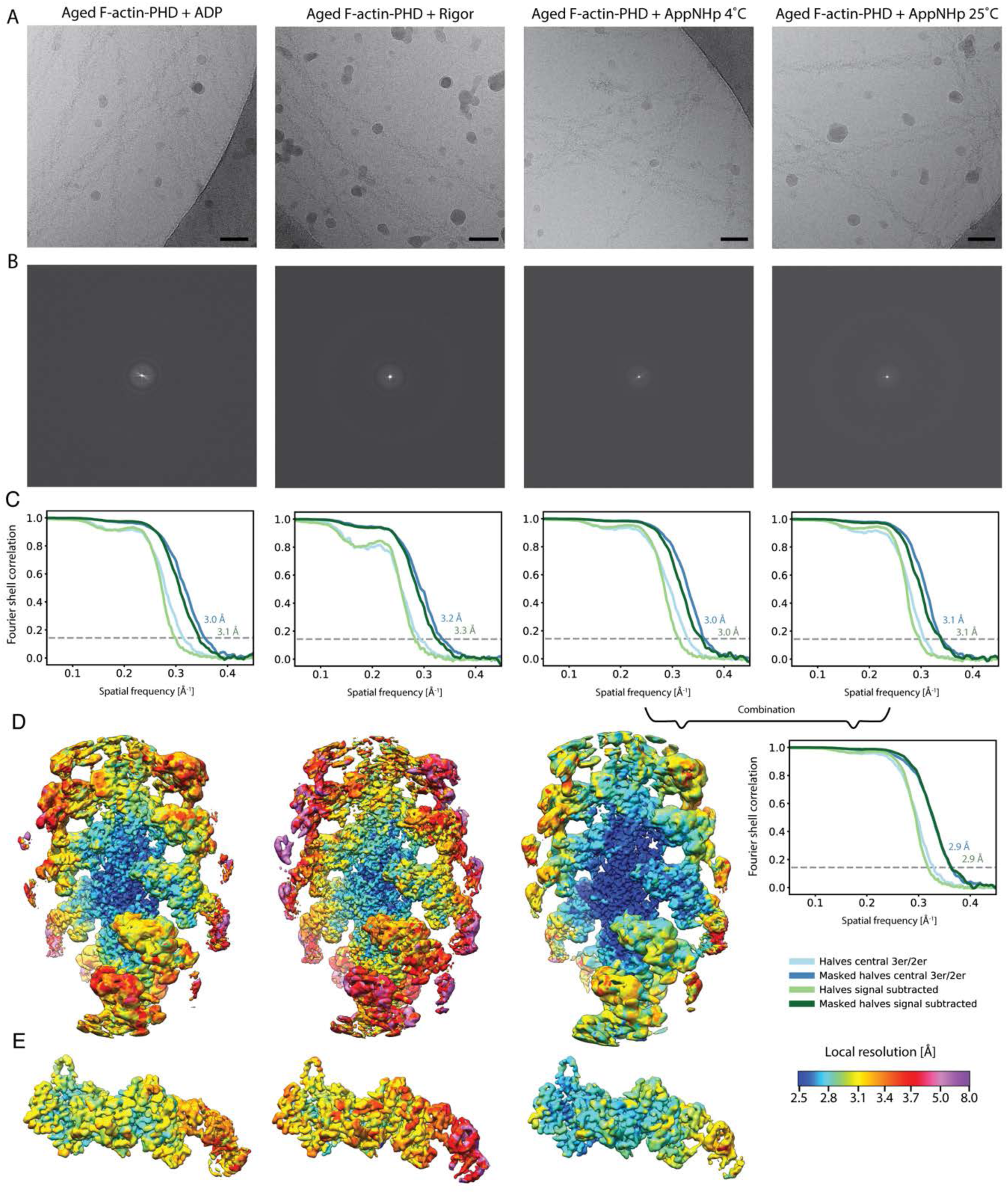
Overview of the cryo-EM data and resolution of aged F-actin-PHD in complex with myosin-V in the rigor, ADP and AppNHp state. **(A)** Representative micrographs at -1.3 μm defocus and **(B)** their power spectra. **(C)** Fourier shell correlation (FSC) curves for masked (darker shade, with resolution values) and unmasked half maps (lighter shade) including either three actin subunits and two myosin molecules (central 3er/2er, shades of blue, also see Table 1—Supplementary Figure 1) or one actomyosin subunit (signal subtracted, central 1er, shades of green). **(D)** Color-coded local resolution of full filaments and **(E)** signal subtracted actomyosin subunits for all three states. Note that the two AppNHp data sets (4 °C and 25 °C) were combined to increase the overall resolution. Scale bar 500 Å.

**Table 1 —Supplementary Table 1.**
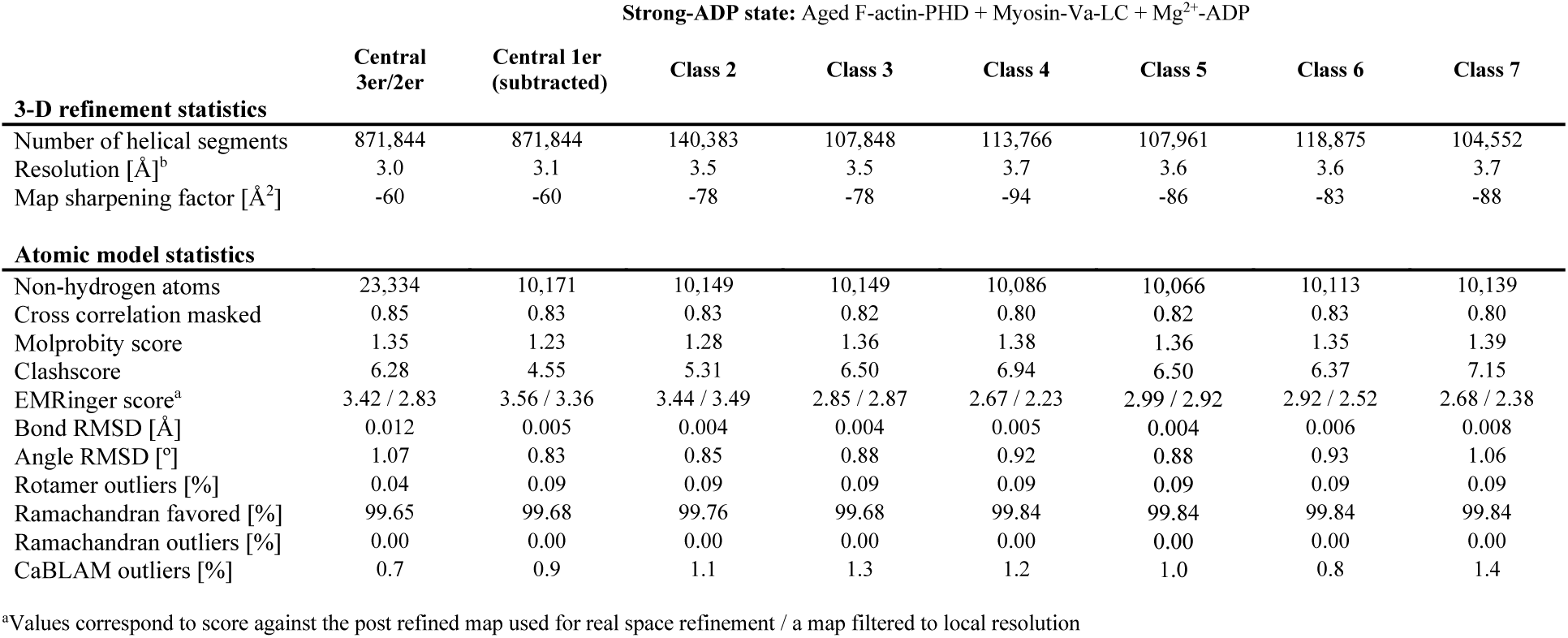
Refinement and model building statistics of aged F-actin-PHD in complex with myosin-V in the strong-ADP state.

**Table 1 —Supplementary Table 2.**
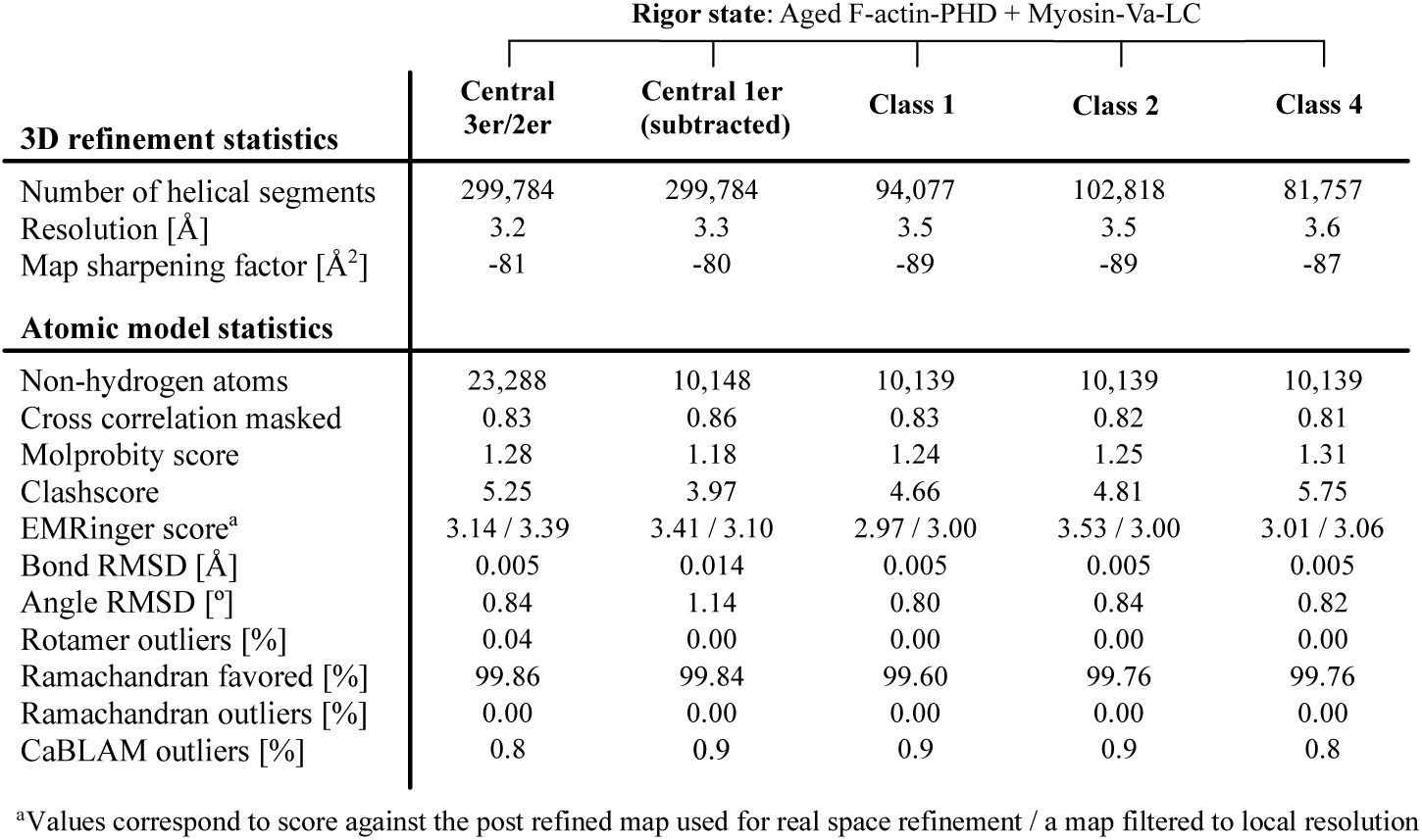
Refinement and model building statistics of aged F-actin-PHD in complex with myosin-V in the rigor state.

**Table 1 —Supplementary Table 3.** Refinement and model building statistics of aged F-actin-PHD in complex with myosin-V in the PRT state (bound to AppNHp).

| PRT state: Aged F-actin-PHD + Myosin-Va-LC + Mg<sup>2+</sup>-AppNHP |  |  |  |  |  |  |  |  |
| --- | --- | --- | --- | --- | --- | --- | --- | --- |
|  | Central<br>3er/2er | Central 1er<br>(subtracted) | Class 1 | Class 3 | Class 4 | Class 5 | Class 6 | Class 8 |
| **3D refinement statistics** |  |  |  |  |  |  |  |  |
| Number of helical segments | 2,446,218 | 2,446,218 | 330,197 | 365,722 | 350,069 | 321,218 | 277,487 | 343,500 |
| Resolution [Å] | 2.9 | 2.9 | 3.4 | 3.3 | 3.4 | 3.3 | 3.4 | 3.3 |
| Map sharpening factor [Å<sup>2</sup>] | -80 | -100 | -113 | -106 | -114 | -106 | -111 | -104 |
| **Atomic model statistics** |  |  |  |  |  |  |  |  |
| Non-hydrogen atoms | 23,370 | 10,189 | 10,125 | 10,189 | 10,154 | 10,189 | 10,085 | 10,189 |
| Cross correlation masked | 0.85 | 0.84 | 0.84 | 0.86 | 0.85 | 0.85 | 0.83 | 0.84 |
| Molprobity score | 1.25 | 1.15 | 1.17 | 1.24 | 1.20 | 1.26 | 1.37 | 1.18 |
| Clashscore | 4.76 | 3.56 | 3.78 | 4.64 | 4.12 | 4.99 | 6.74 | 3.95 |
| EMRinger score<sup>a</sup> | 3.29 / 3.45 | 3.82 / 3.40 | 3.35 / 3.07 | 3.58 / 3.45 | 3.18 / 3.35 | 2.94 / 2.97 | 3.01 / 3.01 | 3.09 / 2.88 |
| Bond RMSD [Å] | 0.004 | 0.012 | 0.009 | 0.014 | 0.009 | 0.014 | 0.009 | 0.005 |
| Angle RMSD [°] | 0.78 | 1.01 | 0.96 | 1.15 | 0.97 | 1.14 | 1.08 | 0.81 |
| Rotamer outliers [%] | 0.08 | 0.09 | 0.09 | 0.09 | 0.09 | 0.09 | 0.09 | 0.09 |
| Ramachandran favored [%] | 99.86 | 99.84 | 99.84 | 99.84 | 99.84 | 99.84 | 99.84 | 99.84 |
| Ramachandran outliers [%] | 0.00 | 0.00 | 0.00 | 0.00 | 0.00 | 0.00 | 0.00 | 0.00 |
| CaBLAM outliers [%] | 1.1 | 1.2 | 1.1 | 1.1 | 1.2 | 1.2 | 1.4 | 0.7 |

<sup>a</sup>Values correspond to score against the post refined map used for real space refinement / a map filtered to local resolution

**Figure 1—figure supplement 1.**
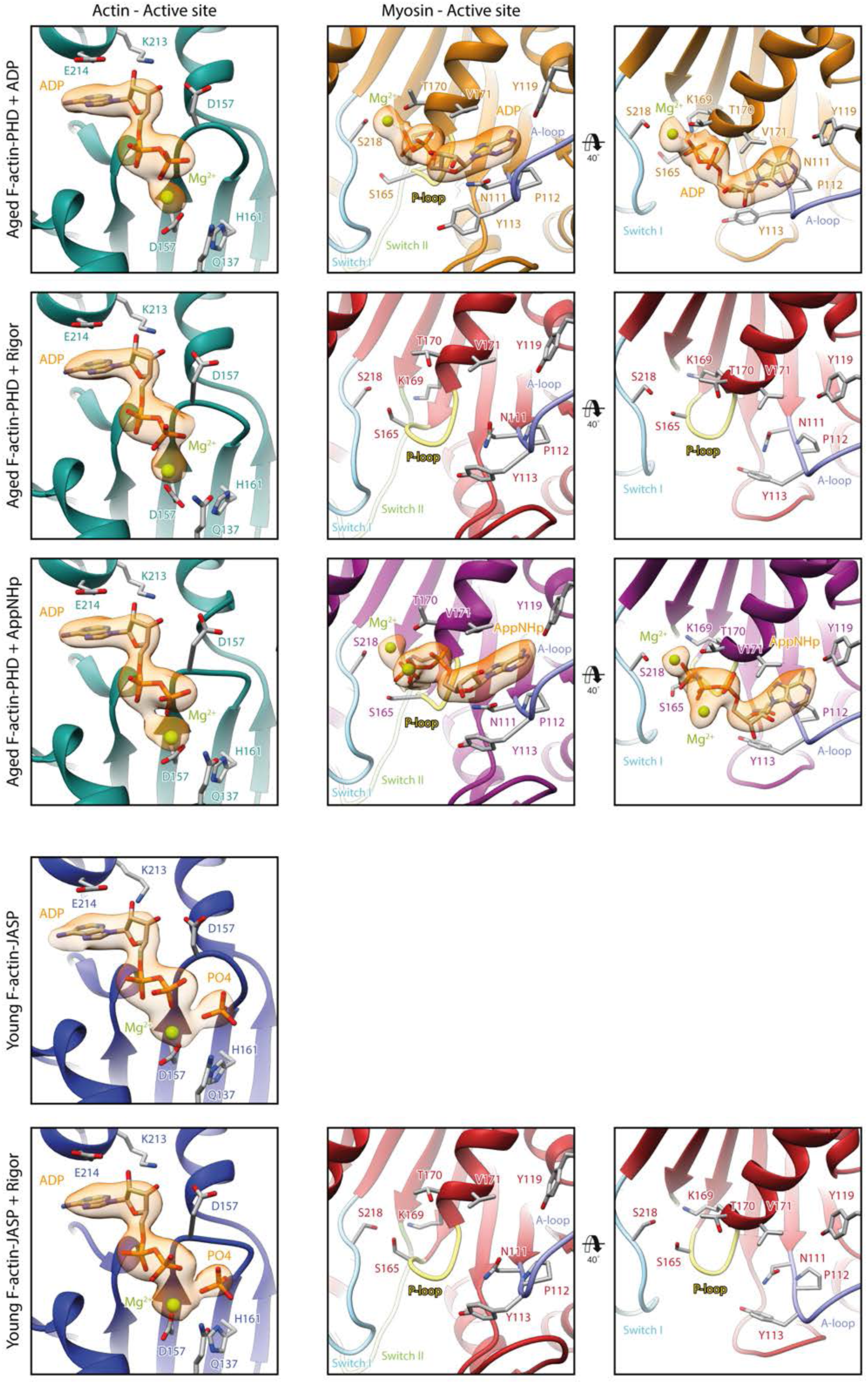
Nucleotide densities at and organization of the active sites of actin and myosin. Close-up views of the active site of F-actin (left column) and myosin-V (middle and right column) of all five structures. Ribbons are color-coded according to the respective structural state; aged F-actin-PHD: sea green, young F-actin-JASP: blue, and myosin-V in the rigor: red, ADP: orange and AppNHp state: purple. Key loops of the myosin active site are highlighted by pastel colors; P-loop: yellow, switch I: blue, switch II: green and A-loop: purple. Nucleotide densities are shown in orange and clearly support the presence of P_i_ in young JASP-stabilized F-actin. While there is density for a Mg^2+^ ion in all occupied active sites, there is an additional density, likely corresponding to a second Mg^2+^ ion, in AppNHp-bound myosin.

**Figure 1—figure supplement 2.**
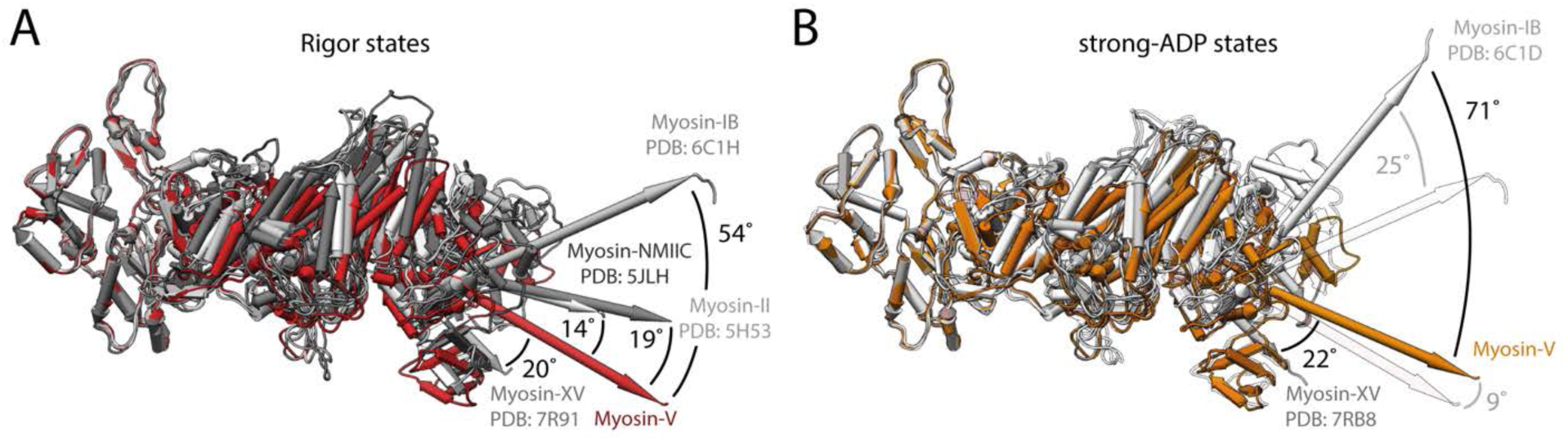
Structural variations of the rigor and strong-ADP states of different actomyosin complexes. Comparison of atomic models of the rigor and strong-ADP states of different actomyosin complexes solved by cryo-EM. **(A)** Superposition of the rigor states of myosin-V (red), myosin-II (PDB: 5H53, (Fujii & Namba, 2017)), myosin-NMIIC (PDB: 5JLH, (von der Ecken et al., 2016)), myosin-IB (PDB: 6C1H, (Mentes et al., 2018)) and myosin-XV (PDB: 7R91, (Gong et al., 2021)) (shades of gray), illustrating strongly varying conformations and lever arm orientations. **(B)** Superposition of the strong-ADP states of myosin-V (orange), myosin-IB (PDB: 6C1D, (Mentes et al., 2018)) and myosin-XV (PDB: 7RB8, (Gong et al., 2021)) (shades of gray). The corresponding rigor states are shown as transparent. The difference in the orientation of the lever arm is even more pronounced in the strong-ADP state, increasing from a relative rotation of 54° to 71° for myosin V and myosin-IB. Structures are shown without the light chain after alignment on the central actin subunit.

**Figure 2—figure supplement 1.**
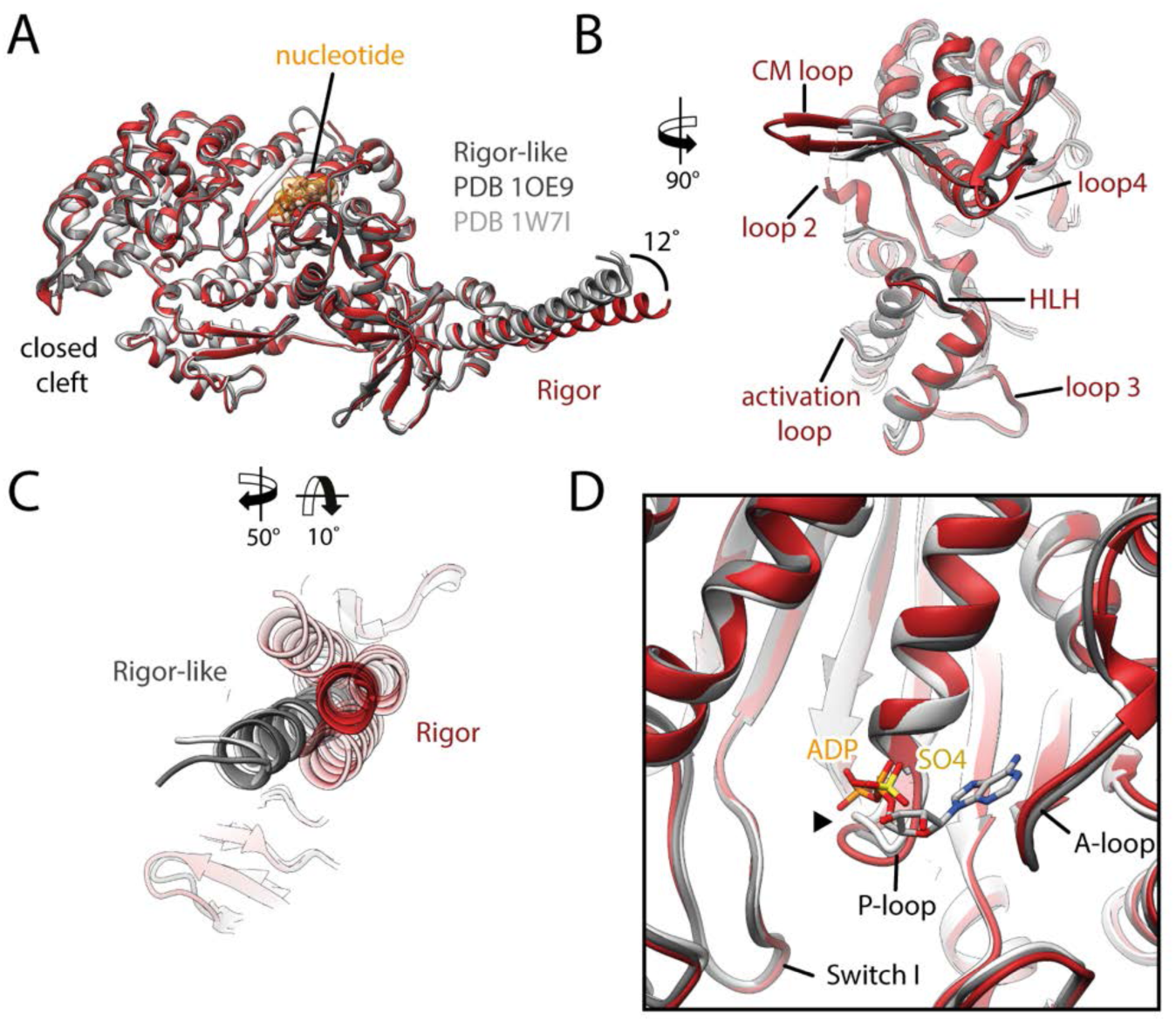
Comparison of the rigor state of myosin-V with crystal structures of the rigor-like state. Comparison of the rigor state actomyosin-V cryo-EM structure (red) with crystal structures of the same myosin in the rigor-like state (PDB: 1OE9, (Coureux et al., 2003) and PDB: 1W7I, also called weak-ADP state (Coureux et al., 2004), shades of gray). **(A)** Superposition of the rigor state with two rigor-like crystal structures. Deviations are limited to the actin interface, in particular the CM loop, loop 4 and loop 2 **(B**) and the lever arm **(C)**. Interestingly, the lever arm orientation seen in the rigor-like states does not superimpose with any conformation seen for the rigor complex (average: red and 3D classes: transparent red), but localizes outside of its conformational space. **(D)** The active site is open in both the rigor and rigor-like states and the SO_4_ and ADP bound to the rigor-like crystal structures only give rise to small, isolated changes of the P-loop (highlighted by a black arrowhead).

**Figure 4—figure supplement 1.**
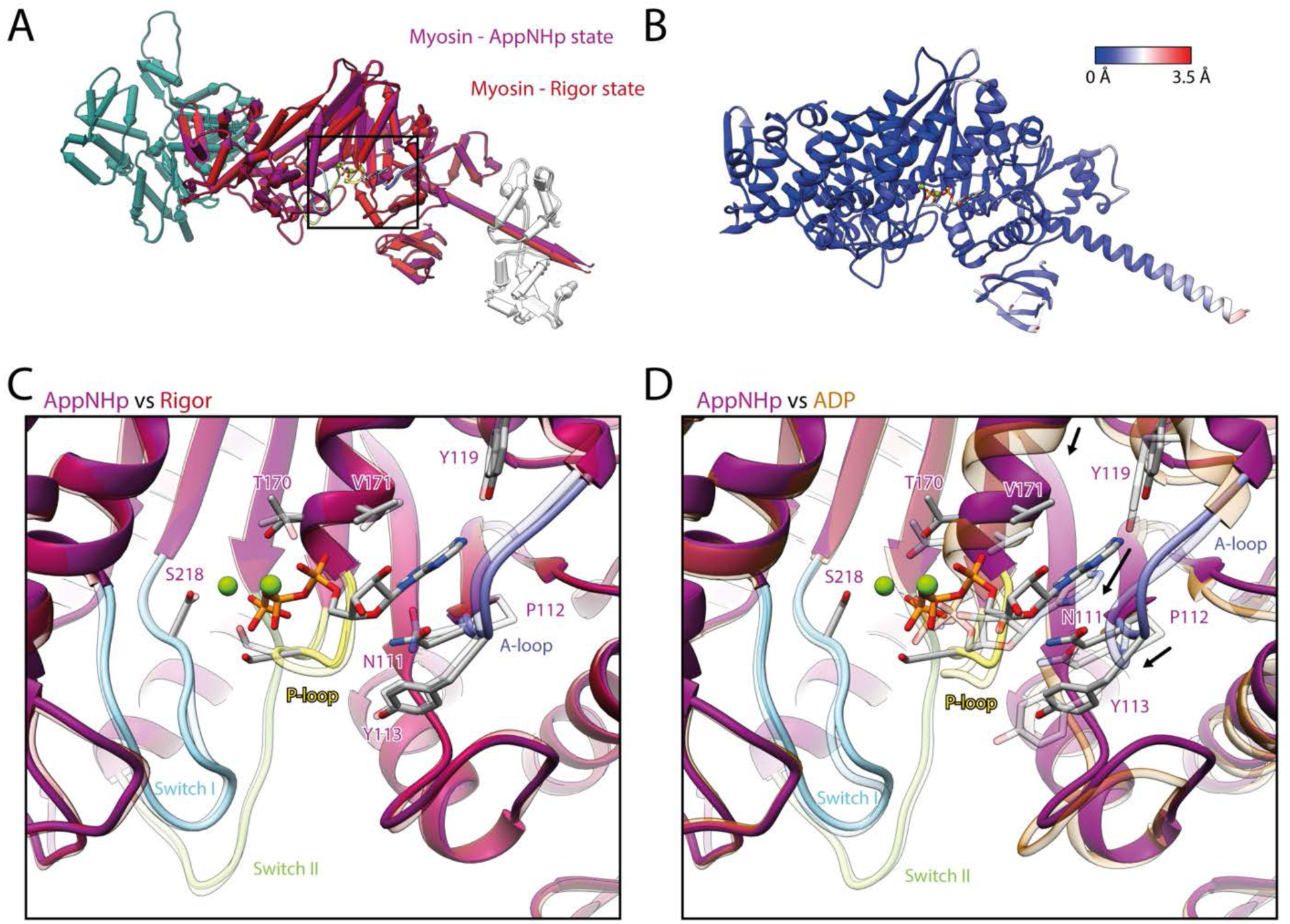
Comparison of AppNHp-bound myosin-V with myosin-V in the rigor and strong-ADP state. **(A)** Superposition and **(B)** color-coded backbone root mean square deviation (RMSD) of the rigor (red) and the AppNHp (purple) atomic models, illustrating their close resemblance. Higher RMSD values localize exclusively to regions of lower local resolution and thus are likely due to modeling inaccuracies. A black box highlights the position of the active site. **(C-D)** Comparison of the active site of myosin bound to AppNHp (purple) with the one of myosin in the rigor (transparent red) **(C)** and strong-ADP state (transparent orange) **(D)**. While only small, local changes are associated with the binding of AppNHp **(C)**, the active sites of AppNHp- and ADP-bound myosin-V differ markedly (rearrangements are indicated by black arrows). Models were aligned on F-actin, for color code see Figure 1.

**Figure 4—figure supplement 2.**
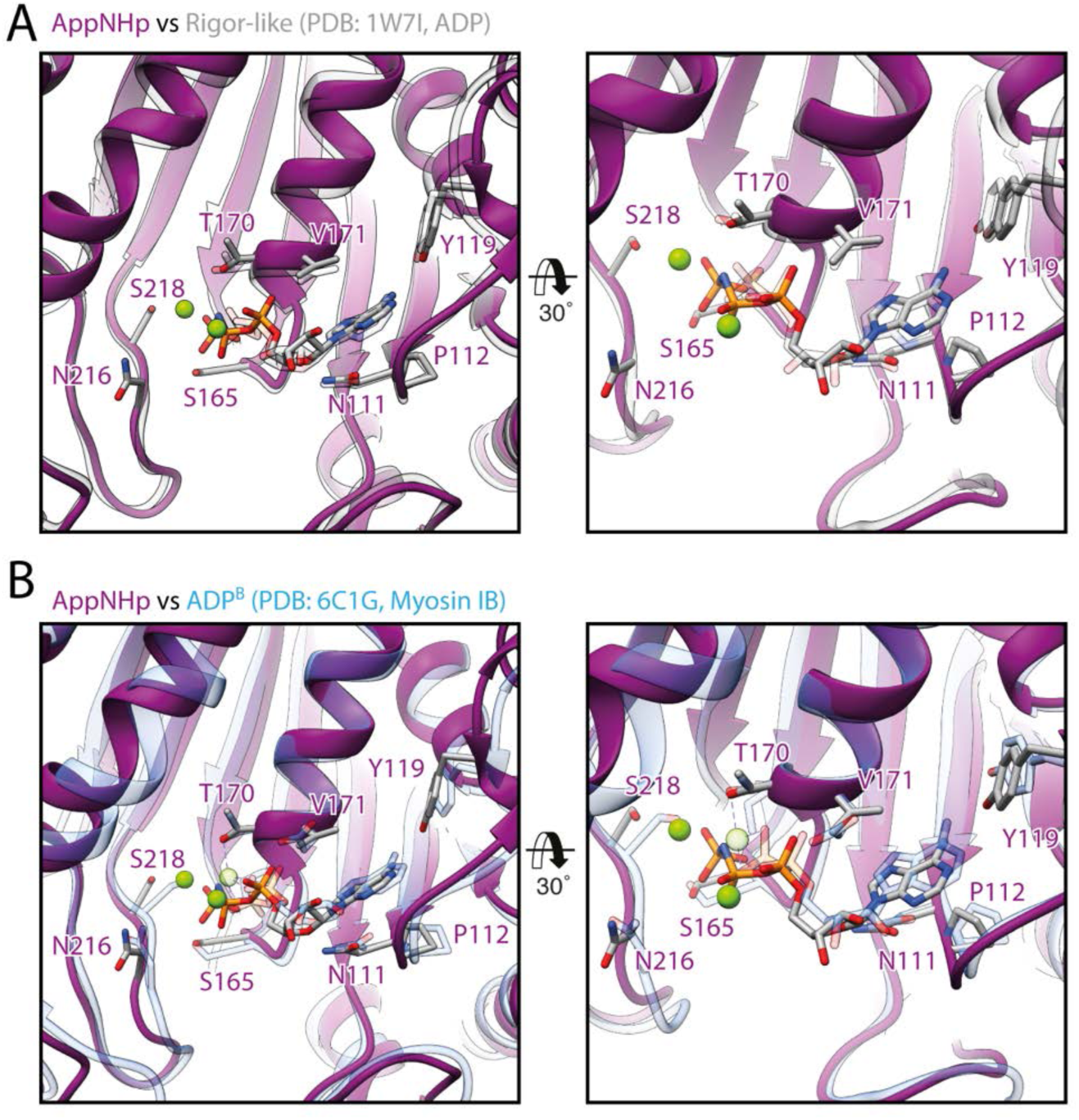
Common active site conformation in transition states with weakly-bound nucleotide. **(A-B)** Comparison of the active site of myosin-V bound to AppNHp with the one of **(A)** rigor-like myosin-V with ADP weakly-bound (PDB: 17WI, also known as weak-ADP state, transparent gray, (Coureux et al., 2004)) and **(B)** ADP-bound myosin-IB in a strong-ADP to rigor transition state (PDB: 6C1G, transparent blue, (Mentes et al., 2018)). Models were aligned on the HF helix (aa 169-190, trailing the P-loop). The overall conformation of the active site as well as the coordination of the nucleotide is remarkably similar in all three structures suggesting it to be characteristic for actin-bound transition states with weakly-bound nucleotide.

**Figure 4—figure supplement 3.**
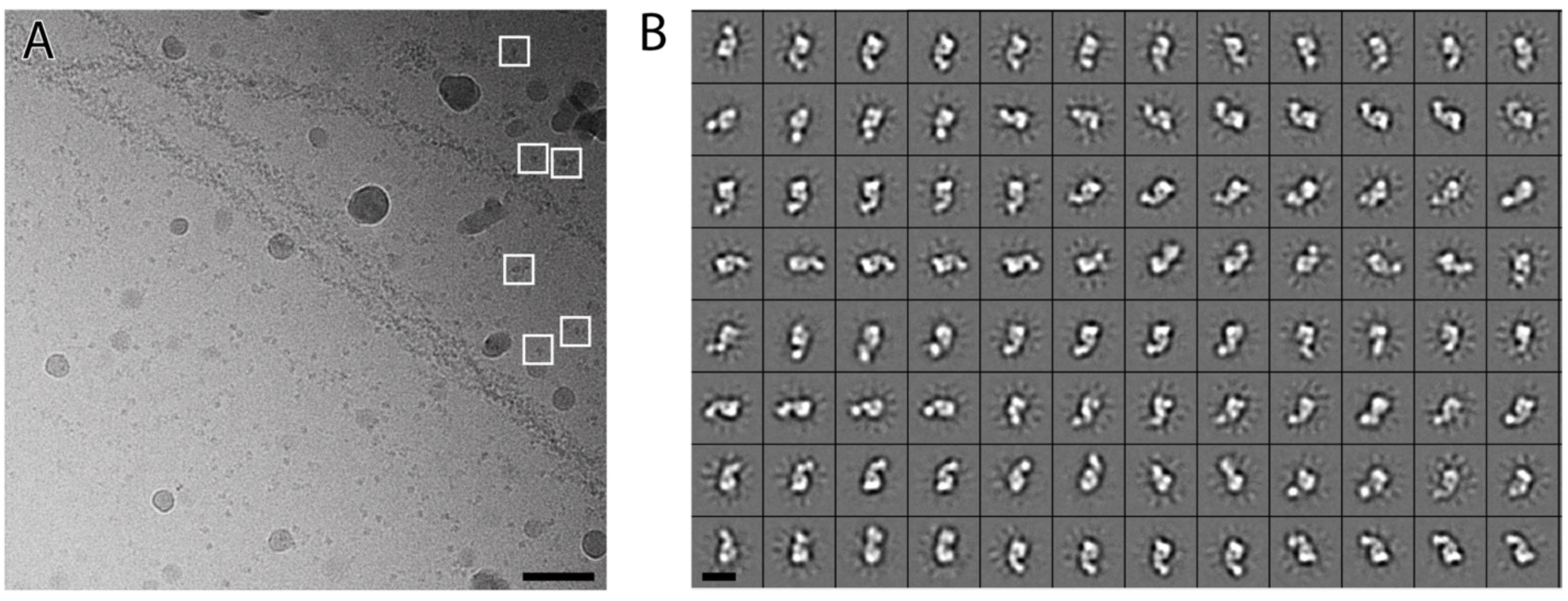
2D cryo-EM data of unbound myosin-V in complex with AppNHp. **(A**) Representative contrast-enhanced micrograph of the aged actomyosin-V complex bound to AppNHp (sample plunged at 4 °C) illustrating many unbound particles in the background, which likely correspond to myosin-V molecules. Particle boxes (white) are shown for a few particles to highlight their size. Scale bar 500 Å. **(B)** Representative 2D class averages confirming that the particles in the background are indeed unbound myosin-V molecules. Scale bar 100 Å.

**Figure 6—Table 1.** Overview of helical symmetry parameters of aged PHD-stabilized and young JASP-stabilized actomyosin-V complexes. For a direct comparison, the parameters of aged F-actin-PHD (PDB: 6T20, (Pospich et al., 2020)) and young F-actin-JASP (PDB: 5OOD, (Merino et al., 2018)) are shown alongside. Differences in both the helical rise and twist can be readily explained by errors of the pixel size, which is not identical for all data sets. Helical parameters were estimated from the atomic model of five consecutive subunits independently fitted into the map, see (Pospich et al., 2017) for details. To make results more comparable, only actin subunits were considered during fitting. Note that fitting inaccuracies can also give rise to small deviations.

| Helical symmetry | Rise [Å] | Twist [°] | Pixel size [Å] |
| --- | --- | --- | --- |
| Aged F-actin-PHD + Rigor | 27.82 +/- 0.02 | -167.27 +/- 0.02 | 1.06 |
| Aged F-actin-PHD + ADP | 27.81 +/- 0.02 | -167.32 +/- 0.02 | 1.06 |
| Aged F-actin-PHD + AppNHP | 27.77 +/- 0.02 | -167.32 +/- 0.02 | 1.10 |
| Aged F-actin-PHD (PDB: 6T20) | 27.59 +/- 0.02 | -166.9 +/- 0.1 | 1.14 |
| Young F-actin-JASP | 27.85 +/- 0.08 | -166.87 +/- 0.02 | 1.10 |
| Young F-actin-JASP + Rigor | 27.72 +/- 0.01 | -167.06 +/- 0.02 | 1.10 |
| Young F-actin-JASP (PDB: 5OOD) | 27.39 | -166.41 | 1.09 |

**Figure 7—Table 1.**
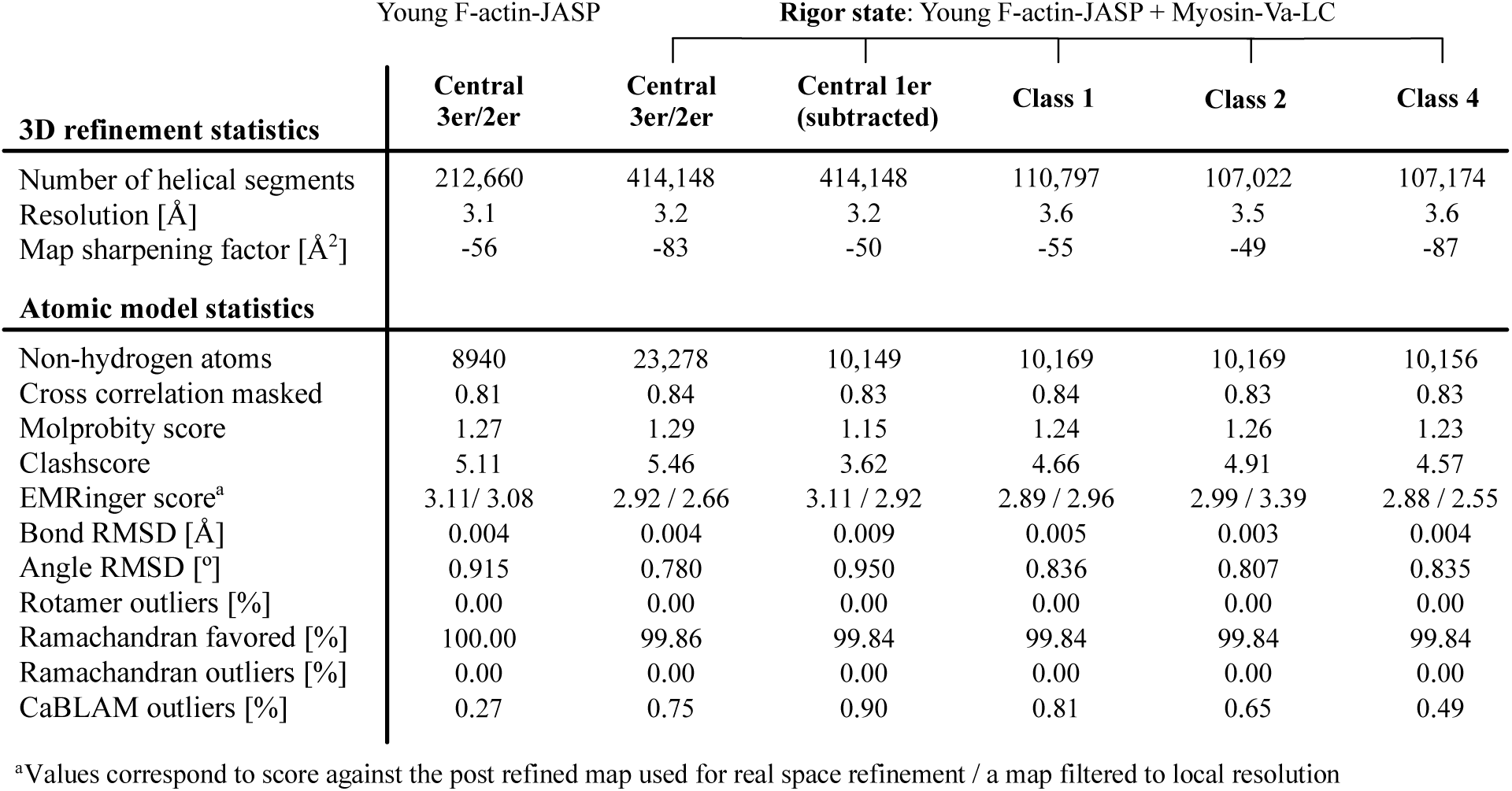
Refinement and model building statistics of young F-actin-JASP alone and in complex with myosin-V in the rigor state.

**Figure 7—figure supplement 1.**
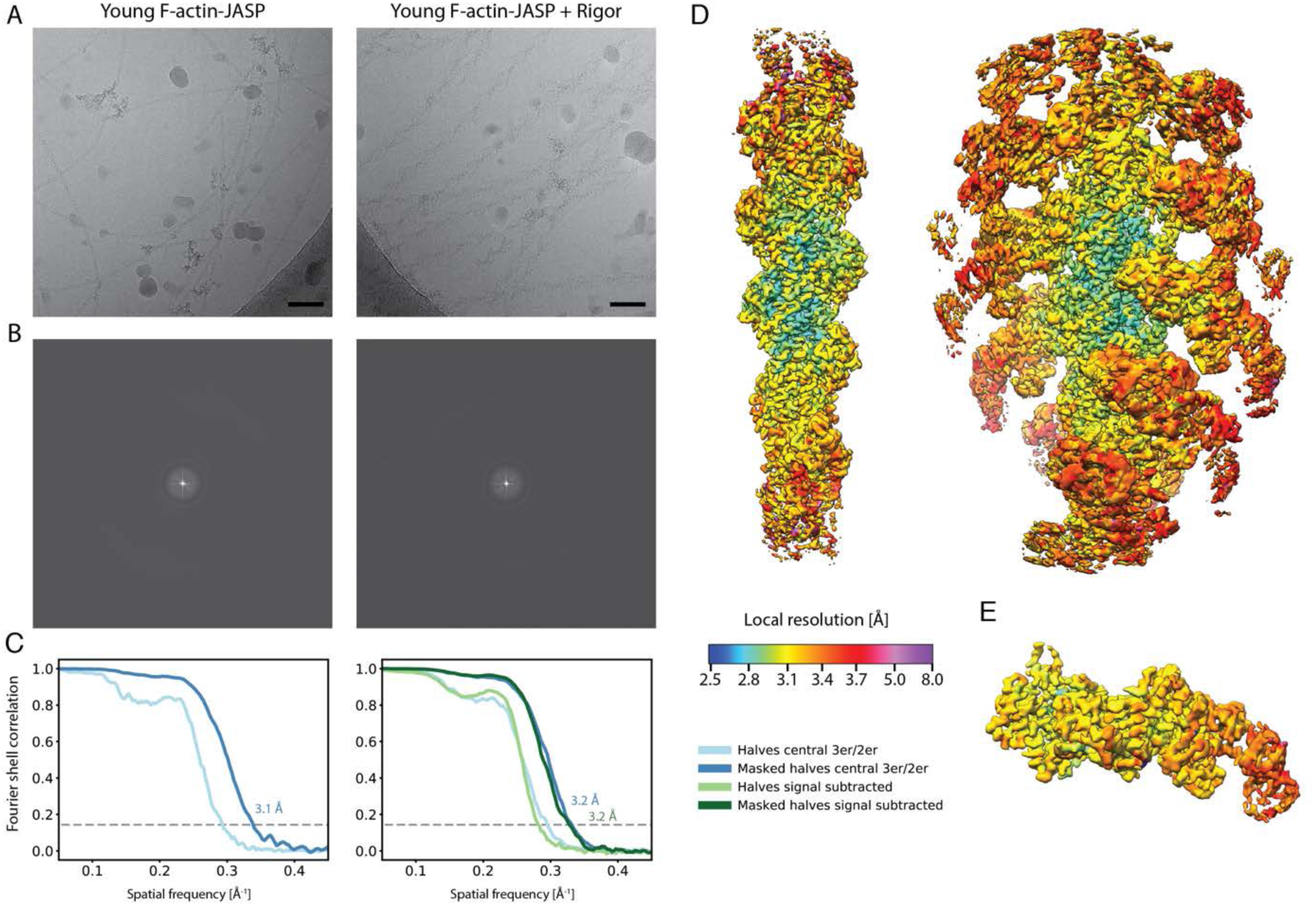
Overview of the cryo-EM data and resolution of young F-actin-JASP alone and in complex with myosin-V in the rigor state. **(A)** Representative micrographs at -1.3 μm defocus and **(B)** their power spectra. **(C)** Fourier shell correlation (FSC) curves for masked (darker shade, with resolution values) and unmasked (lighter shade) half maps. For bare F-actin only the FSC of a map covering the central three subunits is shown (shades of blue), while for actomyosin either the FSC for three actin subunits and two myosin molecules (central 3er/2er, shades of blue) or for one actomyosin subunit (signal subtracted, central 1er, shades of green) is shown. **(D)** Color-coded local resolution of full filaments for both data sets and of the **(E)** signal subtracted central subunit of the young actomyosin complex. Note that signal subtraction was only performed for actomyosin complexes, also see Table 1—Supplementary Figure 1. Scale bar 500 Å.

**Figure 7— figure supplement 2.**
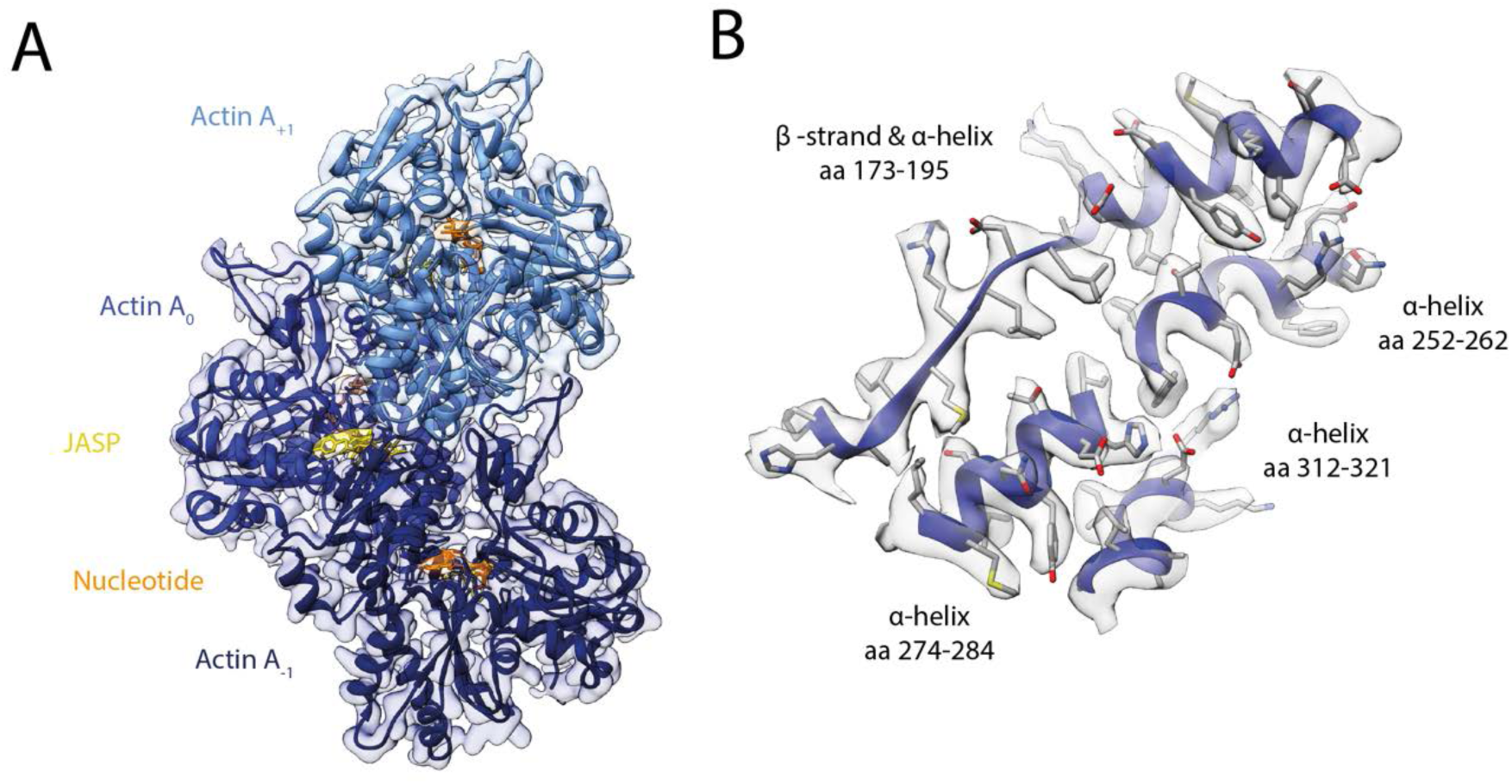
Structure of young JASP-stabilized F-actin. **(A**) Atomic model and LAFTER density map of young F-actin-JASP (shades of blue, three subunits shown, A_-1_ to A_+1_). Nucleotides and JASP are highlighted in orange and yellow, respectively; also see Figure 8—Video 1A-C. **(B)** Illustration of the model-map agreement within a central section of myosin. Most side chains are resolved by the post-refined density map (transparent gray).

**Figure 8—figure supplement 1.**
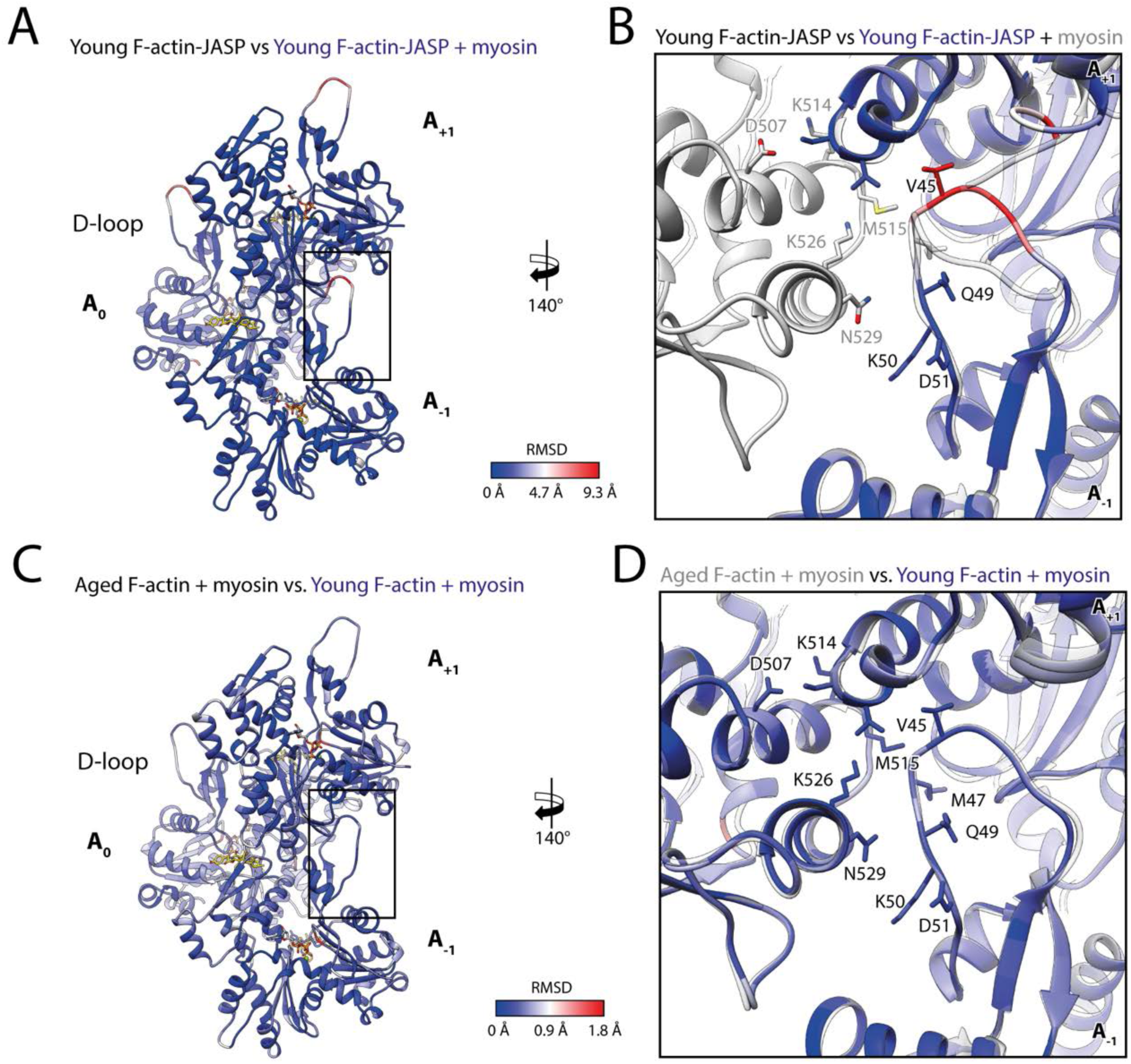
Myosin-V selects a specific conformation of F-actin. **(A-B)** Root mean square deviation (RMSD) of young F-actin-JASP before and after myosin binding illustrating a major, but spatially confined rearrangement of the D-loop and C-terminus interface. **(C-D)** RMSD highlighting the remarkable similarity of myosin-bound aged F-actin-PHD and myosin-bound young F-actin-JASP. Subunits were aligned individually to account for errors in the calibration of the pixel size.

**Figure 8—figure supplement 2.**
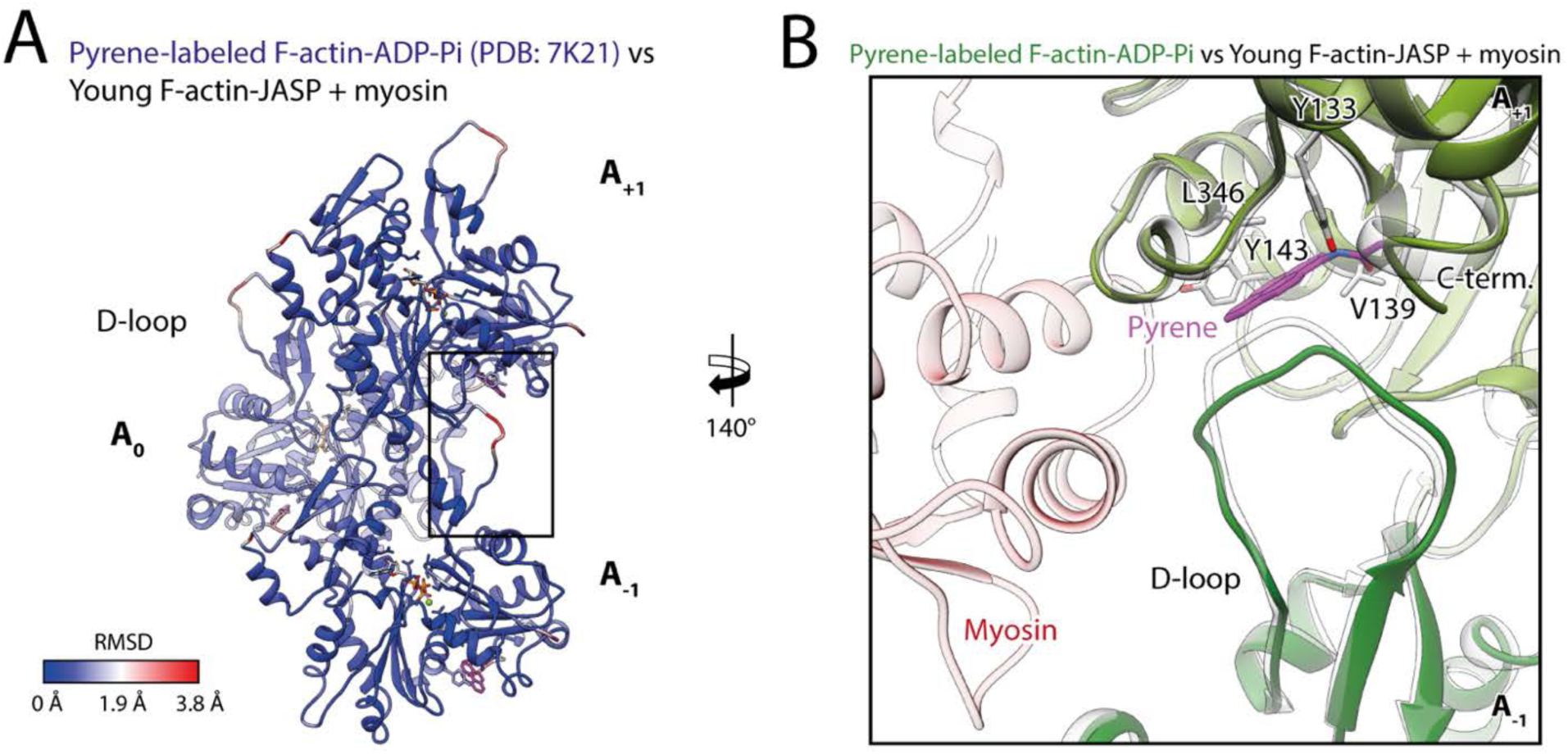
Pyrene-labeling potentially impedes selection of the closed D-loop. **(A)** Root mean square deviation (RMSD) of pyrene-labeled F-actin bound to ADP-P_i_ (PDB: 7K21, (Chou & Pollard, 2020)) and young F-actin-JASP in complex with myosin-V illustrating that differences primarily localize to the D-loop and C-terminus interface (black box). Subunits were aligned individually to account for errors in the calibration of the pixel size. **(B)** Close-up view of the D-loop C-terminus interface of pyrene-labeled F-actin bound to ADP-P_i_ (shades of green, PDB: 7K21, (Chou & Pollard, 2020)). Pyrene (magenta) wedges in-between the D-loop and C-terminus and thereby displaces the D-loop. In this way, pyrene likely interferes with myosin (transparent red) selecting the closed D-loop conformation (transparent gray).

**Figure 9—figure supplement 1.**
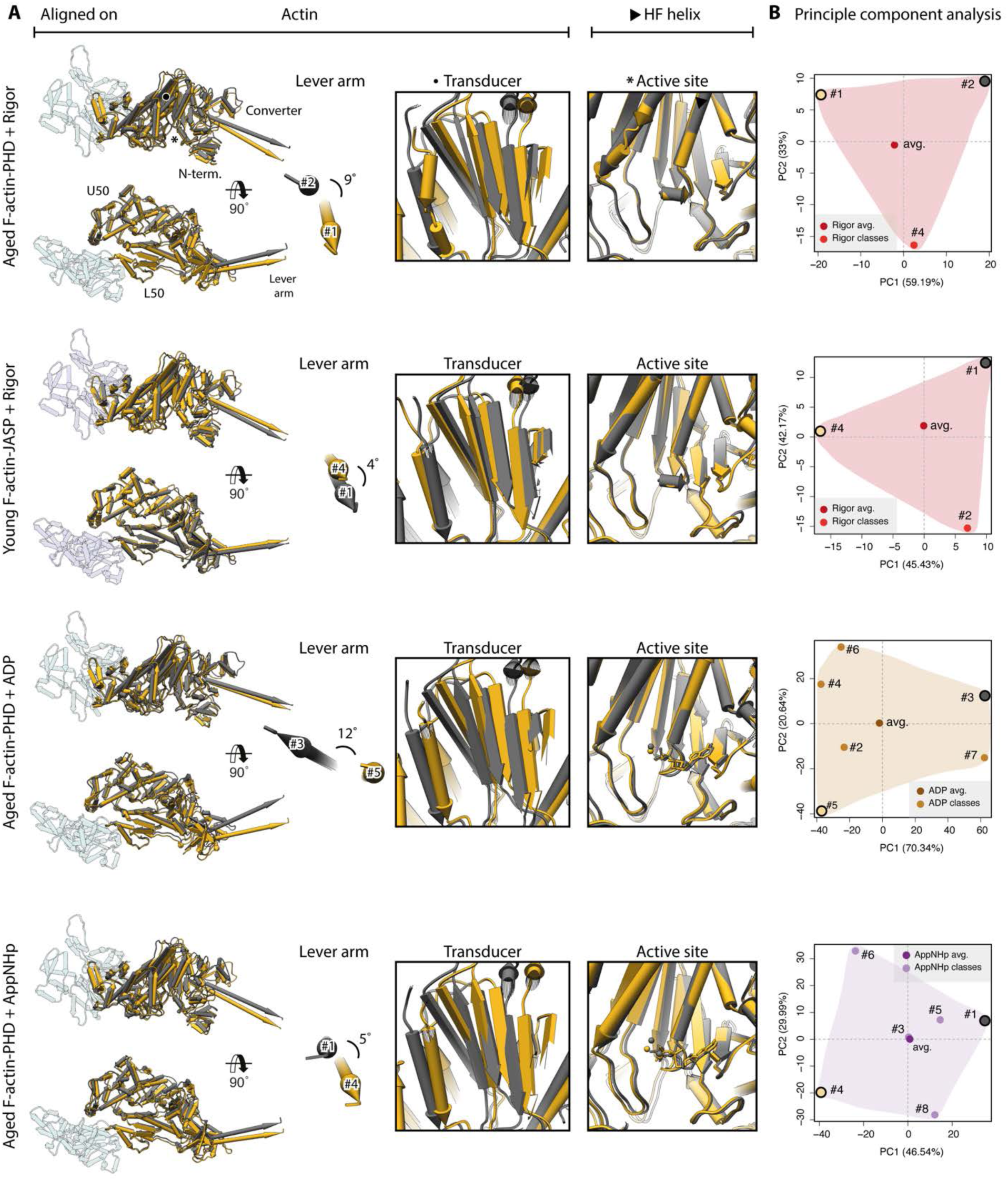
Extreme conformations of myosin-V. Extreme conformations of myosin-V in the rigor, strong-ADP and AppNHp-bound PRT state. **(A)** Superposition of atomic models as shown in Figure 9, but displaying only the extreme structures along the first principal component (yellow and gray). **(B)** Mapping of atomic models (average and 3D classes) into the first two principal components as shown in Figure 9. The localization of the extreme structures shown in A) is highlighted by a yellow and gray dot, respectively. **Figure 9—Video 1. Structural heterogeneity of myosin-V in the strong-ADP state**. **(A-B)** Three-dimensional visualization of the average structure of aged actomyosin-V in the strong-ADP state (myosin: orange, F-actin: sea green, PHD: yellow, LC: white). **(C)** Morph of 3D class average models illustrating the conformational heterogeneity of myosin (classes are ordered by their number, also see Table 1—Supplementary Figure 1). **(D-E)** Morph of extreme structures along the first **(D)** and second **(E)** principal component, also see Figure 9—figure supplement 1. **Figure 9—Video 2. Structural heterogeneity of myosin-V in the rigor state.** **(A-B)** Three-dimensional visualization of the average structure of aged actomyosin-V in the rigor state (myosin: red, F-actin: sea green, PHD: yellow, LC: white). **(C)** Morph of 3D class average models illustrating the conformational heterogeneity of myosin (classes are ordered by their number, also see Table 1—Supplementary Figure 1). **(D-E)** Morph of extreme structures along the first **(D)** and second **(E)** principal component, also see Figure 9—figure supplement 1. **(F-G)** Three-dimensional visualization of the average structure of young actomyosin-V in the rigor state (myosin: red, F-actin: blue, JASP: yellow, LC: white). Morph of 3D class average models **(H)** and morph of extreme structures along the first **(I)** and second **(J)** principal component. **Figure 9—Video 3. Structural heterogeneity of myosin-V in the PRT state (AppNHp).** **(A-B)** Three-dimensional visualization of the average structure of aged actomyosin-V in the AppNHp-bound PRT state (myosin: purple, F-actin: sea green, PHD: yellow, LC: white). **(C)** Morph of 3D class average models illustrating the conformational heterogeneity of myosin (classes are ordered by their number, also see Table 1—Supplementary Figure 1). **(D-E)** Morph of extreme structures along the first **(D)** and second **(E)** principal component, also see Figure 9—figure supplement 1.

**Figure 11—figure supplement 1.**
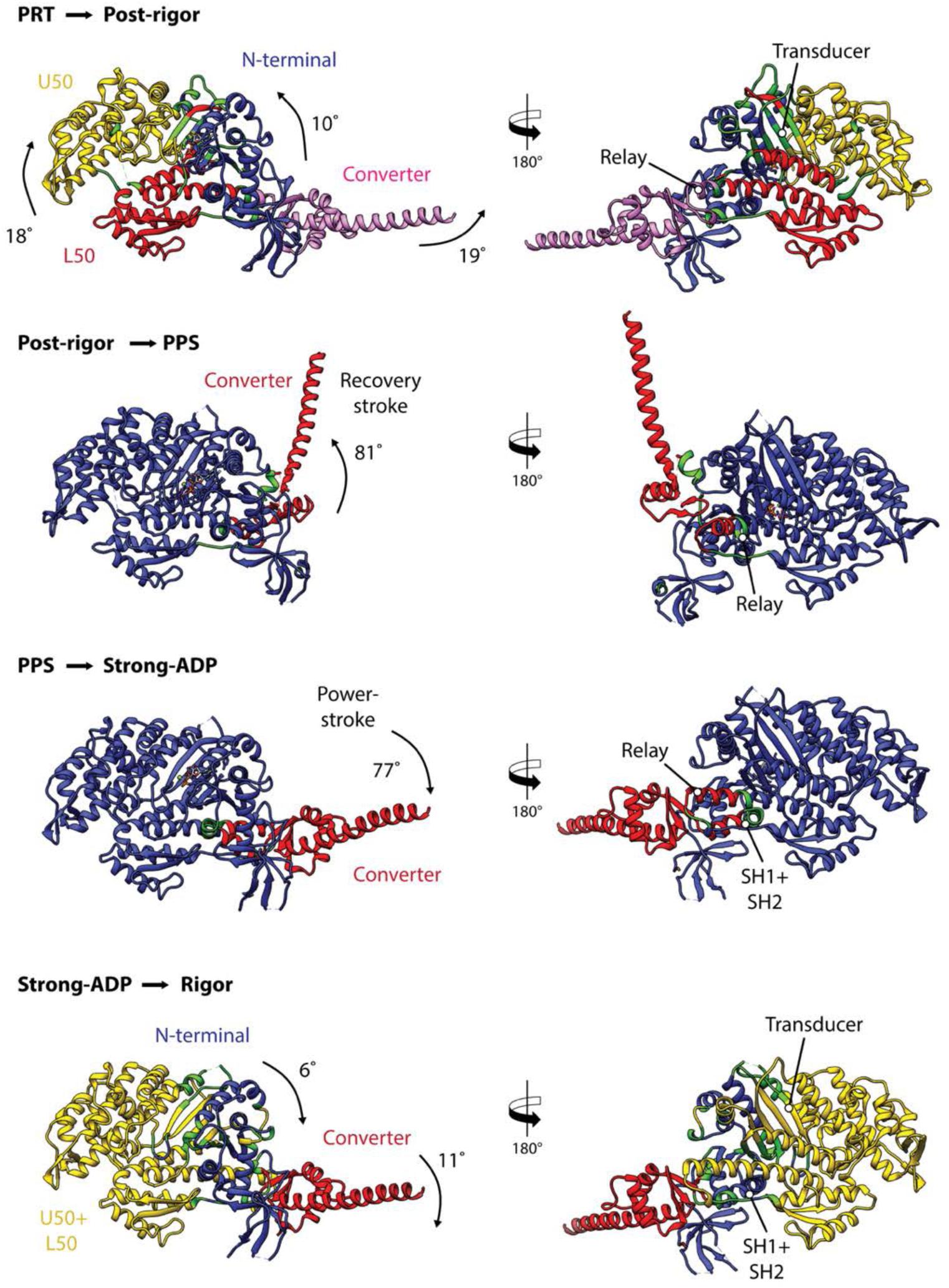
Domain motions during the motor cycle of myosin-V. Illustration of the domain motions during the motor cycle of myosin-V predicted by DynDom. Structures are shown according to their sequence in the motor cycle from top to bottom. Homology models were omitted. Hinge regions and bending residues are shown in green, while domains are shown in yellow, red, blue and pink, respectively. Only key structural elements are labeled. Note that some structural rearrangements, e.g., the closure of the actin binding cleft during the PPS to strong-ADP state transition, were not detected. This indicates that the underlying motions can possibly not be described by semi-independent domain movements, for example, due to local deformations of actin binding regions.

**Figure 11—figure supplement 2.**
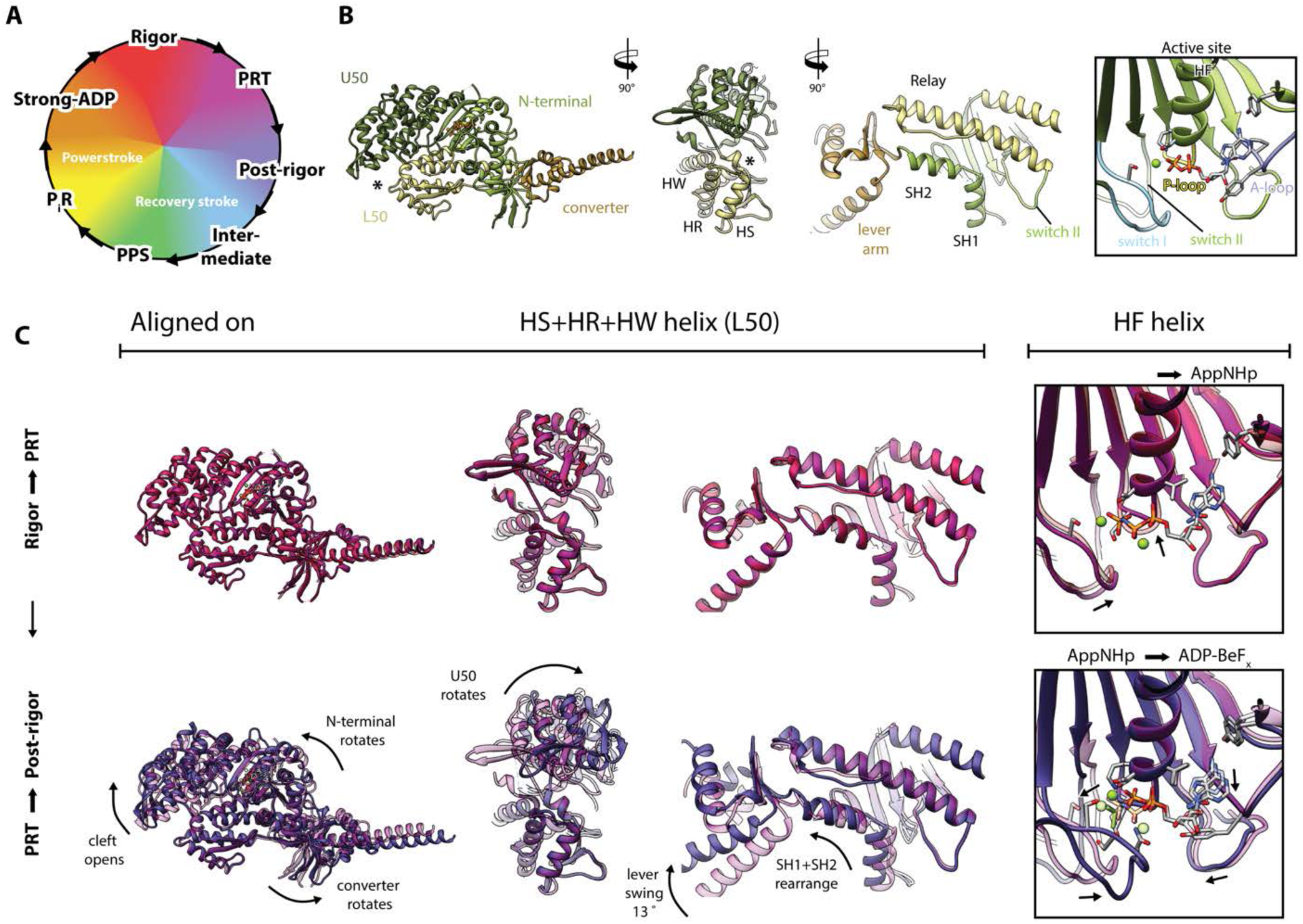
Structural transition of myosin-V along its motor cycle (Rigor to post-rigor state). Illustration of the conformational changes of myosin-V during its transition between key structural states. **(A)** Pictogram of the myosin motor cycle displaying the rainbow color-scheme used for the depiction of structural states (rigor: red, PRT: purple, post-rigor: slate blue, intermediate: light blue, PPS: green, P_i_R: yellow, strong-ADP: orange). **(B)** Color-coded legend of myosin highlighting domains and key structural elements. The actin binding cleft is labeled by an asterisk. **(C)** Superposition of atomic models of successive structural states along the myosin motor cycle. The precursor structure is shown as transparent. Major conformational changes are highlighted by black arrows. Structures are shown according to their sequence in the motor cycle from top to bottom and are continued in Figure 11—figure supplement 3 and Figure 11—figure supplement 4. Models are either aligned on the HR, HS and HW helices (overall conformation) or on the HF helix (active site).

**Figure 11—figure supplement 3.**
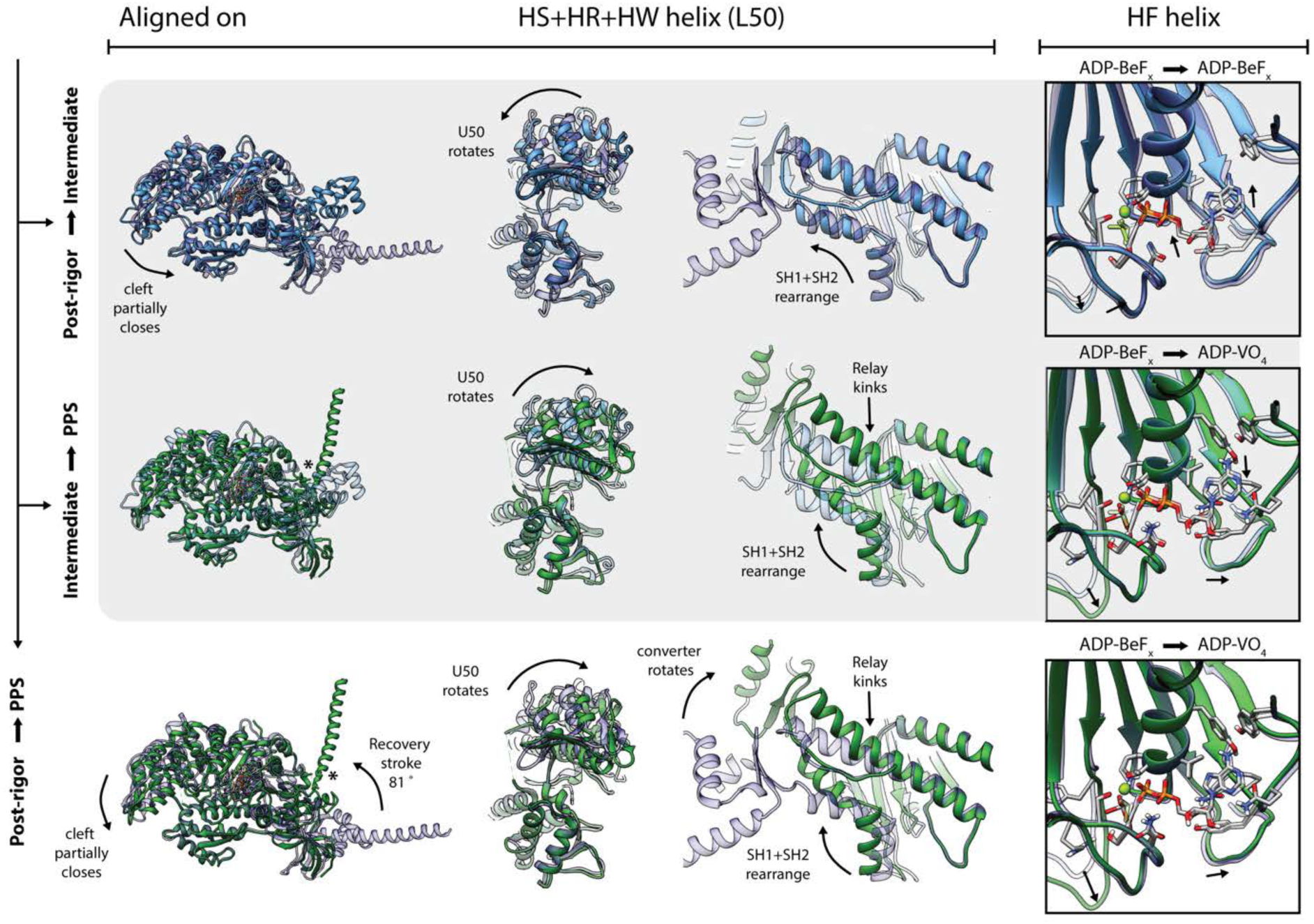
Structural transition of myosin-V along its motor cycle (Post-rigor to PPS state). Continuation of the superposition of atomic models of successive structural states along the myosin motor cycle shown in Figure 11—figure supplement 2, for continuation see Figure 11—figure supplement 4. The lever arm of the PPS state was computationally elongated (original length indicated by asterisk). Transitions involving the intermediate state homology model are grayed out to illustrate its limitations.

**Figure 11—figure supplement 4.**
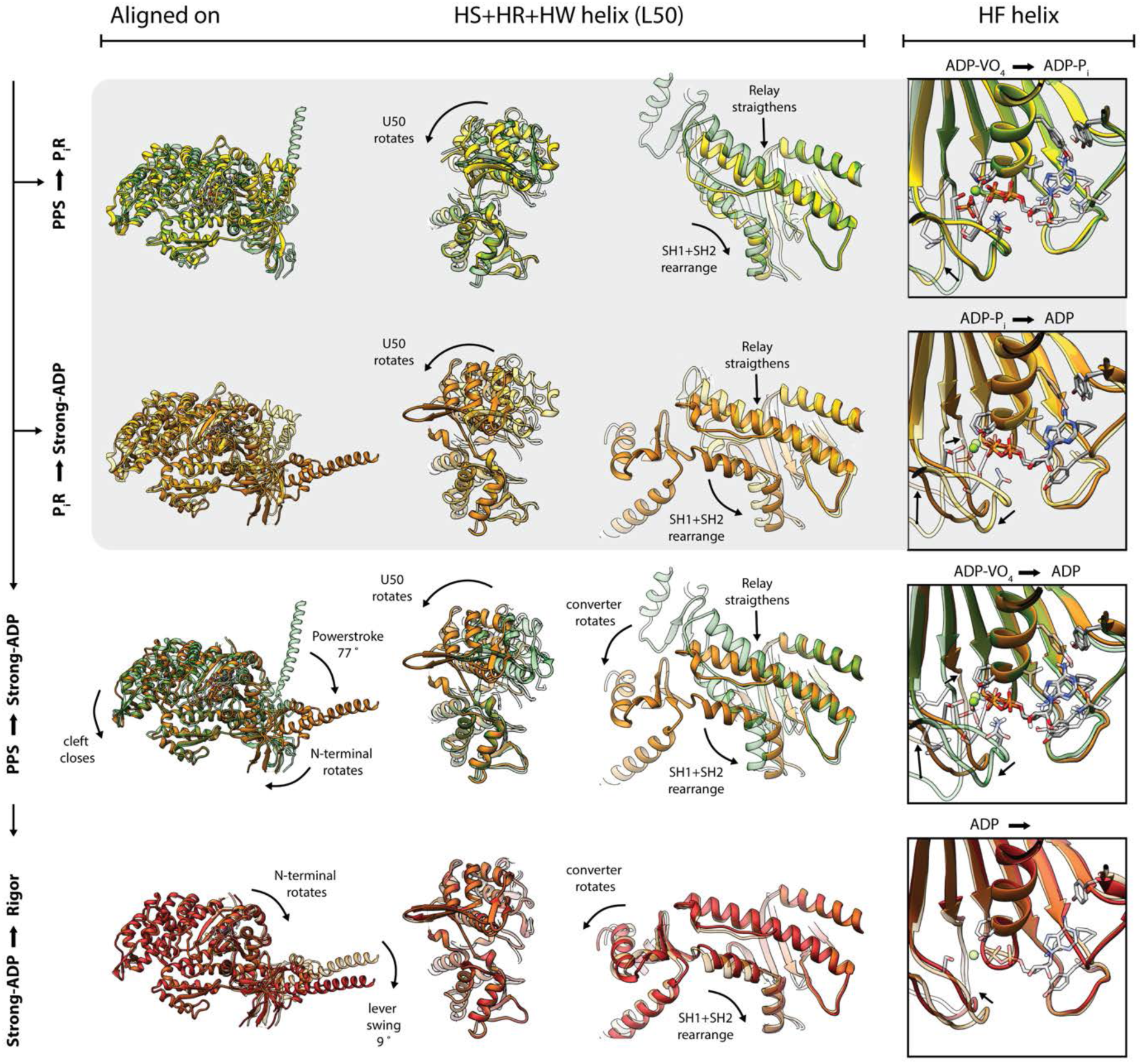
Structural transition of myosin-V along its motor cycle (PPS to Rigor state). Continuation of the superposition of atomic models of successive structural states along the myosin motor cycle shown in Figure 11—figure supplement 3. The lever arm of the PPS state was computationally elongated. Transitions involving the P_i_R state homology model are grayed out to illustrate its limitations. **Video 1. Structural transition of myosin-V along its motor cycle.** Animation of the motor cycle of myosin-V highlighting the allosteric coupling of the active site, the actin binding cleft and the lever arm. Transitions between successive structural states are illustrated by morphs of the corresponding atomic models, also see Figure 11. In addition to the overall conformation, close-up views of the active site, the actin-binding interface and the SH1-SH2-relay helix interface are shown for every transition. A pictogram of the motor cycle illustrates the state currently displayed as well as the rainbow color-scheme used for the depiction of myosin states (rigor: red, PRT: purple, post-rigor: slate blue, intermediate: light blue, PPS: green, PiR: yellow, strong-ADP: orange). F-actin is shown in gray and nucleotides as orange surfaces when showing their binding or release. Major conformational changes are indicated by black arrows and descriptive labels. Note that binding of ATP, release of ADP as well as binding to and detachment from F-actin are only illustrated schematically, intentionally leaving out all molecular details and intermediate states identified to date. Phosphate release and the associated structural changes at the active site are visualized based on a structure-based homology model of the proposed P_i_R state (Llinas et al., 2015). Beyond that, the overall structures of the P_i_R and intermediate state homology models deviate markedly from the ones of other states, in line with the intrinsic structural differences of myosin-VI and myosin-V. Consequently, they cannot provide any further insight into the structural transition of myosin-V and were thus omitted in the animation. Atomic models were either aligned on the HR, HS and HW helices (overall conformation) or on the HF helix (active site). For superpositions of consecutive states, also including the P_i_R and intermediate states, and an illustration of myosin domains and key structural elements see Figure 11—figure supplements 2-4.

